# Multi-omic profiling of tyrosine kinase inhibitor-resistant K562 cells suggests metabolic reprogramming to promote cell survival

**DOI:** 10.1101/308528

**Authors:** Brett M. Noel, Steven B. Ouellette, Laura Marholz, Deborah Dickey, Connor Navis, Tzu-Yi Yang, Vinh Nguyen, Sarah J. Parker, David Bernlohr, Zohar Sachs, Laurie L. Parker

## Abstract

Resistance to chemotherapy can occur through a wide variety of mechanisms. Resistance to tyrosine kinase inhibitors (TKIs) often arises from kinase mutations-however, “off-target” resistance occurs but is poorly understood. Previously, we established cell line resistance models for three TKIs used in chronic myeloid leukemia treatment, and found that resistance was not attributed entirely to failure of kinase inhibition. Here, we performed global, integrated proteomic and transcriptomic profiling of these cell lines to describe mechanisms of resistance at the protein and gene expression level. We used whole transcriptome sequencing and SWATH-based data-independent acquisition mass spectrometry (DIA-MS), which does not require isotopic labels and provides quantitative measurements of proteins in a comprehensive, unbiased fashion. The proteomic and transcriptional data were correlated to generate an integrated understanding of the gene expression and protein alterations associated with TKI resistance. We defined mechanisms of resistance and two novel markers, CA1 and alpha-synuclein, that were common to all TKIs tested. Resistance to all of the TKIs was associated with oxidative stress responses, hypoxia signatures, and apparent metabolic reprogramming of the cells. Metabolite profiling and glucose-dependence experiments showed that resistant cells had routed their metabolism through glycolysis (particularly through the pentose phosphate pathway) and exhibited disruptions in mitochondrial metabolism. These experiments are the first to report a global, integrated proteomic, transcriptomic and metabolic analysis of TKI resistance. These data suggest that although the mechanisms are complex, targeting metabolic pathways along with TKI treatment may overcome pan-TKI resistance.

**Key Points:**

- Alterations to metabolism are a common feature of target-mutation-independent resistance in CML cells across multiple clinically relevant TKIs.
- Carbonic anhydrase 1 (CA1) and a-synuclein (SNCA) are novel markers of metabolic reprogramming in TKI resistant CML cells.

## Introduction

Chronic myelogenous leukemia (CML) is characterized by translocation of chromosomes 9 and 22 to form the Philadelphia chromosome, which generates a fusion between the breakpoint cluster region (*BCR*) gene and the *ABL1* gene. The product of this fusion is the Bcr-Abl protein, in which several of the autoregulatory features of the Abl protein tyrosine kinase are disrupted, leading to its constitutive activity. Tyrosine kinase inhibitors (TKIs) inhibit Abl (and other kinase) activity and are the major treatment modality for CML. The first blockbuster TKI, imatinib, was introduced in the 1990s and provided a transformational improvement in outcomes for CML patients, increasing the five year survival rate from ∼45% to >80% and launching a new paradigm for molecularly targeted cancer therapy that has resulted in development of additional inhibitors for second, third, and further lines of therapy in CML and other cancers. ^(2)^

However, and perhaps inevitably, resistance or failure to respond has emerged as a significant clinical problem, overall affecting about 30% of CML patients and leading to disease progression. ^(3-4)^ Increasing clinical evidence is accumulating that sequential treatment with first, then second, then third line kinase inhibitors (starting with imatinib) does not result in better survival, and in fact, increases the risk of multidrug resistance. ^(5)^ Suboptimal response to imatinib is associated with lack of Bcr-Abl inhibition by 1 month, ^(6)^ and is observed at 18 months in up to 40% of CML patients. ^(3)^ Second line dasatinib and/or nilotinib is effective for about half of imatinib-resistant patients, but third line TKIs do little to improve the long term outlook: patients who fail to respond to two TKIs are unlikely to achieve durable responses with a third TKI. ^(7-8)^ *ABL* mutation (e.g. T315I in *BCR-ABL*) is a clinically significant mechanism of early first line response failure, with incidence between 40-90% (depending on definitions and detection methods), ^(3)^ however second line failure is mutation-independent in 40-70% of cases. ^(7-8)^

In general, mutation-independent resistance can arise from several potential mechanisms, including amplification of the target protein (through copy number increases, increased transcription or translation, and/or decreased turnover), increased drug efflux or decreased drug influx through transporter proteins, or alteration in dependence on other signaling pathways (often termed “kinome reprogramming” ^(9-10)^). Increasing evidence is also accumulating that metabolic reprogramming (relying more heavily on glycolysis as a response to oxidative stress) can enable cancer cells to adapt to stress (including chemotherapeutic stress) and survive in the presence of apoptotic signaling. ^(11)^ Despite a breadth of literature on the role of metabolic reprogramming in solid tumors and other leukemias and lymphomas, ^(11-14)^ there has been less exploration of the potential for metabolic reprogramming to influence sensitivity of CML cells to kinase inhibitors. Evidence for imatinib resistance dependent on metabolic reprogramming through HIF1α has been reported, ^(15)^ however no global evaluation of markers or comparisons to resistance to other TKIs has been performed so far.

Because CML patients that fail two TKIs face dismal outcomes, it is imperative to find new avenues for therapy. Understanding the cooperating molecular pathways that allow CML cells to survive TKI treatment could be instrumental in guiding therapy for these patients. Several studies have examined mechanisms of TKI resistance in CML, typically for one or at most two TKIs, using either RNA sequencing (RNAseq) or protein analyses. However, without complex algorithmic normalization, there is often surprisingly little correlation between transcript and protein levels in the cell. ^(16)^ While gene expression data is highly informative, the absence of protein-level data in TKI resistance severely limits our ability to understand the role of kinome and metabolic reprogramming. To our knowledge, combined comparisons of RNA and protein resistance profiles for all three of commonly used TKIs have not previously been performed. In this report, we sought to define these relationships to obtain a comprehensive characterization of resistance common to multiple TKIs.

We performed transcriptomic and proteomic analyses of the TKI-sensitive human CML cell line K562 and three TKI-resistant derivatives developed in our laboratory: K562-IR (imatinib resistant), K562-NR (nilotinib resistant), and K562-DR (dasatinib resistant). ^(17)^ These cell lines were generated by continuous, increasing dosage exposure to kinase inhibitors over >90 days until a resistant population was generated. Quantitative, whole transcriptome RNAseq and data-independent acquisition (DIA) SWATH-MS datasets were generated. SWATH-MS combines the best features of both narrowly targeted (e.g. multiple reaction monitoring, MRM) and broad, unbiased protein profiling mass spectrometry methods: a significantly greater proportion of proteins are reliably quantified with this method, in comparison to traditional MS methods. This enabled label-free quantitative analysis of relative protein abundance on a global proteome level. Each dataset was analyzed to identify differential gene and protein abundance between cells lines. An integrated analysis of the transcriptomic and proteomic data revealed common features of resistance to these kinase inhibitors: downregulation of Myc targets, engagement of hypoxia-related signaling, alteration of post-transcriptional/translational regulation, and metabolic reprogramming. Labeled glucose feeding experiments confirmed differences in metabolism for the resistant vs. sensitive cells, including increased glycolysis and shunting to the pentose phosphate pathway. Phenotypic experiments testing the ability of the cells to grow in galactose vs. glucose provided evidence that the resistant cells were less able to grow with galactose as an energy source than the sensitive cells, which were better able to adapt likely by routing ATP production through OXPHOS. Overall, these data support the concept that targeting metabolic adaptation may provide options for avoiding and/or overcoming target mutation-independent resistance in CML.

## Materials and Methods

### Cell culture

K562 cells were purchased from ATCC. Imatinib resistant (IR), nilotinib resistant (NR), and dasatinib resistant (DR) K562 cells were generated in the lab as described in our previous report of these cells ^(17)^ by culturing K562 cells in the presence of low, but increasing concentrations of TKIs for 90 days, followed by a constant concentration from then on (1µM imatinib or 10nM nilotinib or 1nM dasatinib supplemented IMDM with 10% fetal bovine serum and 1% penicillium/streptomycin). Cells were grown to 7.5 x 10^5^ cells/mL and split into two equal parts for RNAseq analysis and SWATH-MS.

### RNA sequencing sample preparation and analysis

Cellular RNA was extracted in triplicate for each sample with the RNEasy kit (Qiagen) following manufacturer’s instructions. RNA concentration was measured, diluted to 100ng/µL, and submitted to the University of Minnesota Genomics Center (UMGC) for paired-end RNA sequencing. Twelve barcoded TruSeq RNA v2 libraries (four cell lines, triplicate RNA samples) were created from the samples and combined for sequencing on a HiSeq 2500 using rapid mode with 50bp reads, for a total of ∼5-7 million reads per sample. Data format was converted to FASTQsanger and mapped to human reference genome (hg19_canonical) using TopHat2 on the Galaxy platform installed on the high performance cluster in the Minnesota Supercomputing Institute (MSI) at the University of Minnesota. Differential expression testing between the wild type K562 cells and K562-IR, K562-NR, and K562-DR cells, respectively, was computed as log_2_-fold change (log_2_FC) using Cuffdiff via comparison of fragments per kilobase million (FKPM) values (the standard normalized unit of reads in RNAseq) generated using CuffQuant, a transcript computation program available as a package through Cufflinks. ^(18)^ Subsequently, fusion transcripts were detected using DeFuse, ^(19)^ a software package used to detect fusion transcripts from paired-end RNA-seq data.

### Proteomics sample preparation and mass spectrometry analysis

Cells were washed three times with ice-cold PBS and re-suspended in lysis buffer (50mM ammonium bicarbonate pH 7.0, 4mM EDTA, Roche phosphatase inhibitors cocktail) and immediately incubated at 95°C for five minutes to seize all enzymatic activities. Subsequently, cells were sonicated for 30 minutes in a water bath and the insoluble fraction was separated from the cell lysate by centrifugation at 25,200 RCF for 20 minutes at 4°C. Each sample was reduced (20mM dithiothreitol for 1 hour at 60°C), and alkylated (40mM iodoacetamide for 30 minutes at room temperature in dark.) Reactions were quenched by adding dithiothreitol to a final concentration of 10mM. The samples were then trypsin digested overnight at 37°C at a ratio of 1:50 (w/w). Subsequently, samples were cleaned up using MCX-type stage tips ^(20)^ and re-suspended in 90% water/10% acetonitrile/0.1% formic acid to a final concentration of 0.16 mg/mL. Samples (800ng tryptic digest) were separated on a LC/MS system which included an Eksigent NanoLC 400 system and an AB SCIEX 5600 TripleTOF mass spectrometer. Samples were analyzed using a “trap and elute” configuration on the Eksigent nanoFlex system. Samples were loaded at 2 µL/min for 10 minutes onto a trap column (200 µm x 0.5 mm ChromXP C18-CL chip column) and resolved on an analytical column (75 µm x 15 cm ChromXP C18-CL 3µm). Mobile phase of the liquid chromatography systems are: 0.1% (v/v) formic acid in LCMS grade water (solvent A) and 0.1% (v/v) formic acid in LCMS grade acetonitrile (solvent B). The LC method was over 90 min with a gradient from 5% to 35% solvent B at a flow rate of 300 nL/min. The mass spectrometer was set to acquire SWATH data (i.e. data independent acquisition, DIA). ^(21)^ In a cycle time of about 1.8 sec, one survey scan and thirty-four 26 Da SWATH scans were performed. These 26 Da-wide scan events sequentially cover the mass range of 400 – 1250 Da, with a 1 Da for the window overlap. The collision energy for each SWATH scan was increased sequentially to better fragment peptide ions. Data have been deposited in the SWATHAtlas at http://www.peptideatlas.org/PASS/PASS01196.

### Ingenuity Pathways Analysis

Canonical pathways were associated with both the RNAseq and SWATH-MS datasets using Ingenuity Pathways Analysis. ^(22)^ Canonical pathways analysis identified the pathways from the Ingenuity Pathways Analysis library of canonical pathways that were most significant to the respective data sets. Molecules from the data set that met the log_2_-fold change cutoffs of >1 or <-1, and that were associated with a canonical pathway in Ingenuity’s Knowledge Base, were considered for the analysis. The significance of the association between the data set and the canonical pathway was measured in two ways: 1) A ratio of the number of molecules from the data set that map to the pathway divided by the total number of molecules that map to the canonical pathway is displayed as a Z-score for pathways with sufficient numbers of molecules, and/or 2) Fisher’s exact test was used to calculate an enrichment p-value determining the probability that the association between the genes in the dataset and the canonical pathway is explained by chance alone.

### Metabolite profiling

#### Sample preparation

A total of 21 million of both the wild type (WT) and imatinib-resistant (IR) K562 cells were diluted to 800,000 cells per mL in glucose-free RPMI 1640 medium (Thermo Fisher Scientific) with 5% fetal bovine serum and 1% penicillin-streptomycin, which was supplemented with 25 mM 6-^13^C labeled D-glucose (Cambridge Isotope Laboratories). The cells were subjected to this treatment for 12.5 h, and then the metabolites were extracted in triplicate (7 million cells per replicate) from both the WT and IR lines according to instructions and with materials provided by Human Metabolome Technologies, Inc. (HMT). Briefly, the cells were first centrifuged down at 1,200 rpm for 5 min at room temperature and the culture medium was removed by aspiration. 10 mL of a 5% (w/w) solution of D-mannitol in Milli-Q water was added to each sample and the pellets were resuspended and centrifuged as before. The supernatant was removed by aspiration and then 800 µL of LC-MS grade methanol was added to each cell pellet and vortexed for 30 seconds. Then, 550 µL of an internal standard solution (provided by HMT and diluted 1:1000 in Milli-Q water) was added to each sample and vortexed for 30 seconds. A total of 1000 µL of this mixture was transferred to microtubes which were centrifuged at 2,300 g for 5 min at 4° C. Next, 350 µL of the supernatant was transferred to pre-washed centrifugal filter units in duplicate (due to volume constraints of the filter units) and centrifuged at 9,100 g for 3 h at 4° C. The filtered solutions were then evaporated to dryness using a SpeedVac. The samples were then stored at −80° C until they were shipped to HMT for analysis. At HMT, samples were resuspended in 50 μl of ultrapure water immediately before the CE-TOFMS measurement.

#### CE-TOFMS analysis

Metabolite profiling measurements (HMT’s “F-Scope” service) were performed on each of the 6 samples (3 WT and 3 IR) using capillary electrophoresis time-of-flight mass spectrometry (CE-TOFMS). The compounds were separated on a fused silica capillary column (50 μm x 80 cm) and measured on a CE-TOFMS system (Agilent Technologies) in the anion mode of CE-TOFMS based metabolome analysis. ^(23-25)^ The run buffer and rinse buffer were the HMT Anion Buffer Solution (p/n: H3302-1021). The sheath liquid was HMT Sheath Liquid (p/n: H3301-1020). Samples were injected by pressure injection at 50 mbar over 25 sec. The CE voltage was held constant at positive 30 kV. The MS capillary voltage was 3,500 V, and m/z measurements were taken in ESI negative mode with a scan range of m/z 50-1,000.

#### CE-TOFMS Data Processing and Analysis (performed by HMT)

Peaks detected in CE-TOFMS analysis were extracted using automatic integration software (MasterHands ver. 2.17.1.11 developed at Keio University) in order to obtain peak information including m/z, migration time (MT), and peak area. The peak area was then converted to relative peak area by the following equation†2. The peak detection limit was determined based on signal-noise ratio; S/N = 3.

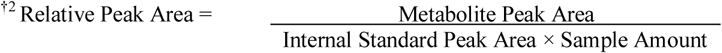

Putative metabolites were then assigned from HMT’s target library and their isotopic ions on the basis of *m/z* and MT. The tolerance was ±0.5 min in MT and ±30 ppm^†3^ in *m/z*.

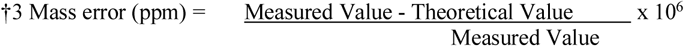

Absolute quantification was performed in total amount of each detected metabolite. All the metabolite concentrations were calculated by normalizing the peak area of each metabolite with respect to the area of the internal standard and by using standard curves, which were obtained by single-point (100 µM) calibrations. Data results are available in the Supplementary Information (“Metabolite Profiling.zip”).

### Oxygen consumption rate experiments

Oxygen consumption was analyzed in duplicate using the XF24 Analyzer and system software (Seahorse Bioscience, Billerica, MA). The cell lines were grown as described above. On the day of the assay, approximately 200,000 cells were seeded per well in XF24 V7 culture dishes precoated with 0.2% polylysine (Sigma). The cells were allowed to adhere to the plate for 1-2 hrs in a 5% CO_2_ 37°C incubator. The cells were then switched to serum-free modified, no-bicarbonate, low-phosphate DMEM (D5030) supplemented with 1× glutamax (Invitrogen), 1 mM sodium pyruvate, 25 mM glucose, and allowed to equilibrate for 1 h in a non-CO_2_ 37°C incubator. After cartridge calibration, cells were placed in the XF24 analyzer where cellular oxygen consumption rate was measured three times under basal conditions and then three times after the addition of 1 µM oligomycin, three times after FCCP addition (0.25 – 2uM), and twice after 1 µM antimycin A. After each experiment, protein content per well was measured using the BCA assay to normalize number of cells analyzed per well. Normalized oxygen consumption rate (pmol/min/mg protein) data obtained from the Seahorse XF24 Software were plotted using GraphPad Prism. Brown-Forsythe and Welch ANOVA tests with Holm-Sidak’s multiple comparisons tests were performed to evaluate significance of differences between the control and TKI resistant cell lines.

## Results

### Whole transcriptome analysis: Resistance is mediated by pathway alterations and not *BCR-ABL1* gatekeeper mutations

In order to detect differences in gene expression associated with TKI resistance, we performed whole transcriptome RNA sequencing analysis on parental K562 human chronic myeloid leukemia cells and three drug-resistant derivatives, K562-IR (imatinib-resistant), K562-NR (nilotinib-resistant), and K562-DR (dasatinib-resistant). Sequencing was performed for three replicate samples from each cell line. Fusion transcripts were detected using the DeFuse package ^(19)^ in Galaxy. The *BCR-ABL1* t(9;22) fusion transcript was validated in each cell line, and several other fusions were also observed (including e.g. the known fusion *NUP214-XKR3* t(9;22) ^(26-27)^) (Supplementary Table S1). To examine the *BCR-ABL1* transcripts for potential drug-resistant point mutations, a custom version of the human hg19 genome was built to incorporate the *BCR-ABL1* fusion gene, map the specific fusion transcripts and identify whether point mutations in the gatekeeper residue were associated with inhibitor resistance. Using IGV Browser (Broad Institute) to view the mapped reads of each TKI-resistant derivative against this custom genome, we did not identify any point mutations that were significantly different in the resistant vs. the sensitive cell lines. In particular, the gatekeeper residue T315 was not modified, strongly suggesting that gatekeeper mutations were not contributing to drug resistance in these cell line models (Supporting information Fig S1).

We compared the differentially expressed genes of each TKI resistant cell line relative to the parental, sensitive cell line (Supplementary Tables S2-S5). Each TKI resistant cell line differentially expressed a unique set of genes (227 for the imatinib-resistant cells, 327 for the dasatinib-resistant cells, and 1930 for the nilotinib-resistant cells). We found 370 genes that were differentially expressed in common across all three TKI resistant cell lines (Fig. 2A). Of these, 117 were downregulated and 253 were upregulated by log_2_ fold-change of at least at least −1 or 1, respectively in each TKI resistant sample, with 97% concordance of log_2_ fold-change direction per transcript across all three cell lines (Table S7). Overall, 842 genes were differentially expressed in at least one of the TKI resistant cell lines by log_2_FC >1 or <-1 (Figure 2B). Generally, the IR and DR cell lines showed more similarities in differential gene expression than with the NR cell line, both in terms of number of shared differential gene identifications (Figure 2A: 243 for IR:DR, compared to 212 for IR:NR and 117 for NR:DR) and as determined through clustering of the cell lines based on the 842 genes differentially expressed in at least one cell line (using the HeatMapper tool, ^(1)^ columns clustered by Euclidean distances using complete linkage). However, because we were most interested in the mechanisms that might be common targets in TKI resistance, we focused on genes that were commonly differentially expressed in all the drug resistant cell lines. Several of these transcripts have previously been identified as associated with TKI resistance, including MDR1/ABCB1, CD36, CD44, β-catenin, FYN, and AXL (Fig. S2, S5), indicating that our models were comparable to others developed in the literature. ^(28-32)^ Ingenuity Pathway Analysis (IPA) performed on the differential expression data for the 370 common genes (Figure 2C) identified oxidative stress-related and inflammatory processes as altered, e.g. NFκB signaling, as well as decreased signaling by PTEN (a tumor suppressor pathway) and PPAR (a metabolic signaling pathway). Gene Set Enrichment Analysis (GSEA) (Fig. S3 and S4) also identified alterations to Myc programming and activation of HIF1α pathways in the TKI resistant cells. These represent novel pathways associated with TKI resistance. Interestingly, all of these alterations involve metabolic changes. ^(33), (34)^

**Figure 1.**
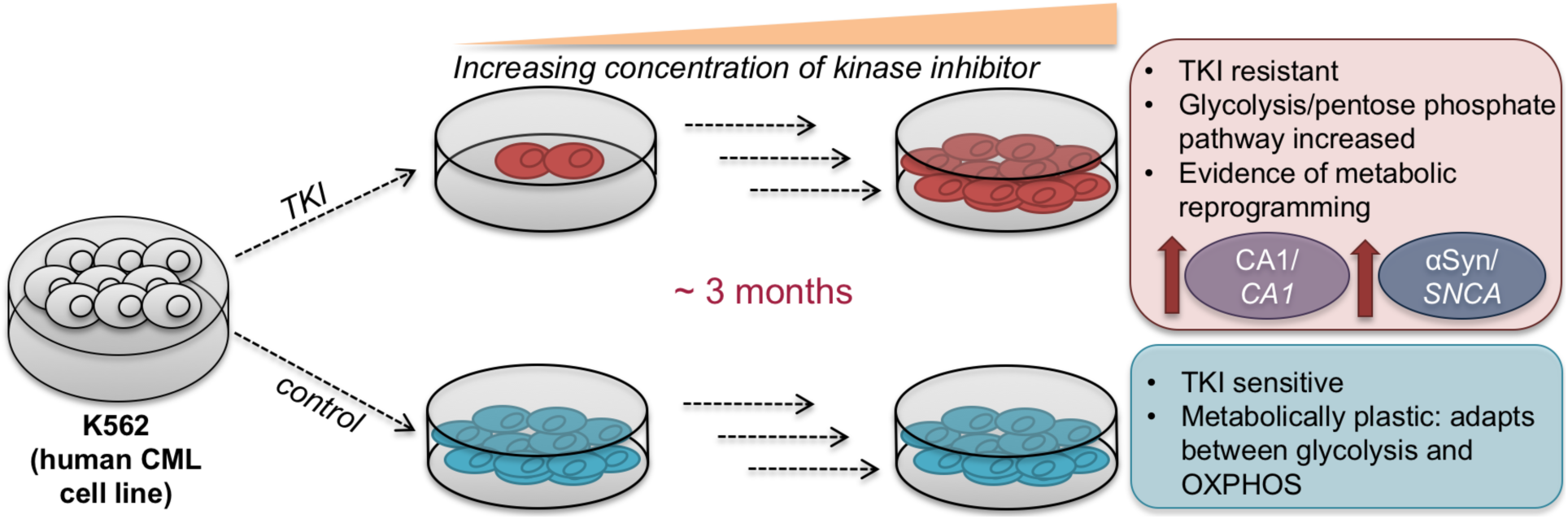
Developed resistance of a CML cell line to three different clinically-relevant tyrosine kinase inhibitors (TKIs) is associated with metabolic reprogramming that alters glycolysis and metabolic plasticity. Carbonic anhydrase 1 (CA1/*CA1*) and α-synuclein (αSyn/*SNCA*) protein and mRNA levels emerged as a potential functional biomarker signature for this type of metabolic reprogramming in K562 CML cells.

**Figure 2.**
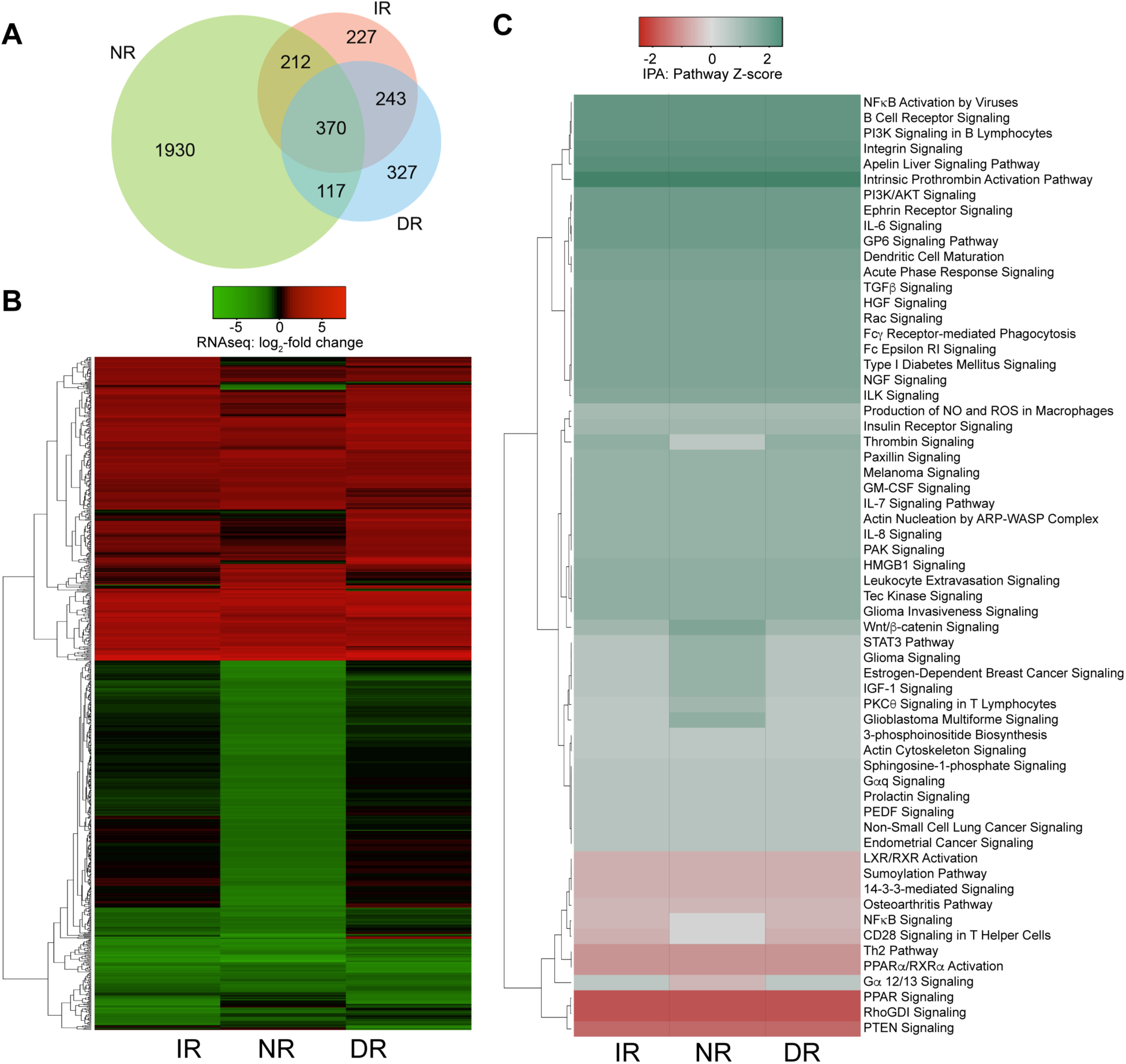
RNAseq differential gene expression and pathway analysis summary. A) Venn diagram showing number of overlapping and unique genes differentially expressed (calculated as log_2_ fold change) observed in RNAseq data from the different TKI resistant cell lines relative to the control K562 cells. IR = imatinib-resistant, DR = dasatinib-resistant, NR = nilotinib resistant. B) Heatmap of RNAseq data generated using *HeatMapper*: ^(1)^ log_2_ fold change (log_2_FC) values for 842 genes differentially expressed at log_2_FC >1 or <-1 in at least one of the resistant lines, clustered according to complete linkage using Euclidean distances. Color scheme represents log_2_FC of each gene’s expression for each cell line. C) Heatmap of Ingenuity Pathway Analysis Z-score values for enrichment of pathways represented by the 842 genes shown in 2B.

### Quantitative protein level profiling: Proteome-level pathway alterations are consistent with gene expression-level pathway observations

To define mechanisms of TKI-resistance at the protein level, we also performed quantitative proteomics (on the same set of samples as for RNAseq) using SWATH-MS data on a 5600+ TripleTOF^®^ instrument (SCIEX). Protein extracts were analyzed using a fixed window (25 Da) in SWATH mode, which allowed for recording of fragment ion spectra from all detectable peptides in the samples and enabled unbiased label-free quantitative comparisons of the proteome (Tables S8 and S9). These analyses detected a set of uniquely differentially expressed proteins in each TKI-resistant cell line relative to the parental control (8, 7, and 33 for the imatinib-resistant, dasatinib-resistant and nilotinib-resistant cell lines, respectively). 81 proteins were differentially expressed in common amongst the three drug resistant cell lines compared to the sensitive parental control cells, with 70% concordance of log_2_ fold-change direction (Figure 3A,B). The 81 common proteins were analyzed Ingenuity Pathway Analysis. While the number of molecules available per pathway was not sufficient to calculate Z-scores, Ingenuity Pathway Analysis pathway enrichment p-values (Figure 3C) were able to be calculated and supported the gene expression analyses that oxidative stress and metabolic processes were different (relative to the TKI-sensitive cells) in all three of the TKI resistant cell lines. As discussed below, the protein-level data provided important information about whether transcripts observed in the RNAseq experiment actually gave rise to altered levels of proteins, and illustrate how even with substantial mRNA log_2_ fold-change concordance, protein alterations were more variable between the different cell lines.

**Figure 3.**
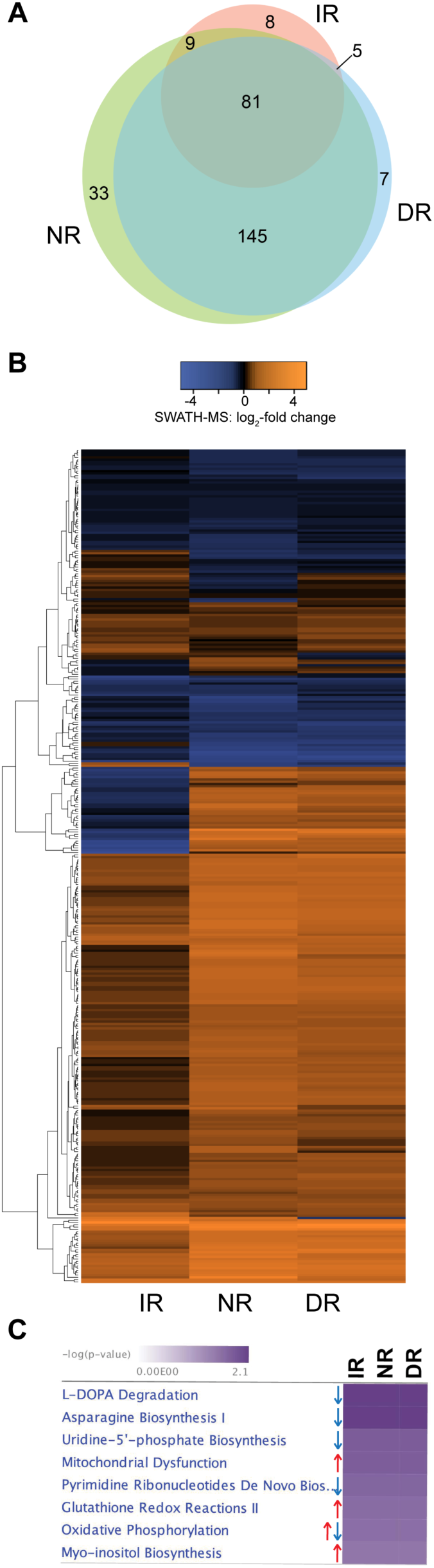
SWATH-MS differential protein and pathway analysis summary. A) Venn diagram showing number of overlapping and unique proteins measured by SWATH-MS (calculated as log_2_ fold change) from the different TKI resistant cell lines relative to the control K562 cells. IR = imatinib-resistant, DR = dasatinib-resistant, NR = nilotinib resistant. B) Heatmap of SWATH-MS data generated using *HeatMapper*: ^(1)^ log_2_ fold change (log_2_FC) values for 365 proteins differentially detected at log_2_FC >1 or <-1 in at least one of the resistant lines, clustered according to complete linkage using Euclidean distances. Color scale represents log_2_FC value. C) Heatmap of Ingenuity Pathway Analysis emrichment p-values for pathways represented by the 365 proteins shown in 3B.

### Combined analysis: Relationships between gene expression and protein level patterns in kinase inhibitor resistance

We performed an analysis to integrated transcriptomic and proteomic datasets to examine the relationships between mRNA and protein alterations in TKI-resistant cells. RNAseq and SWATH-MS datasets were aligned using the OneOmics MultiOmics tool to identify transcript observations that had corresponding protein observations (Supplementary Table S10). 588 were matched in common across all cell lines with sufficient confidence at both the mRNA and protein levels. The relationship between mRNA and protein fold changes was examined for transcripts and their associated proteins that were observed in both. Overall, the correlation (Spearman non-parametric with two-tailed p-value) between mRNA and protein level fold changes in each resistant cell line relative to the control were weak to moderate (Fig. 4A). In general, the magnitude of differential expression for proteins varied more widely than it did for mRNAs between the TKI sensitive vs. the TKI resistant cells: in many cases for a gene, the protein levels exceeded the fold change cutoff (log_2_FC >1 or log_2_FC<-1) but the respective mRNA levels did not (log_2_FC = −1 to 1). In particular, for example, the mRNA levels for the 40S ribosomal subunit RPS29 were increased in all three resistant cell lines, while its protein levels were not. This finding and the overall lack of strong correlation between mRNA and protein levels are consistent with the enriched GO annotations observed for the SWATH-MS results suggesting that post-transcriptional or translational regulation may be an important mechanism in TKI resistance and highlight the importance of matched, protein-level analysis in the study of TKI resistance.

**Figure 4.**
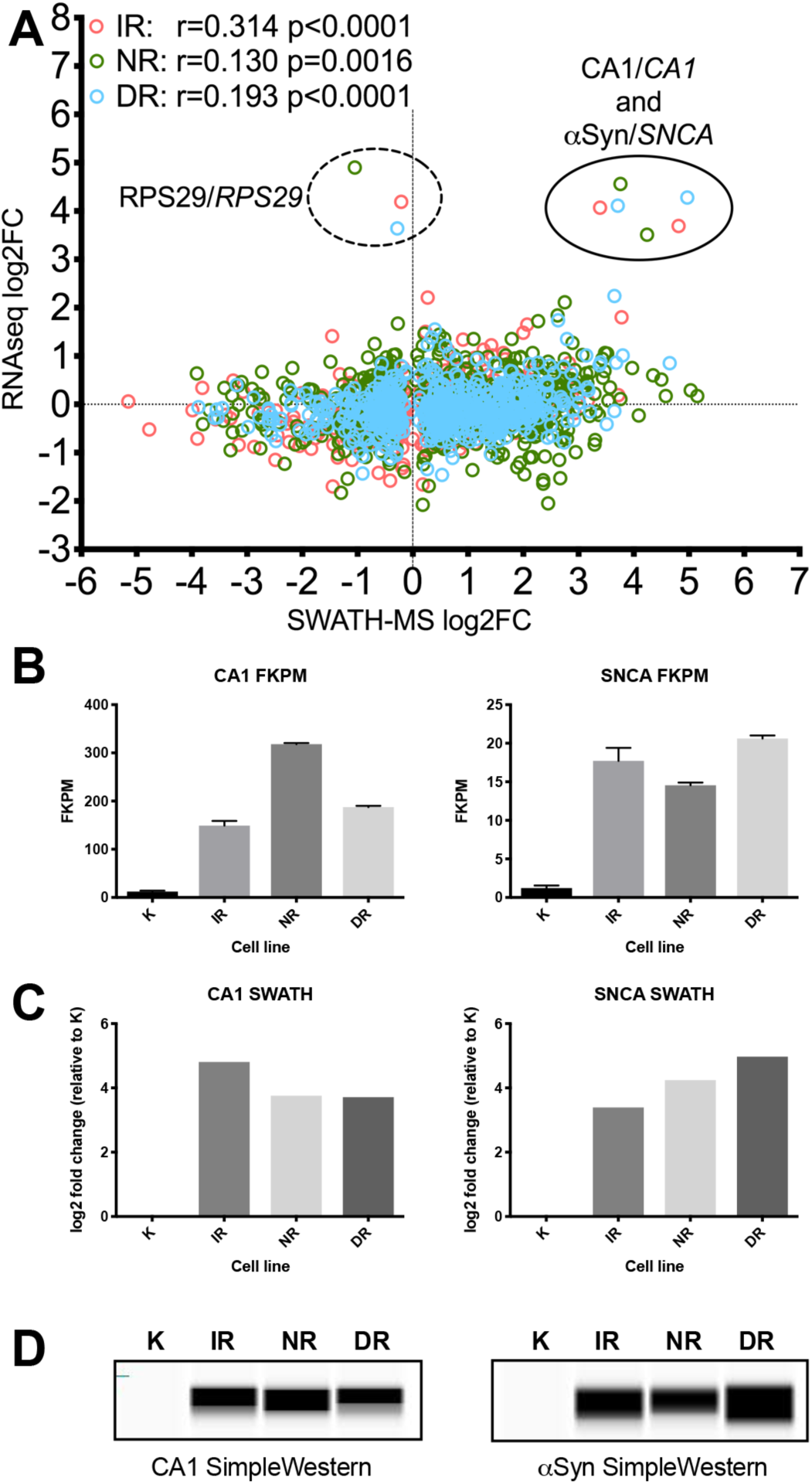
Relationships between mRNA and protein observations. A) Relationship between mRNA and protein log_2_-fold change levels from RNAseq and SWATH-MS. IR = imatinib resistant, NR = nilotinib resistant, DR = dasatinib resistant. Spearman non-parametric correlation analyses were performed in GraphPad Prism. B) FKPM values plotted for selected gene transcripts showing upregulation at both mRNA and protein levels: CA1 and SNCA (-Synuclein). Error bars represent SEM for three replicate sequencing runs. K = control K562 cells, IR, NR and DR as above. C) SWATH-MS quantitation values from the OneOmics workflow plotted as log_2_-fold change for resistant lines relative to K, with unchanged K column shown for reference. D) ProteinSimple SimpleWestern chemiluminescence pseudo-blots showing immunodetection of these three proteins. Full SimpleWestern traces and pseudo-blot images available in the Supporting Information.

We identified two robustly expressed proteins with a large increase in expression at both the mRNA and protein level in all of the resistant samples: carbonic anhydrase 1 (CA1) and α-synuclein (αSyn) (Fig. 4B-C). CA1 is an enzyme that, among other activities, aids in maintaining pH balance both intra- and extracellularly in many systems and participates in H+/lactate co-transport. Increased carbonic anhydrase activity has previously been linked to cells’ adaptation to increased lactate levels in response to high rates of glycolysis, ^(33, 35-36)^ therefore its upregulation in these cell lines may be related to metabolic changes arising from mitochondrial dysfunction suggested by Ingenuity Pathway Analysis (Fig. 3C). A role for αSyn in K562 cells has not been previously described; this protein is primarily studied in the setting of Parkinson’s disease. However, αSyn interacts with the mitochondrial membrane and is required for mitochondrial fission in response to mitochondrial stress, ^(37)^ and plays a role in mitochondrial membrane depolarization ^(38-39)^ suggesting a relationship between this protein and the metabolic alterations detected in the transcriptional analysis. Additionally, αSyn upregulation has been observed in myelodysplastic syndrome and megakaryoblastic leukemias, ^(40)^ and given that myeloid progenitor cells give rise to megakaryocytes ^(41)^ it is possible that αSyn induction in these cell lines may be related to functional changes in the differentiation state of the resistant cells. αSyn is also associated with extracellular vesicle release from platelets ^(42)^ (which arise from the megakaryocyte lineage), and increased extracellular vesicle release has previously been observed in cells with drug resistance associated with metabolic alterations. ^(14)^ However, in the case of our models, exosome extraction from the media followed by nanoparticle counting to quantify numbers of exosomes (further described in the supporting information) showed no significant increase in exosome secretion in the drug resistant cells relative to the control cells (Fig. S6).

### Metabolic reprogramming in TKI resistance: increased dependence on glycolysis and shunting through the pentose phosphate pathway

Our integrated proteomic and transcriptional data suggest that metabolic reprogramming may be an important mechanism of survival in TKI resistant cells. To test this possibility in one of the cell lines as a pilot, we performed a ^13^C-labeled glucose pulse-chase feeding experiment to compare glucose metabolism in IR cells relative to TKI sensitive cells. The cells were passaged into media containing 6-^13^C-labeled glucose and allowed to grow for 12.5 hours. After this treatment, the metabolites were harvested and analyzed to study the differences in relative abundance and degree of ^13^C incorporation (which can suggest degree of glucose usage in a given pathway) for a panel of metabolites. IR cells showed a consistent and significant increase of relative abundance of ^13^C-labeled glycolysis and pentose phosphate pathway (PPP) metabolites in comparison to the parental line (Figure 5A and B).

**Figure 5.**
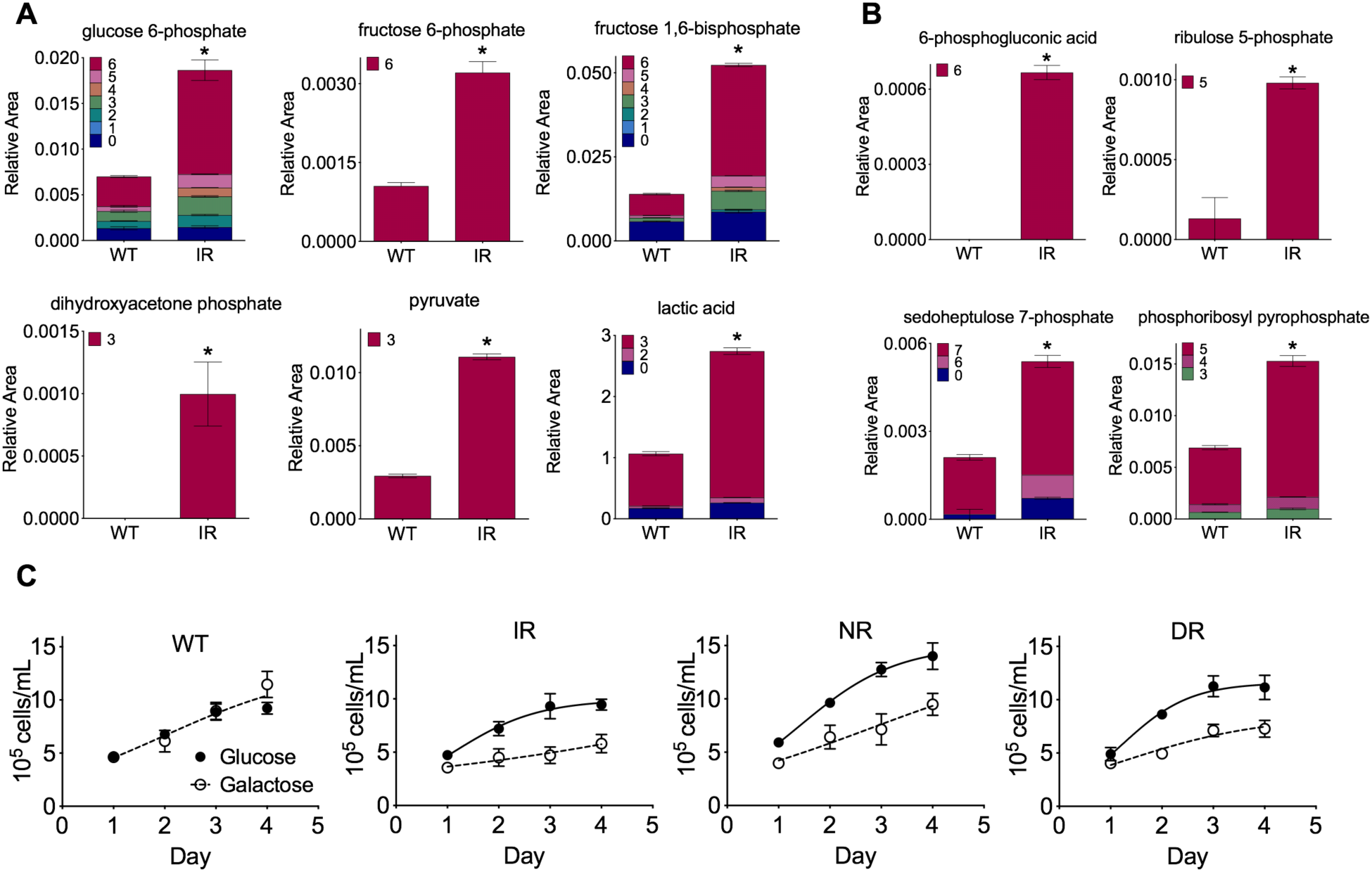
Metabolite profiling, galactose feeding and mitochondrial dysfunction experiments. Metabolite profiling via labeled glucose feeding, comparing “WT” (parental control) with imatinib-resistant (IR). (A) Comparisons for six metabolites involved in the glycolysis pathway of energy production; (B) Comparisons for four metabolites involved in the pentose phosphate pathway (PPP) or, in the case of phosphoribosyl pyrophosphate, synthesized downstream from the PPP intermediate ribulose 5-phosphate. Color schemes represent number of ^13^C-labeled carbons incorporated (i.e. from labeled glucose), red indicates maximum number of carbons in molecule were labeled. Values are expressed as Relative Area (x-axis) compared to internal standard. Error bars represent standard error of the mean for three technical replicates. * = statistically significant difference measured by multiple t-tests with Bonferroni-Sidak correction. (C) Cell growth curves for untreated control (WT), imatinib-resistant (IR), nilotinib-resistant (NR) and dasatinib-resistant (DR) cells. Cells were grown in duplicate in either glucose-containing (solid circles and lines) or galactose-containing (open circles and dashed lines) media for a total of four passages over 16 days. Growth curves for each set of four days after a passage from both duplicates were averaged, and error bars represent standard error of the mean (SEM). Differences between the curves were calculated by measured by extra sum of squares F-test in GraphPad Prism.

As for many cancers, K562 cells exhibit increased glycolytic activity (relative to non-leukemia cells), commonly known as the Warburg effect. ^(43-44)^ Prior evidence from the literature suggests that this is initially suppressed by imatinib exposure (creating the “reverse Warburg effect”, in which metabolism in cancer cells shifts away from glycolysis). ^(43-44)^ Those studies showed that glucose uptake and lactate production (by glycolysis) were suppressed and mitochondrial oxidative phosphorylation through the citric acid cycle was enhanced by imatinib treatment. In other words, imatinib treatment of TKI-sensitive K562 cells leads to a shift of metabolism towards that of a non-malignant cell. Our data reveal that over the longer-term IR cells likely reverse this effect: they display significantly elevated levels of metabolites generated by glycolysis (Figure 4A) and the PPP (Figure 5B). These data suggest that this imatinib resistance is associated with a reversion to increased glycolysis, even using glycolysis to a higher degree than observed in untreated K562 cells. Citric acid cycle intermediates, such as α-ketoglutarate, were either not changed or slightly decreased (raw and plotted data available in supplemental file “Metabolite Profiling.zip”), suggesting that glycolysis was affected more than TCA metabolism. Furthermore, the proportion of G6P containing all six ^13^C-labeled carbons was higher in the IR than in the control cells (Fig. 5A, red portion of bars), indicating that the resistant cells more readily take up glucose—meaning they must have either reversed or not exhibited the suppression of glucose uptake which may have initially occurred from imatinib exposure early in their development that others have described previously. ^(43-44)^

The significantly increased relative abundance and nearly comprehensive ^13^C-labeling (as indicated by red portion of bars) of metabolites in the PPP in IR cells (Figure 5B) suggests that the resistant cells processed the labeled glucose rapidly through that pathway as well. Activation of the PPP has previously been identified in the cellular redox response, ^(45)^ and intriguingly, has also been observed as a key feature of multidrug resistance in acute myeloid leukemia (including resistance to a kinase inhibitor). ^(13)^ Separately, it also has been associated with decreased pyruvate kinase (PKM) activity ^(46)^ causing accumulation of phosphoenolpyruvate (PEP), which in turn inhibits triosephosphate isomerase (TPI). TPI is a glycolytic enzyme that interconverts dihydroxyacetone phosphate (DHAP) to glyceraldehyde-3-phosphate, and its inhibition would lead to accumulation of DHAP and absence of glyceraldehyde-3-phosphate. While we did not detect increased PEP levels in IR cells, we did see a marked increase in DHAP levels (from none detected in control cells to a clearly detectable level in the IR cells, Figure 5A) and did not detect any glyceraldehyde-3-phosphate (in either cell type), consistent with TPI inhibition. Therefore, these ^13^C-labeling experiments also suggest increased PPP activity in IR cells. Together, these data confirm our transcriptomic and proteomic findings and demonstrate that TKI-resistance is associated with metabolic reprogramming towards glycolytic metabolism and a shift into the pentose phosphate pathway. Interestingly, these data show that TKI resistance is associated with the reversion of the metabolic changes that have been previously associated with TKI treatment in sensitive cells. The shifts in metabolism we observed have been associated with a variety of malignant phenotypes in cancer cells, including drug resistance. ^(47)^

Next, we tested whether all three of the TKI-resistant cell lines were using glycolysis as their major source of energy for growth. This served as a phenotypic confirmation of the apparent metabolic changes indicated by the ^13^C-labeled glucose feeding experiment in the IR cells. Cells that are grown in high glucose are known to adaptively suppress oxidative phosphorylation and use glycolysis through a phenomenon known as the Crabtree effect, ^(48)^ therefore increased glycolytic activity (such as that we observed via metabolite profiling) could have just resulted from the high glucose culture conditions. However under conditions of lower glucose availability, cells that are competent for oxidative phosphorylation adapt to instead generate ATP through oxidative phosphorylation via mitochondrial respiration whereas cells that cannot compensate with oxidative phosphorylation exhibit growth defects. ^(49-50)^ This metabolic plasticity can be used to test glycolysis dependence by using galactose as an energy source in place of glucose. Glycolysis is inefficient with galactose because one equivalent of ATP is required to metabolize galactose first via conversion to a UDP-galactose intermediate and finally to glucose, ^(51)^ upon which the glucose is further processed to generate two equivalents of ATP, resulting in net energy production of just one ATP equivalent. When switched to galactose-containing medium, cells that can compensate with oxidative phosphorylation are able to route the energy derived from the galactose through mitochondrial metabolism and the citric acid cycle (which generates much higher-fold ATP per molecule of sugar) grow as well as or nearly as well as those in glucose. In contrast, cells with defects in galactose metabolism or intrinsic downregulation of oxidative phosphorylation grow slowly or not at all. ^(49-50)^ Pathway analysis of the protein expression data had detected downregulation of pathways that are required for metabolism of galactose to glucose, including proteins responsible for uridine-5’-phosphate biosynthesis and pyrimidine ribonucleotide de novo biosynthesis (Figure 3C). This suggested that the TKI resistant cells would be less competent for galactose metabolism. To test this, we compared the growth of IR, NR and DR cells in glucose- or galactose-containing medium relative to the control cells. The control cells grew equally well in both media. In contrast, the TKI resistance cells exhibited a marked, significant growth inhibition in galactose (p <0.0001 by Fisher’s exact test comparison of growth curves) and did not reach the same population levels (Figure 5C), indicating that the TKI resistant cell lines are defective in obtaining sufficient energy through oxidative phosphorylation. ^(49-50)^

Mitochondrial dysfunction was further evaluated in oxygen consumption rate experiments (Fig. 6A). Control or TKI resistant cells were harvested in log phase growth and seeded onto polylysine-coated Seahorse XF24 analyzer plates, and oxygen consumption rates were measured before and after sequential treatment with oligomycin (ATP synthase inhibitor), carbonyl cyanide-4(trifluoroxymethoxy)phenylhydrazone (FCCP) (an oxidative phosphorylation uncoupling agent), and antimycin A (which completely inhibits cellular respiration) (Figure 6A). Basal mitochondrial respiration, ATP turnover, and maximum respiratory capacity were all decreased in the TKI resistant cells relative to the control cells (Figure 6B), which is consistent with some mitochondrial dysfunction in the TKI resistant cell lines. We also observed that twice as many of the proteins observed by SWATH-MS to be altered by log_2_-fold change of >1 or <-1 in the resistant cells (59 of 489 total, 12.1%) are assigned as mitochondrial by the Human MitoCarta ^(52)^ relative to the proportion of mitochondrial proteins in the whole proteome (1158 of 21,118 total, 5.6%), a difference that was statistically significant (p <0.0001) according to Fisher’s exact test. We extracted log_2_-fold change values for these proteins and their corresponding mRNA from our SWATH-MS and RNAseq data sets (as previously described in Figures 2 and 3), and replotted them in a separate heat map (Fig. 6C). Notably, alterations in several key proteins for ATP production and utilization were observed (marked with * in Figure 6C), including decreased ADT2 (which is responsible for transport of ADP into the mitochondria and export of ATP out into the cytoplasm), decreased MPCP (the mitochondrial phosphate carrier protein that imports phosphate into the mitochondria for use in ATP synthesis), and imbalance in the levels of multiple ATP synthase components. Voltage dependent anion channels VDAC1 and 2 (which are key mitochondrial membrane transporters for metabolites) were also decreased. These findings are consistent with disruptions to mitochondrial metabolism. Also VDAC1 may participate in the formation of the permeability transition pore complex that triggers apoptosis, and thus we speculate that its decrease might be related to the survival of the TKI resistant cells—however this would require further confirmation to determine.

In summary, the data here demonstrate the pan-TKI resistance is associated with metabolic reprogramming that shifts glucose metabolism into the pentose phosphate pathway, and appears to also involve a decreased ability to compensate for impaired glycolysis via oxidative phosphorylation that may arise from alterations to mitochondrial protein levels. It is possible that pharmacological interventions that target this effect may be useful for preventing the development of TKI resistance, however, further work would be needed to identify viable targets in the pathways that regulate this process.

## Discussion

This study used transcriptomic, proteomic and metabolite profiling of *in vitro* models to broadly evaluate potential mechanisms of TKI resistance in CML. Overall, the gene expression and protein level data from our study suggest that K562 CML cells adapt at multiple levels to grow in the presence of the three commonly used kinase inhibitor drugs, but that metabolic changes underlie many aspects of these levels. Several markers previously found in clinical and animal model-based studies of TKI resistance were observed suggesting that this model does recapitulate well-validated features of TKI-resistance. These features included MDR1/ABCB1, ^(28)^ the β-catenin/CD44 pathway, ^(30)^ and Axl kinase. ^(32, 53)^ We also observed features of metabolically-related resistance mechanisms, including apparent induction of hypoxia signaling and PI3K signaling (observed as described above by GSEA and Ingenuity Pathway Analysis), enrichment of proteins related to myo-inositol biosynthesis (part of the pentose phosphate pathway), upregulation of CA1 (indicative of increased need for management of lactate in the cell), as well as the fatty acid transporter CD36, a metabolism-related protein that previously had only been found in TKI resistant CML cells residing in an adipose niche *in vivo*, and a fatty acid binding protein FABP5 (family member of FABP4, which was part of the CD36-related mechanism). ^(29)^ The observations of MYC, HIF1 and PI3K pathways is consistent with what is known about tumor metabolic reprogramming and drug resistance, ^(54)^ including in BcrAbl-dependent cells, ^(15)^ providing further support for the likelihood that these cells have undergone changes to their metabolism in order to evade cell death in the presence of these TKIs. ^(11)^ While there are many caveats to using such a model, including the lack of bone marrow niche or other relevant *in vivo* microenvironment, and the differences in dosages present in suspension culture vs. the drug concentration fluctuations and gradients that would be present *in vivo*, this collection of literature-corroborated observations also suggests that despite the limitations of *in vitro* culture of long-established cell lines, these models provide a reasonable degree of relevance to the systems they are intended to mimic. Based on that, the multi-omic data reported in this work should be of value for considering potential alternative therapeutic mechanisms to pursue for TKI resistant CML.

### Metabolic reprogramming as a general drug resistance strategy for K562 CML cells

Multi-drug resistance through metabolic reprogramming and imbalance of the redox equilibrium is increasingly being recognized as an important consideration in chemotherapy for leukemias. ^(55-56)^ While TKIs have been very successful in managing CML for about 70-80% of patients, non-response and/or resistance are still significant problems. Drugs such as ponatinib (a next-generation TKI that can inhibit the T315I mutated BcrAbl) have been developed to address the most common BcrAbl mutations, but these do not solve the problems of metabolic and redox-related multi-drug resistance. Finding ways to combat more general forms of resistance could have broad utility for off-target TKI resistance in CML and other cancers. Our studies identified these types of metabolic reprogramming and redox features associated with resistance to all three commonly used TKIs, imatinib, nilotinib and dasatinib, indicating that they could play a role in multi-drug TKI resistance in CML.

In particular, the upregulation of CA1 and αSyn could be mechanistically relevant markers of metabolic dysregulation in TKI resistance (Figure 7). While this study has not identified specific mechanistic roles for these proteins in metabolic reprogramming, there is literature evidence that these markers may be reporters of the two metabolic alterations observed (increased glycolysis activity and apparent defects in mitochondrial OXPHOS). CA1 is important for buffering cells in the presence of increased lactate concentrations, which result from increased flux through glycolytic pathways. αSyn is involved in mitochondrial dysfunction and handling increased ROS, both of which are related to metabolism and redox balance in the cell. ^(38-39), (57), (58)^ CA1 and αSyn upregulation in these TKI resistant CML cell models could thus be part of a response to metabolic shifts, enabling the cells to better manage the increased lactate and ROS and avoid apoptosis. Although αSyn has been observed in certain cell lines (but not in K562) in a study looking at synucleins in a few hematological malignancies, ^(40)^ to our knowledge it has not previously been linked to metabolic reprogramming or drug resistance in leukemias. Ultimately, these metabolic adaptations seem to be relatively independent of the specific kinases or cancer type. ^(59), (60-61)^ The metabolic adaptations we observed were not consistent with recently proposed models in which increased OXPHOS is a marker of early-stage non-response to chemotherapies (as opposed to apparent deficiency in OXPHOS, as in our case), ^(56, 62-63)^ suggesting that longer time courses of exposure may be needed in order to understand longer term resistance. The detailed mechanisms of how the metabolic alterations enable cell survival in the presence of the three TKIs still need to be elucidated, and studies are ongoing to identify the key metabolic processes (e.g. NADPH production through the PPP ^(13)^, and/or fatty acid metabolism ^(64)^) so they could be tested as targets to prevent resistance or resensitize cells to TKIs.

**Figure 7.**
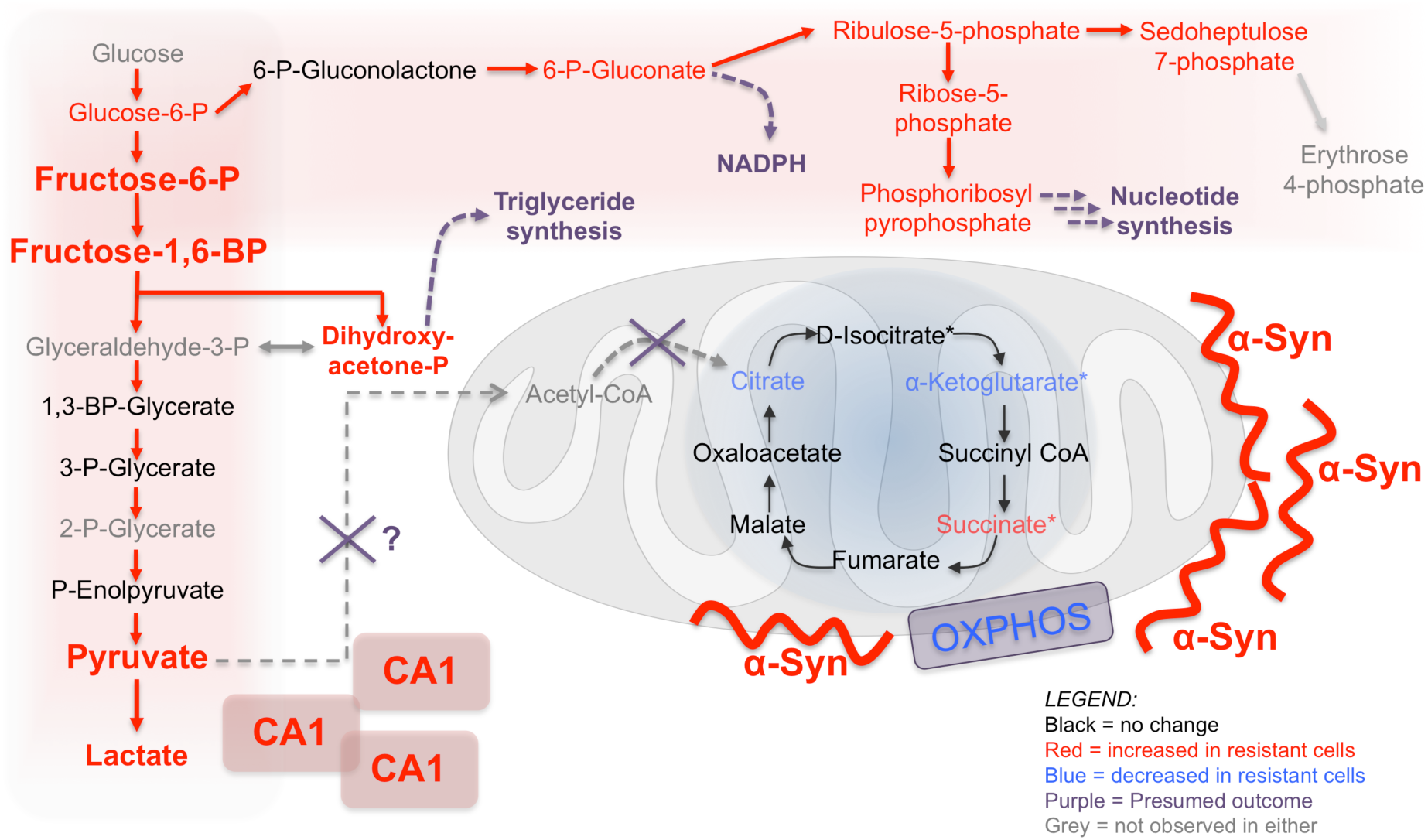
Functional markers of metabolic adaptation in TKI resistant cells. Pan-TKI resistant K562 cells exhibit increased glycolysis and PPP metabolism, with alternate use of dihydroxyacetone-phosphate (presumably re-routed for triglyceride synthesis) and a disconnection of the glycolysis-citric acid cycle cross-talk that is normally mediated through pyruvate. Carbonic anhydrase 1 is upregulated, likely to help buffer excess lactic acid from the increased glycolysis rate, and α-synuclein is upregulated, perhaps to modulate mitochondrial membrane polarization and mitochondrial stress related to the metabolic alterations.

## Supporting information

Exosome supplemental file

SWATH library file

Metabolite profiling data and report

OneOmics results files

## Acknowledgements

We thank the University of Minnesota Genomics Center for next generation sequencing data collection and Juan Abrahante (UMII) for assistance with RNAseq data analysis, and the University of Minnesota Center for Mass Spectrometry and Proteomics, Stephen Tate and Christie Hunter (SCIEX) for assistance with SWATH-MS data collection and analysis. This work was supported by the National Institutes of Health/National Cancer Institute (R01CA182546 and R33CA183671 to LLP). LJM was supported on the UMN Cancer Biology Training Grant (T32 CA009138) and the UMN Physical Sciences Oncology Center grant (U54CA210190).

## References cited

1. Babicki, S.; Arndt, D.; Marcu, A.; Liang, Y.; Grant, J. R.; Maciejewski, A.; Wishart, D. S., Heatmapper: web-enabled heat mapping for all. Nucleic Acids Res 2016, 44 (W1), W147–53.

2. Druker, B. J.; O’Brien, S. G.; Cortes, J.; Radich, J., Chronic myelogenous leukemia. Hematology / the Education Program of the American Society of Hematology. American Society of Hematology. Education Program 2002, 111–35.

3. Jabbour, E. J.; Cortes, J. E.; Kantarjian, H. M., Resistance to tyrosine kinase inhibition therapy for chronic myelogenous leukemia: a clinical perspective and emerging treatment options. Clinical lymphoma, myeloma & leukemia 2013, 13 (5), 515–29.

4. Carella, A. M.; Saglio, G.; Mahon, X. F.; Mauro, M. J., Present results and future perspectives in optimizing chronic myeloid leukemia therapy. Haematologica 2018, 103 (6), 928–930.

5. Sawyers, C. L., Perspective: combined forces. Nature 2013, 498 (7455), S7.

6. White, D.; Saunders, V.; Grigg, A.; Arthur, C.; Filshie, R.; Leahy, M. F.; Lynch, K.; To, L. B.; Hughes, T., Measurement of in vivo BCR-ABL kinase inhibition to monitor imatinib-induced target blockade and predict response in chronic myeloid leukemia. J Clin Oncol 2007, 25 (28), 4445–51.

7. Garg, R. J.; Kantarjian, H.; O’Brien, S.; Quintas-Cardama, A.; Faderl, S.; Estrov, Z.; Cortes, J., The use of nilotinib or dasatinib after failure to 2 prior tyrosine kinase inhibitors: long-term follow-up. Blood 2009, 114 (20), 4361–8.

8. Shah, N. P.; Rousselot, P.; Schiffer, C.; Rea, D.; Cortes, J. E.; Milone, J.; Mohamed, H.; Healey, D.; Kantarjian, H.; Hochhaus, A.; Saglio, G., Dasatinib in imatinib-resistant or - intolerant chronic-phase, chronic myeloid leukemia patients: 7-year follow-up of study CA180-034. American journal of hematology 2016, 91 (9), 869–74.

9. Cooper, M. J.; Cox, N. J.; Zimmerman, E. I.; Dewar, B. J.; Duncan, J. S.; Whittle, M. C.; Nguyen, T. A.; Jones, L. S.; Ghose Roy, S.; Smalley, D. M.; Kuan, P. F.; Richards, K. L.; Christopherson, R. I.; Jin, J.; Frye, S. V.; Johnson, G. L.; Baldwin, A. S.; Graves, L. M., Application of multiplexed kinase inhibitor beads to study kinome adaptations in drug-resistant leukemia. PloS one 2013, 8 (6), e66755.

10. Graves, L. M.; Duncan, J. S.; Whittle, M. C.; Johnson, G. L., The dynamic nature of the kinome. Biochem J 2013, 450 (1), 1–8.

11. Tarrado-Castellarnau, M.; de Atauri, P.; Cascante, M., Oncogenic regulation of tumor metabolic reprogramming. Oncotarget 2016, 7 (38), 62726–62753.

12. Herranz, D.; Ambesi-Impiombato, A.; Sudderth, J.; Sanchez-Martin, M.; Belver, L.; Tosello, V.; Xu, L.; Wendorff, A. A.; Castillo, M.; Haydu, J. E.; Marquez, J.; Mates, J. M.; Kung, A. L.; Rayport, S.; Cordon-Cardo, C.; DeBerardinis, R. J.; Ferrando, A. A., Metabolic reprogramming induces resistance to anti-NOTCH1 therapies in T cell acute lymphoblastic leukemia. Nat Med 2015, 21 (10), 1182–9.

13. Bhanot, H.; Weisberg, E. L.; Reddy, M. M.; Nonami, A.; Neuberg, D.; Stone, R. M.; Podar, K.; Salgia, R.; Griffin, J. D.; Sattler, M., Acute myeloid leukemia cells require 6- phosphogluconate dehydrogenase for cell growth and NADPH-dependent metabolic reprogramming. Oncotarget 2017, 8 (40), 67639–67650.

14. Lopes-Rodrigues, V.; Di Luca, A.; Mleczko, J.; Meleady, P.; Henry, M.; Pesic, M.; Cabrera, D.; van Liempd, S.; Lima, R. T.; O’Connor, R.; Falcon-Perez, J. M.; Vasconcelos, M. H., Identification of the metabolic alterations associated with the multidrug resistant phenotype in cancer and their intercellular transfer mediated by extracellular vesicles. Sci Rep 2017, 7, 44541.

15. Zhao, F.; Mancuso, A.; Bui, T. V.; Tong, X.; Gruber, J. J.; Swider, C. R.; Sanchez, P. V.; Lum, J. J.; Sayed, N.; Melo, J. V.; Perl, A. E.; Carroll, M.; Tuttle, S. W.; Thompson, C. B., Imatinib resistance associated with BCR-ABL upregulation is dependent on HIF-1alpha-induced metabolic reprograming. Oncogene 2010, 29 (20), 2962–72.

16. Edfors, F.; Danielsson, F.; Hallstrom, B. M.; Kall, L.; Lundberg, E.; Ponten, F.; Forsstrom, B.; Uhlen, M., Gene-specific correlation of RNA and protein levels in human cells and tissues. Mol Syst Biol 2016, 12 (10), 883.

17. Ouellette, S. B.; Noel, B. M.; Parker, L. L., A Cell-Based Assay for Measuring Endogenous BcrAbl Kinase Activity and Inhibitor Resistance. PloS one 2016, 11 (9), e0161748.

18. Trapnell, C.; Williams, B. A.; Pertea, G.; Mortazavi, A.; Kwan, G.; van Baren, M. J.; Salzberg, S. L.; Wold, B. J.; Pachter, L., Transcript assembly and quantification by RNA-Seq reveals unannotated transcripts and isoform switching during cell differentiation. Nat Biotechnol 2010, 28 (5), 511–5.

19. McPherson, A.; Hormozdiari, F.; Zayed, A.; Giuliany, R.; Ha, G.; Sun, M. G.; Griffith, M.; Heravi Moussavi, A.; Senz, J.; Melnyk, N.; Pacheco, M.; Marra, M. A.; Hirst, M.; Nielsen, T. O.; Sahinalp, S. C.; Huntsman, D.; Shah, S. P., deFuse: an algorithm for gene fusion discovery in tumor RNA-Seq data. PLoS computational biology 2011, 7 (5), e1001138.

20. Kumarakulasingham, M.; Rooney, P. H.; Dundas, S. R.; Telfer, C.; Melvin, W. T.; Curran, S.; Murray, G. I., Cytochrome p450 profile of colorectal cancer: identification of markers of prognosis. Clin Cancer Res 2005, 11 (10), 3758–65.

21. Gillet, L. C.; Navarro, P.; Tate, S.; Rost, H.; Selevsek, N.; Reiter, L.; Bonner, R.; Aebersold, R., Targeted data extraction of the MS/MS spectra generated by data-independent acquisition: a new concept for consistent and accurate proteome analysis. Mol Cell Proteomics 2012, 11 (6), O111 016717.

22. Kramer, A.; Green, J.; Pollard, J., Jr.; Tugendreich, S., Causal analysis approaches in Ingenuity Pathway Analysis. Bioinformatics 2014, 30 (4), 523–30.

23. Soga, T.; Heiger, D. N., Amino acid analysis by capillary electrophoresis electrospray ionization mass spectrometry. Anal Chem 2000, 72 (6), 1236–41.

24. Soga, T.; Ueno, Y.; Naraoka, H.; Ohashi, Y.; Tomita, M.; Nishioka, T., Simultaneous determination of anionic intermediates for Bacillus subtilis metabolic pathways by capillary electrophoresis electrospray ionization mass spectrometry. Anal Chem 2002, 74 (10), 2233–9.

25. Soga, T.; Ohashi, Y.; Ueno, Y.; Naraoka, H.; Tomita, M.; Nishioka, T., Quantitative metabolome analysis using capillary electrophoresis mass spectrometry. J Proteome Res 2003, 2 (5), 488–94.

26. Levin, J. Z.; Berger, M. F.; Adiconis, X.; Rogov, P.; Melnikov, A.; Fennell, T.; Nusbaum, C.; Garraway, L. A.; Gnirke, A., Targeted next-generation sequencing of a cancer transcriptome enhances detection of sequence variants and novel fusion transcripts. Genome biology 2009, 10 (10), R115.

27. Zhou, M. H.; Yang, Q. M., NUP214 fusion genes in acute leukemia (Review). Oncol Lett 2014, 8 (3), 959–962.

28. Eadie, L. N.; Dang, P.; Saunders, V. A.; Yeung, D. T.; Osborn, M. P.; Grigg, A. P.; Hughes, T. P.; White, D. L., The clinical significance of ABCB1 overexpression in predicting outcome of CML patients undergoing first-line imatinib treatment. Leukemia 2017, 31 (1), 75– 82.

29. Ye, H.; Adane, B.; Khan, N.; Sullivan, T.; Minhajuddin, M.; Gasparetto, M.; Stevens, B.; Pei, S.; Balys, M.; Ashton, J. M.; Klemm, D. J.; Woolthuis, C. M.; Stranahan, A. W.; Park, C. Y.; Jordan, C. T., Leukemic Stem Cells Evade Chemotherapy by Metabolic Adaptation to an Adipose Tissue Niche. Cell Stem Cell 2016, 19 (1), 23–37.

30. Zhou, H.; Mak, P. Y.; Mu, H.; Mak, D. H.; Zeng, Z.; Cortes, J.; Liu, Q.; Andreeff, M.; Carter, B. Z., Combined inhibition of beta-catenin and Bcr-Abl synergistically targets tyrosine kinase inhibitor-resistant blast crisis chronic myeloid leukemia blasts and progenitors in vitro and in vivo. Leukemia 2017, 31 (10), 2065–2074.

31. Grosso, S.; Puissant, A.; Dufies, M.; Colosetti, P.; Jacquel, A.; Lebrigand, K.; Barbry, P.; Deckert, M.; Cassuto, J. P.; Mari, B.; Auberger, P., Gene expression profiling of imatinib and PD166326-resistant CML cell lines identifies Fyn as a gene associated with resistance to BCR-ABL inhibitors. Mol Cancer Ther 2009, 8 (7), 1924–33.

32. Dufies, M.; Jacquel, A.; Belhacene, N.; Robert, G.; Cluzeau, T.; Luciano, F.; Cassuto, J. P.; Raynaud, S.; Auberger, P., Mechanisms of AXL overexpression and function in Imatinib-resistant chronic myeloid leukemia cells. Oncotarget 2011, 2 (11), 874–85.

33. Martinez-Outschoorn, U. E.; Peiris-Pages, M.; Pestell, R. G.; Sotgia, F.; Lisanti, M. P., Cancer metabolism: a therapeutic perspective. Nat Rev Clin Oncol 2017, 14 (1), 11–31.

34. Boulahbel, H.; Duran, R. V.; Gottlieb, E., Prolyl hydroxylases as regulators of cell metabolism. Biochemical Society transactions 2009, 37 (Pt 1), 291–4.

35. Hulikova, A.; Aveyard, N.; Harris, A. L.; Vaughan-Jones, R. D.; Swietach, P., Intracellular carbonic anhydrase activity sensitizes cancer cell pH signaling to dynamic changes in CO2 partial pressure. J Biol Chem 2014, 289 (37), 25418–30.

36. Swietach, P.; Vaughan-Jones, R. D.; Harris, A. L.; Hulikova, A., The chemistry, physiology and pathology of pH in cancer. Philos Trans R Soc Lond B Biol Sci 2014, 369 (1638), 20130099.

37. Norris, K. L.; Hao, R.; Chen, L. F.; Lai, C. H.; Kapur, M.; Shaughnessy, P. J.; Chou, D.; Yan, J.; Taylor, J. P.; Engelender, S.; West, A. E.; Lim, K. L.; Yao, T. P., Convergence of Parkin, PINK1, and alpha-Synuclein on Stress-induced Mitochondrial Morphological Remodeling. J Biol Chem 2015, 290 (22), 13862–74.

38. Rostovtseva, T. K.; Gurnev, P. A.; Protchenko, O.; Hoogerheide, D. P.; Yap, T. L.; Philpott, C. C.; Lee, J. C.; Bezrukov, S. M., alpha-Synuclein Shows High Affinity Interaction with Voltage-dependent Anion Channel, Suggesting Mechanisms of Mitochondrial Regulation and Toxicity in Parkinson Disease. J Biol Chem 2015, 290 (30), 18467–77.

39. Bir, A.; Sen, O.; Anand, S.; Khemka, V. K.; Banerjee, P.; Cappai, R.; Sahoo, A.; Chakrabarti, S., alpha-Synuclein-induced mitochondrial dysfunction in isolated preparation and intact cells: implications in the pathogenesis of Parkinson’s disease. J Neurochem 2014, 131 (6), 868–77.

40. Maitta, R. W.; Wolgast, L. R.; Wang, Q.; Zhang, H.; Bhattacharyya, P.; Gong, J. Z.; Sunkara, J.; Albanese, J. M.; Pizzolo, J. G.; Cannizzaro, L. A.; Ramesh, K. H.; Ratech, H., Alpha- and beta-synucleins are new diagnostic tools for acute erythroid leukemia and acute megakaryoblastic leukemia. Am J Hematol 2011, 86 (2), 230–4.

41. Woolthuis, C. M.; Park, C. Y., Hematopoietic stem/progenitor cell commitment to the megakaryocyte lineage. Blood 2016, 127 (10), 1242–8.

42. Pienimaeki-Roemer, A.; Kuhlmann, K.; Bottcher, A.; Konovalova, T.; Black, A.; Orso, E.; Liebisch, G.; Ahrens, M.; Eisenacher, M.; Meyer, H. E.; Schmitz, G., Lipidomic and proteomic characterization of platelet extracellular vesicle subfractions from senescent platelets. Transfusion 2015, 55 (3), 507–21.

43. Gottschalk, S.; Anderson, N.; Hainz, C.; Eckhardt, S. G.; Serkova, N. J., Imatinib (STI571)-mediated changes in glucose metabolism in human leukemia BCR-ABL-positive cells. Clin Cancer Res 2004, 10 (19), 6661–8.

44. Hirao, T.; Yamaguchi, M.; Kikuya, M.; Chibana, H.; Ito, K.; Aoki, S., Altered intracellular signaling by imatinib increases the anti-cancer effects of tyrosine kinase inhibitors in chronic myelogenous leukemia cells. Cancer science 2018, 109 (1), 121–131.

45. Kruger, A.; Gruning, N. M.; Wamelink, M. M.; Kerick, M.; Kirpy, A.; Parkhomchuk, D.; Bluemlein, K.; Schweiger, M. R.; Soldatov, A.; Lehrach, H.; Jakobs, C.; Ralser, M., The pentose phosphate pathway is a metabolic redox sensor and regulates transcription during the antioxidant response. Antioxidants & redox signaling 2011, 15 (2), 311–24.

46. Gruning, N. M.; Rinnerthaler, M.; Bluemlein, K.; Mulleder, M.; Wamelink, M. M.; Lehrach, H.; Jakobs, C.; Breitenbach, M.; Ralser, M., Pyruvate kinase triggers a metabolic feedback loop that controls redox metabolism in respiring cells. Cell metabolism 2011, 14 (3), 415–27.

47. Patra, K. C.; Hay, N., The pentose phosphate pathway and cancer. Trends Biochem Sci 2014, 39 (8), 347–54.

48. Ibsen, K. H., The Crabtree effect: a review. Cancer Res 1961, 21, 829–41.

49. Robinson, B. H.; Petrova-Benedict, R.; Buncic, J. R.; Wallace, D. C., Nonviability of cells with oxidative defects in galactose medium: a screening test for affected patient fibroblasts. Biochem Med Metab Biol 1992, 48 (2), 122–6.

50. Arroyo, J. D.; Jourdain, A. A.; Calvo, S. E.; Ballarano, C. A.; Doench, J. G.; Root, D. E.; Mootha, V. K., A Genome-wide CRISPR Death Screen Identifies Genes Essential for Oxidative Phosphorylation. Cell metabolism 2016, 24 (6), 875–885.

51. Pelley, J. W., 9 - Minor Carbohydrate Pathways: Ribose, Fructose, and Galactose. In Elsevier’s Integrated Biochemistry, Mosby: Philadelphia, 2007; pp 73–77.

52. Calvo, S. E.; Clauser, K. R.; Mootha, V. K., MitoCarta2.0: an updated inventory of mammalian mitochondrial proteins. Nucleic Acids Res 2016, 44 (D1), D1251–7.

53. Ben-Batalla, I.; Erdmann, R.; Jorgensen, H.; Mitchell, R.; Ernst, T.; von Amsberg, G.; Schafhausen, P.; Velthaus, J. L.; Rankin, S.; Clark, R. E.; Koschmieder, S.; Schultze, A.; Mitra, S.; Vandenberghe, P.; Brummendorf, T. H.; Carmeliet, P.; Hochhaus, A.; Pantel, K.; Bokemeyer, C.; Helgason, G. V.; Holyoake, T. L.; Loges, S., Axl Blockade by BGB324 Inhibits BCR-ABL Tyrosine Kinase Inhibitor-Sensitive and -Resistant Chronic Myeloid Leukemia. Clin Cancer Res 2017, 23 (9), 2289–2300.

54. Schito, L.; Semenza, G. L., Hypoxia-Inducible Factors: Master Regulators of Cancer Progression. Trends Cancer 2016, 2 (12), 758–770.

55. Vidal, R. S.; Quarti, J.; Rumjanek, F. D.; Rumjanek, V. M., Metabolic Reprogramming During Multidrug Resistance in Leukemias. Front Oncol 2018, 8, 90.

56. Kim, H. K.; Noh, Y. H.; Nilius, B.; Ko, K. S.; Rhee, B. D.; Kim, N.; Han, J., Current and upcoming mitochondrial targets for cancer therapy. Semin Cancer Biol 2017, 47, 154–167.

57. Samudio, I.; Fiegl, M.; Andreeff, M., Mitochondrial uncoupling and the Warburg effect: molecular basis for the reprogramming of cancer cell metabolism. Cancer Res 2009, 69 (6), 2163–6.

58. Sack, M. N., Mitochondrial depolarization and the role of uncoupling proteins in ischemia tolerance. Cardiovasc Res 2006, 72 (2), 210–9.

59. Amoedo, N. D.; Punzi, G.; Obre, E.; Lacombe, D.; De Grassi, A.; Pierri, C. L.; Rossignol, R., AGC1/2, the mitochondrial aspartate-glutamate carriers. Biochim Biophys Acta 2016, 1863 (10), 2394–412.

60. Strickland, M.; Stoll, E. A., Metabolic Reprogramming in Glioma. Front Cell Dev Biol 2017, 5, 43.

61. Allegra, A.; Innao, V.; Gerace, D.; Bianco, O.; Musolino, C., The metabolomic signature of hematologic malignancies. Leuk Res 2016, 49, 22–35.

62. Bosc, C.; Selak, M. A.; Sarry, J. E., Resistance Is Futile: Targeting Mitochondrial Energetics and Metabolism to Overcome Drug Resistance in Cancer Treatment. Cell metabolism 2017, 26 (5), 705–707.

63. Farge, T.; Saland, E.; de Toni, F.; Aroua, N.; Hosseini, M.; Perry, R.; Bosc, C.; Sugita, M.; Stuani, L.; Fraisse, M.; Scotland, S.; Larrue, C.; Boutzen, H.; Feliu, V.; Nicolau-Travers, M. L.; Cassant-Sourdy, S.; Broin, N.; David, M.; Serhan, N.; Sarry, A.; Tavitian, S.; Kaoma, T.; Vallar, L.; Iacovoni, J.; Linares, L. K.; Montersino, C.; Castellano, R.; Griessinger, E.; Collette, Y.; Duchamp, O.; Barreira, Y.; Hirsch, P.; Palama, T.; Gales, L.; Delhommeau, F.; Garmy-Susini, B. H.; Portais, J. C.; Vergez, F.; Selak, M.; Danet-Desnoyers, G.; Carroll, M.; Recher, C.; Sarry, J. E., Chemotherapy-Resistant Human Acute Myeloid Leukemia Cells Are Not Enriched for Leukemic Stem Cells but Require Oxidative Metabolism. Cancer discovery 2017, 7 (7), 716–735.

64. Miwa, H.; Shikami, M.; Goto, M.; Mizuno, S.; Takahashi, M.; Tsunekawa-Imai, N.; Ishikawa, T.; Mizutani, M.; Horio, T.; Gotou, M.; Yamamoto, H.; Wakabayashi, M.; Watarai, M.; Hanamura, I.; Imamura, A.; Mihara, H.; Nitta, M., Leukemia cells demonstrate a different metabolic perturbation provoked by 2-deoxyglucose. Oncology reports 2013, 29 (5), 2053–7.

