## Supplementary material for "Multi-omic profiling of tyrosine kinase inhibitor-resistant K562 cells suggests metabolic reprogramming to promote cell survival": Exosome supplemental file: 170126_DREXosome1-001-BATCH-report.pdf

### BATCH ANALYSIS REPORT

NANOSIGHT

Nanoparticle Tracking Analysis (NTA) Version 2.3 Build 0034

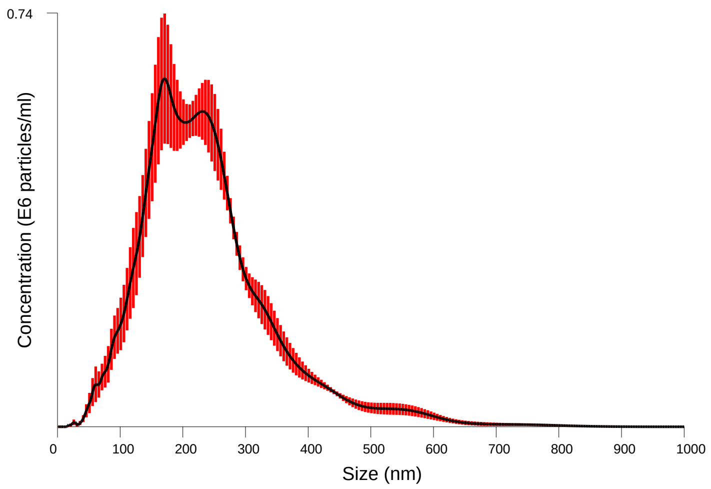


Averaged Size / Concentration
Red error bars indicate +/- 1 standard error of the mean

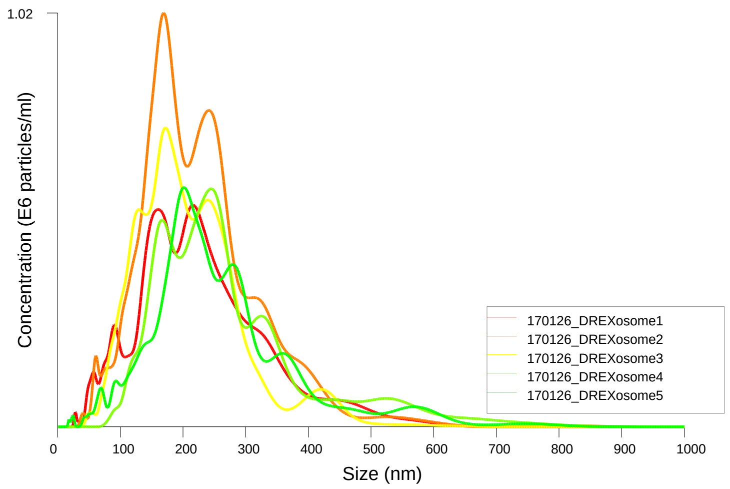


Size / Concentration

| | |
|---|---|
| Included files: | 170126_DREXosome1 |
| | 170126_DREXosome2 |
| | 170126_DREXosome3 |
| | 170126_DREXosome4 |
| | 170126_DREXosome5 |
| Date/Time of Report | 26/01/2017 17:25:27 |

**BATCH AVERAGE RESULTS:**

| | |
|---|---|
| Size Distribution: | Mean: 242 +/- 11.9 nm |
| | Mode: 200 +/- 14.2 nm |
| | SD: 106 +/- 7.6 nm |
| | D10: 128 +/- 7.5 nm |
| | D50: 222 +/- 8.7 nm |
| | D90: 382 +/- 26.0 nm |
| Total Concentration: | 10.72 +/- 0.96 particles/frame |
| | 1.25 +/- 0.11 E8 particles/ml |
| Total Completed Tracks: | 2467 |
| Average Drift Velocity: | 192 nm/s |

**CAPTURE SETTINGS**

| | |
|---|---|
| Camera Type: | sCMOS |
| Shutter length: | Varied |
| Shutter setting: | Varied |
| Camera gain: | 350 |
| Frame rate: | Varied |

**ANALYSIS SETTINGS**

| | |
|---|---|
| Background Extract: | On |
| Detection Threshold: | 28 - Multi |
| Blur: | AUTO |
| Min track length: | AUTO |
| Min expected size: | AUTO |
| Temperature: | 22.0, 22.0, 22.0, 22.0, 22.0 °C |
| Viscosity: | 0.95, 0.95, 0.95, 0.95, 0.95 cP |

**WARNINGS**

Vibration detected during analysis
