## Supplementary material for "Multi-omic profiling of tyrosine kinase inhibitor-resistant K562 cells suggests metabolic reprogramming to promote cell survival": Exosome supplemental file: 170127_NRExosome1-001-BATCH-report.pdf

NANOSIGHT

### BATCH ANALYSIS REPORT

Nanoparticle Tracking Analysis (NTA) Version 2.3 Build 0034

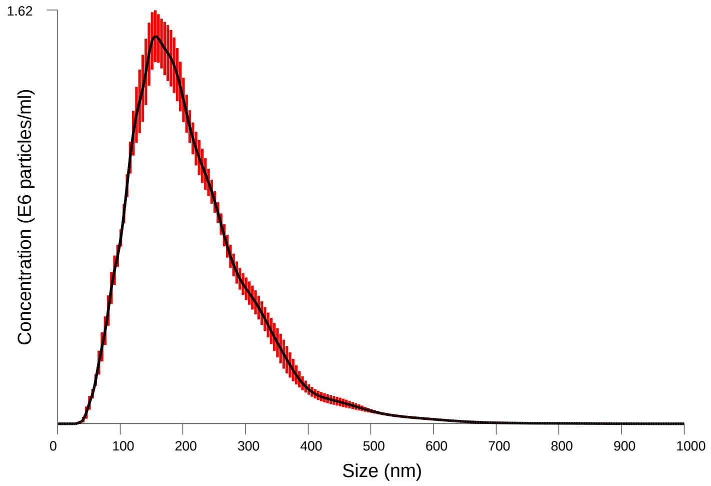


Averaged Size / Concentration
Red error bars indicate +/- 1 standard error of the mean

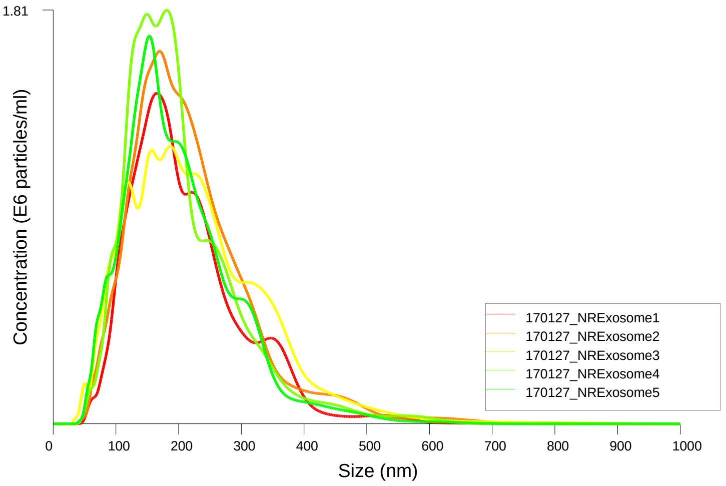


Size / Concentration

| | |
|---|---|
| Included files: | 170127_NRExosome1 |
| | 170127_NRExosome2 |
| | 170127_NRExosome3 |
| | 170127_NRExosome4 |
| | 170127_NRExosome5 |
| Date/Time of Report | 27/01/2017 20:13:00 |

**BATCH AVERAGE RESULTS:**

| | |
|---|---|
| Size Distribution: | Mean: 213 +/- 5.1 nm |
| | Mode: 170 +/- 6.0 nm |
| | SD: 95 +/- 3.8 nm |
| | D10: 110 +/- 2.5 nm |
| | D50: 194 +/- 5.1 nm |
| | D90: 335 +/- 9.3 nm |
| Total Concentration: | 23.89 +/- 0.85 particles/frame |
| | 2.82 +/- 0.10 E8 particles/ml |
| Total Completed Tracks: | 6358 |
| Average Drift Velocity: | 101 nm/s |

**CAPTURE SETTINGS**

| | |
|---|---|
| Camera Type: | sCMOS |
| Shutter length: | 6.245 ms |
| Shutter setting: | 250 |
| Camera gain: | 250 |
| Frame rate: | Varied |

**ANALYSIS SETTINGS**

| | |
|---|---|
| Background Extract: | On |
| Detection Threshold: | 10 - Multi |
| Blur: | AUTO |
| Min track length: | AUTO |
| Min expected size: | AUTO |
| Temperature: | 22.0, 22.0, 22.0, 22.0, 22.0 ºC |
| Viscosity: | 0.95, 0.95, 0.95, 0.95, 0.95 cP |

**WARNINGS**

Vibration detected during analysis
