## Supplementary material for "Multi-omic profiling of tyrosine kinase inhibitor-resistant K562 cells suggests metabolic reprogramming to promote cell survival": Exosome supplemental file: 170201_DRExosome1-001-BATCH-report.pdf

### BATCH ANALYSIS REPORT

Nanoparticle Tracking Analysis (NTA) Version 2.3 Build 0034

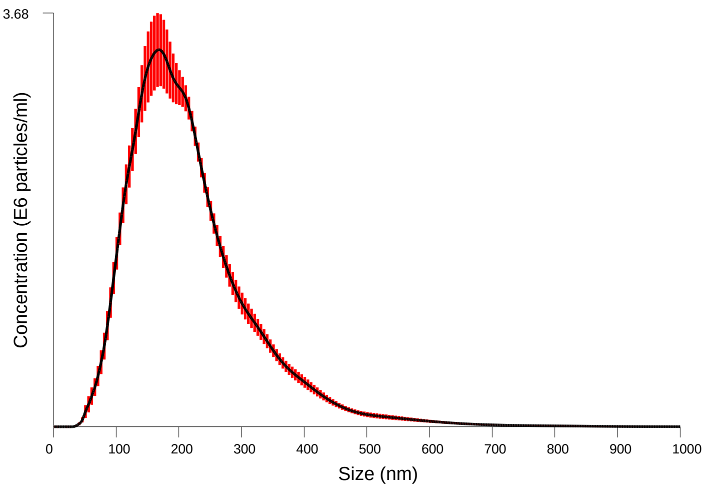


Averaged Size / Concentration
Red error bars indicate +/- 1 standard error of the mean

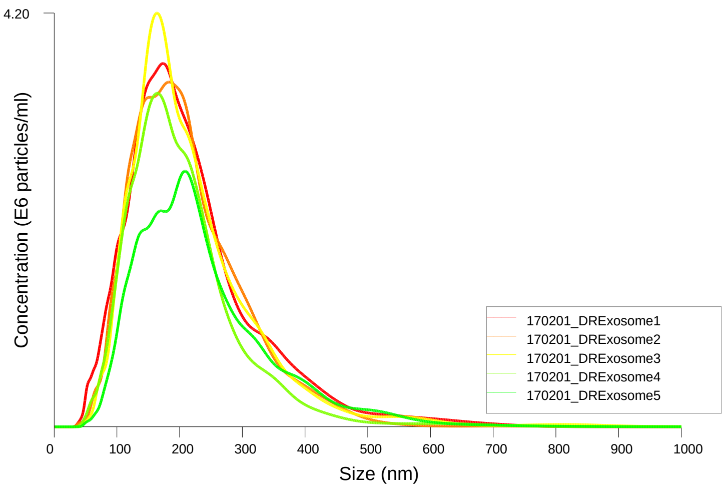


Size / Concentration

| | |
|---|---|
| Included files: | 170201_DRExosome1 |
| | 170201_DRExosome2 |
| | 170201_DRExosome3 |
| | 170201_DRExosome4 |
| | 170201_DRExosome5 |
| Date/Time of Report | 01/02/2017 17:41:34 |

**BATCH AVERAGE RESULTS:**

| | |
|---|---|
| Size Distribution: | Mean: 219 +/- 5.3 nm |
| | Mode: 178 +/- 8.3 nm |
| | SD: 100 +/- 3.9 nm |
| | D10: 116 +/- 2.3 nm |
| | D50: 198 +/- 4.5 nm |
| | D90: 347 +/- 11.7 nm |
| Total Concentration: | 51.12 +/- 2.93 particles/frame |
| | 6.24 +/- 0.39 E8 particles/ml |
| Total Completed Tracks: | 14500 |
| Average Drift Velocity: | 3200 nm/s |

**CAPTURE SETTINGS**

| | |
|---|---|
| Camera Type: | sCMOS |
| Shutter length: | 11.2357 ms |
| Shutter setting: | 450 |
| Camera gain: | 250 |
| Frame rate: | Varied |

**ANALYSIS SETTINGS**

| | |
|---|---|
| Background Extract: | On |
| Detection Threshold: | 10 - Multi |
| Blur: | AUTO |
| Min track length: | AUTO |
| Min expected size: | AUTO |
| Temperature: | 22.0, 22.0, 22.0, 22.0, 22.0 ºC |
| Viscosity: | 0.95, 0.95, 0.95, 0.95, 0.95 cP |

**WARNINGS**

Vibration detected during analysis
