## Supplementary material for "Multi-omic profiling of tyrosine kinase inhibitor-resistant K562 cells suggests metabolic reprogramming to promote cell survival": Exosome supplemental file: 170201_IRExosome1-001-BATCH-report.pdf

### BATCH ANALYSIS REPORT

Nanoparticle Tracking Analysis (NTA) Version 2.3 Build 0034

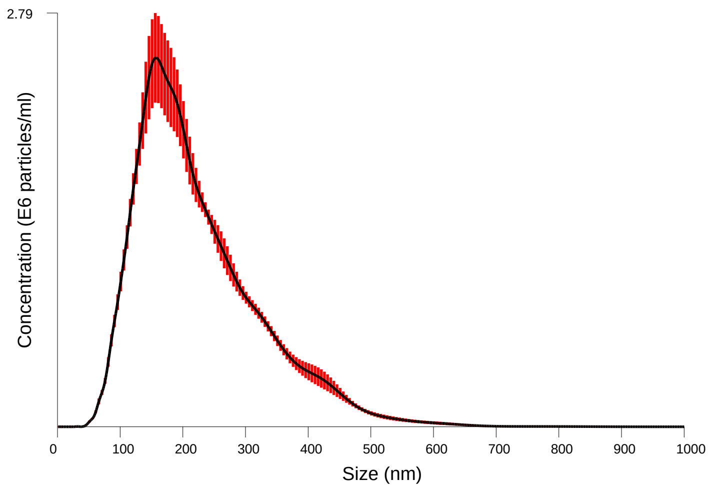


Averaged Size / Concentration
Red error bars indicate +/- 1 standard error of the mean

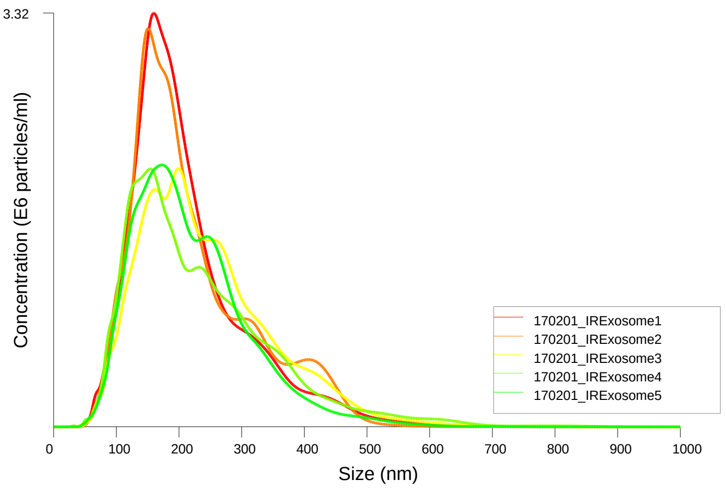


Size / Concentration

| | |
|---|---|
| Included files: | 170201_IRExosome1 |
| | 170201_IRExosome2 |
| | 170201_IRExosome3 |
| | 170201_IRExosome4 |
| | 170201_IRExosome5 |
| Date/Time of Report | 01/02/2017 17:27:07 |

**BATCH AVERAGE RESULTS:**

| | |
|---|---|
| Size Distribution: | Mean: 225 +/- 5.3 nm |
| | Mode: 166 +/- 8.8 nm |
| | SD: 98 +/- 4.8 nm |
| | D10: 120 +/- 1.9 nm |
| | D50: 201 +/- 5.6 nm |
| | D90: 364 +/- 10.4 nm |
| Total Concentration: | 37.51 +/- 1.70 particles/frame |
| | 4.45 +/- 0.19 E8 particles/ml |
| Total Completed Tracks: | 9720 |
| Average Drift Velocity: | 111 nm/s |

**CAPTURE SETTINGS**

| | |
|---|---|
| Camera Type: | sCMOS |
| Shutter length: | 11.2357 ms |
| Shutter setting: | 450 |
| Camera gain: | 250 |
| Frame rate: | Varied |

**ANALYSIS SETTINGS**

| | |
|---|---|
| Background Extract: | On |
| Detection Threshold: | 10 - Multi |
| Blur: | AUTO |
| Min track length: | AUTO |
| Min expected size: | AUTO |
| Temperature: | 22.0, 22.0, 22.0, 22.0, 22.0 $^{o}C$ |
| Viscosity: | 0.95, 0.95, 0.95, 0.95, 0.95 cP |

**WARNINGS**

Vibration detected during analysis
