## Supplementary material for "Multi-omic profiling of tyrosine kinase inhibitor-resistant K562 cells suggests metabolic reprogramming to promote cell survival": Exosome supplemental file: 170206_NRExosome1-001-BATCH-report.pdf

NANOSIGHT

### BATCH ANALYSIS REPORT

Nanoparticle Tracking Analysis (NTA) Version 2.3 Build 0034

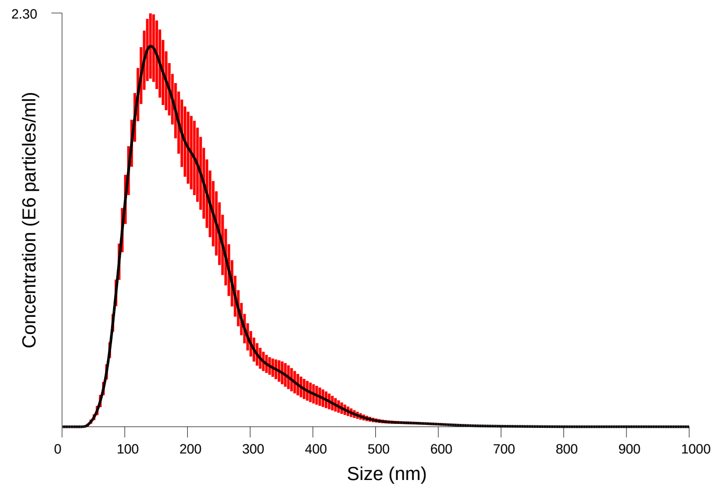


Averaged Size / Concentration
Red error bars indicate +/- 1 standard error of the mean

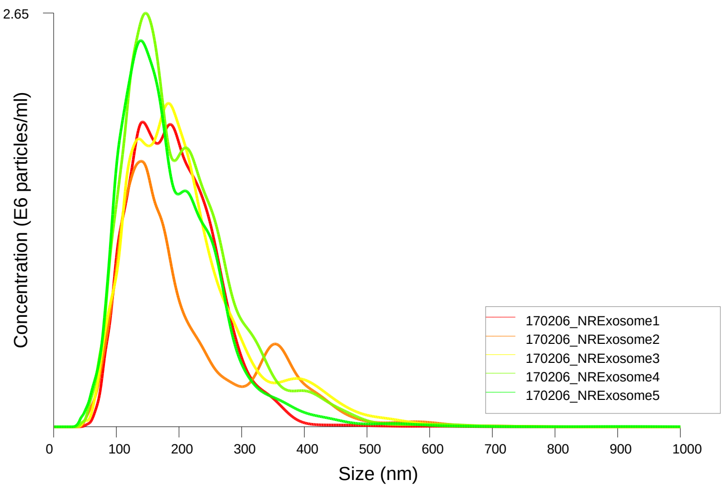


Size / Concentration

| | |
|---|---|
| Included files: | 170206_NRExosome1 |
| | 170206_NRExosome2 |
| | 170206_NRExosome3 |
| | 170206_NRExosome4 |
| | 170206_NRExosome5 |
| Date/Time of Report | 06/02/2017 20:48:29 |

**BATCH AVERAGE RESULTS:**

| | |
|---|---|
| Size Distribution: | Mean: 201 +/- 5.2 nm |
| | Mode: 149 +/- 8.3 nm |
| | SD: 88 +/- 7.3 nm |
| | D10: 107 +/- 1.9 nm |
| | D50: 180 +/- 4.3 nm |
| | D90: 320 +/- 19.2 nm |
| Total Concentration: | 29.97 +/- 2.29 particles/frame |
| | 3.56 +/- 0.28 E8 particles/ml |
| Total Completed Tracks: | 7582 |
| Average Drift Velocity: | 76 nm/s |

**CAPTURE SETTINGS**

| | |
|---|---|
| Camera Type: | sCMOS |
| Shutter length: | 14.9987 ms |
| Shutter setting: | 600 |
| Camera gain: | 300 |
| Frame rate: | Varied |

**ANALYSIS SETTINGS**

| | |
|---|---|
| Background Extract: | On |
| Detection Threshold: | 10 - Multi |
| Blur: | AUTO |
| Min track length: | AUTO |
| Min expected size: | AUTO |
| Temperature: | 22.0, 22.0, 22.0, 22.0, 22.0 °C |
| Viscosity: | 0.95, 0.95, 0.95, 0.95, 0.95 cP |

**WARNINGS**

Vibration detected during analysis
