## Supplementary material for "Multi-omic profiling of tyrosine kinase inhibitor-resistant K562 cells suggests metabolic reprogramming to promote cell survival": Exosome supplemental file: 170206_WTExosome1-001-BATCH-report.pdf

### NANOSIGHT

### BATCH ANALYSIS REPORT

Nanoparticle Tracking Analysis (NTA) Version 2.3 Build 0034

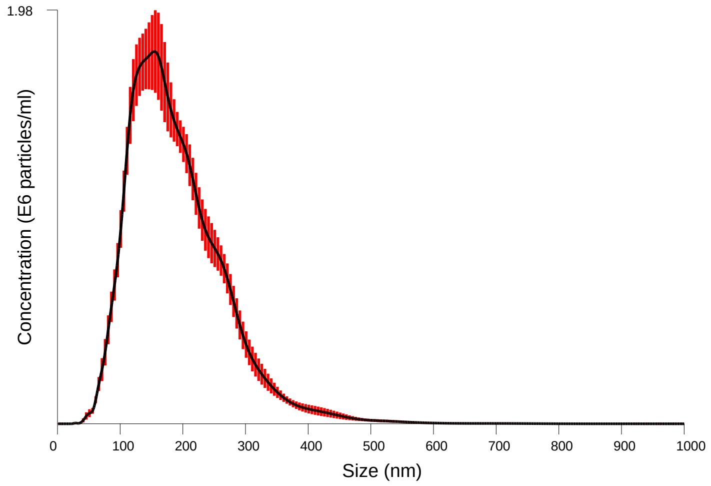


Averaged Size / Concentration
Red error bars indicate +/- 1 standard error of the mean

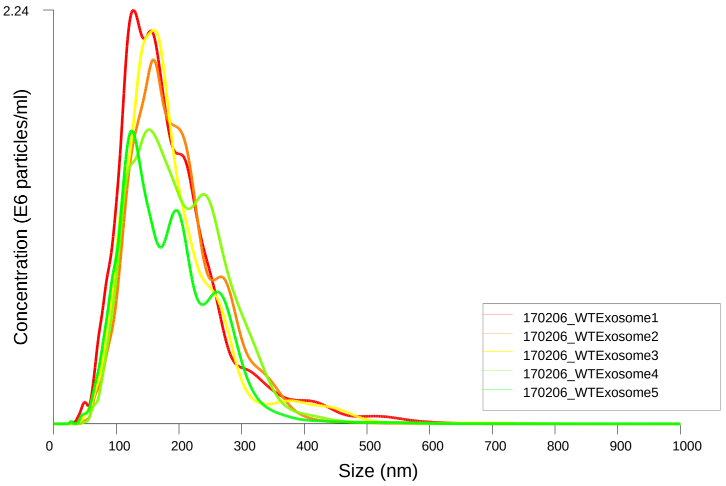


Size / Concentration

| | |
|---|---|
| Included files: | 170206_WTExosome1 |
| | 170206_WTExosome2 |
| | 170206_WTExosome3 |
| | 170206_WTExosome4 |
| | 170206_WTExosome5 |
| Date/Time of Report | 06/02/2017 20:36:57 |

**BATCH AVERAGE RESULTS:**

| | |
|---|---|
| Size Distribution: | Mean: 190 +/- 3.2 nm |
| | Mode: 144 +/- 7.7 nm |
| | SD: 75 +/- 2.8 nm |
| | D10: 108 +/- 2.7 nm |
| | D50: 176 +/- 4.4 nm |
| | D90: 286 +/- 3.7 nm |
| Total Concentration: | 24.18 +/- 1.44 particles/frame |
| | 2.83 +/- 0.17 E8 particles/ml |
| Total Completed Tracks: | 5871 |
| Average Drift Velocity: | 186 nm/s |

**CAPTURE SETTINGS**

| | |
|---|---|
| Camera Type: | sCMOS |
| Shutter length: | 14.9987 ms |
| Shutter setting: | 600 |
| Camera gain: | 300 |
| Frame rate: | Varied |

**ANALYSIS SETTINGS**

| | |
|---|---|
| Background Extract: | On |
| Detection Threshold: | 10 - Multi |
| Blur: | AUTO |
| Min track length: | AUTO |
| Min expected size: | AUTO |
| Temperature: | 22.0, 22.0, 22.0, 22.0, 22.0 ºC |
| Viscosity: | 0.95, 0.95, 0.95, 0.95, 0.95 cP |

**WARNINGS**

Vibration detected during analysis
