## Supplementary material for "Multi-omic profiling of tyrosine kinase inhibitor-resistant K562 cells suggests metabolic reprogramming to promote cell survival": Exosome supplemental file: 170207_IRExosome1-001-BATCH-report.pdf

### BATCH ANALYSIS REPORT

Nanoparticle Tracking Analysis (NTA) Version 2.3 Build 0034

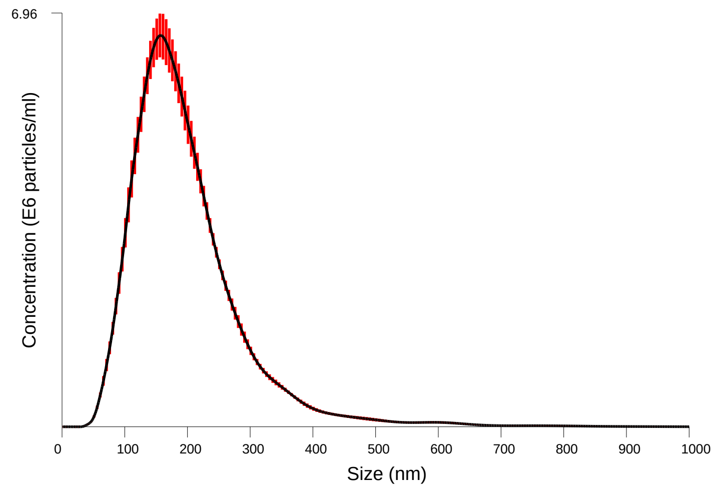


Averaged Size / Concentration
Red error bars indicate +/- 1 standard error of the mean

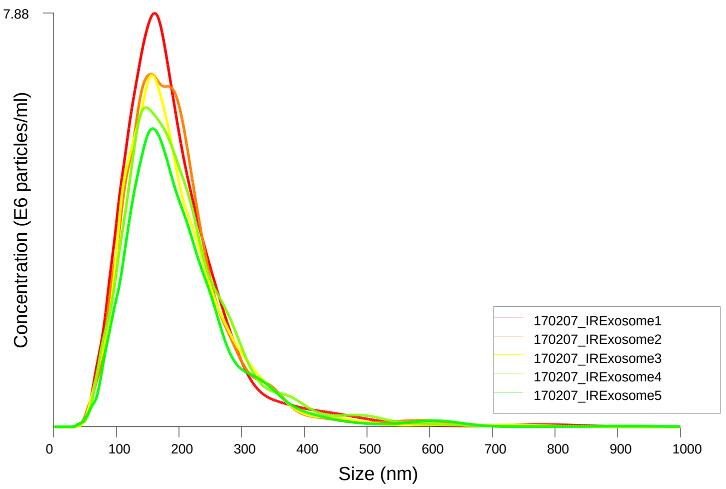


Size / Concentration

| | |
|---|---|
| Included files: | 170207_IRExosome1 |
| | 170207_IRExosome2 |
| | 170207_IRExosome3 |
| | 170207_IRExosome4 |
| | 170207_IRExosome5 |
| Date/Time of Report | 07/02/2017 16:17:36 |

**BATCH AVERAGE RESULTS:**

| | |
|---|---|
| Size Distribution: | Mean: 198 +/- 1.9 nm |
| | Mode: 155 +/- 2.4 nm |
| | SD: 93 +/- 1.8 nm |
| | D10: 108 +/- 0.9 nm |
| | D50: 179 +/- 1.7 nm |
| | D90: 301 +/- 4.6 nm |
| Total Concentration: | 83.62 +/- 3.68 particles/frame |
| | 10.29 +/- 0.46 E8 particles/ml |
| Total Completed Tracks: | 23575 |
| Average Drift Velocity: | 228 nm/s |

**CAPTURE SETTINGS**

| | |
|---|---|
| Camera Type: | sCMOS |
| Shutter length: | 11.2357 ms |
| Shutter setting: | 450 |
| Camera gain: | 250 |
| Frame rate: | Varied |

**ANALYSIS SETTINGS**

| | |
|---|---|
| Background Extract: | On |
| Detection Threshold: | 10 - Multi |
| Blur: | AUTO |
| Min track length: | AUTO |
| Min expected size: | AUTO |
| Temperature: | 22.0, 22.0, 22.0, 22.0, 22.0 °C |
| Viscosity: | 0.95, 0.95, 0.95, 0.95, 0.95 cP |

**WARNINGS**

Vibration detected during analysis
