## Supplementary material for "Multi-omic profiling of tyrosine kinase inhibitor-resistant K562 cells suggests metabolic reprogramming to promote cell survival": Exosome supplemental file: 170207_NRExosome1-001-BATCH-report.pdf

NANOSIGHT

Nanoparticle Tracking Analysis (NTA) Version 2.3 Build 0034

### BATCH ANALYSIS REPORT

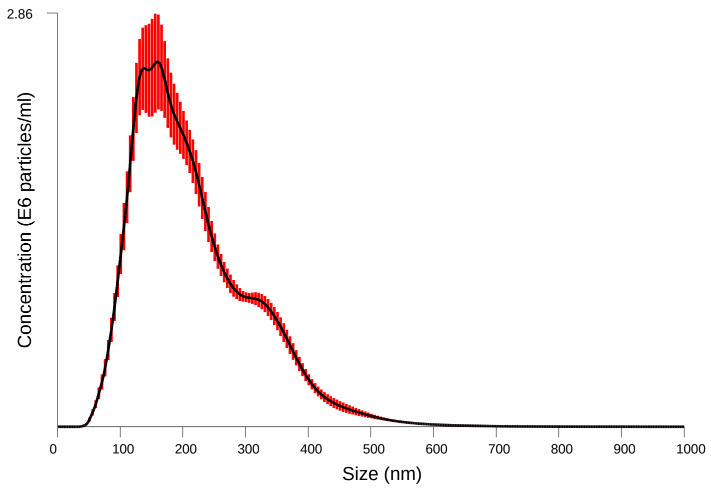


Averaged Size / Concentration
Red error bars indicate +/- 1 standard error of the mean

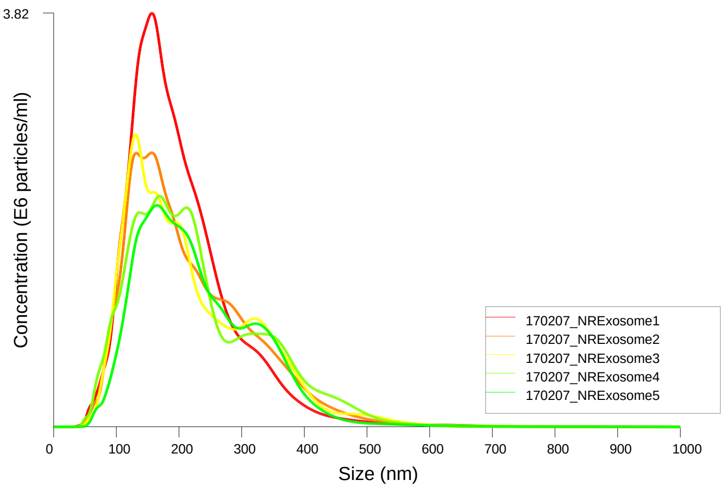


Size / Concentration

| | |
|---|---|
| Included files: | 170207_NRExosome1 |
| | 170207_NRExosome2 |
| | 170207_NRExosome3 |
| | 170207_NRExosome4 |
| | 170207_NRExosome5 |
| Date/Time of Report | 07/02/2017 16:24:14 |

**BATCH AVERAGE RESULTS:**

| | |
|---|---|
| Size Distribution: | Mean: 218 +/- 4.9 nm |
| | Mode: 155 +/- 6.6 nm |
| | SD: 94 +/- 3.8 nm |
| | D10: 115 +/- 2.5 nm |
| | D50: 197 +/- 4.8 nm |
| | D90: 349 +/- 10.6 nm |
| Total Concentration: | 38.84 +/- 1.98 particles/frame |
| | 4.66 +/- 0.25 E8 particles/ml |
| Total Completed Tracks: | 10006 |
| Average Drift Velocity: | 204 nm/s |

**CAPTURE SETTINGS**

| | |
|---|---|
| Camera Type: | sCMOS |
| Shutter length: | 11.2357 ms |
| Shutter setting: | 450 |
| Camera gain: | 250 |
| Frame rate: | Varied |

**ANALYSIS SETTINGS**

| | |
|---|---|
| Background Extract: | On |
| Detection Threshold: | 10 - Multi |
| Blur: | AUTO |
| Min track length: | AUTO |
| Min expected size: | AUTO |
| Temperature: | 22.0, 22.0, 22.0, 22.0, 22.0 ºC |
| Viscosity: | 0.95, 0.95, 0.95, 0.95, 0.95 cP |

**WARNINGS**

Vibration detected during analysis
