## Supplementary material for "Multi-omic profiling of tyrosine kinase inhibitor-resistant K562 cells suggests metabolic reprogramming to promote cell survival": Exosome supplemental file: 170207_WTExosome1-001-BATCH-report.pdf

### BATCH ANALYSIS REPORT

Nanoparticle Tracking Analysis (NTA) Version 2.3 Build 0034

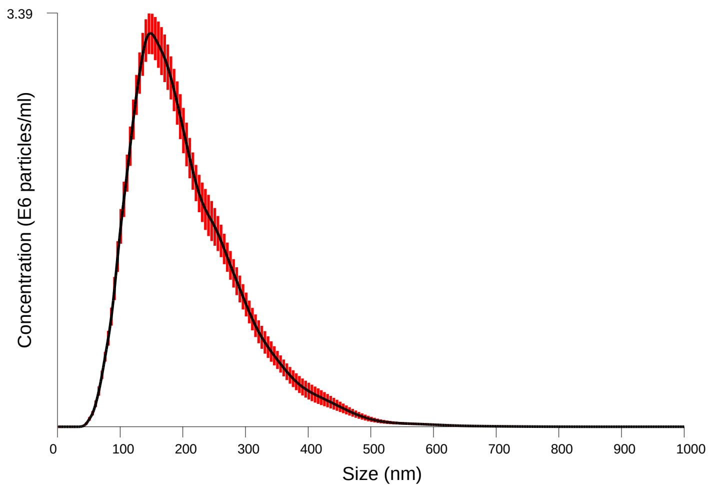


Averaged Size / Concentration
Red error bars indicate +/- 1 standard error of the mean

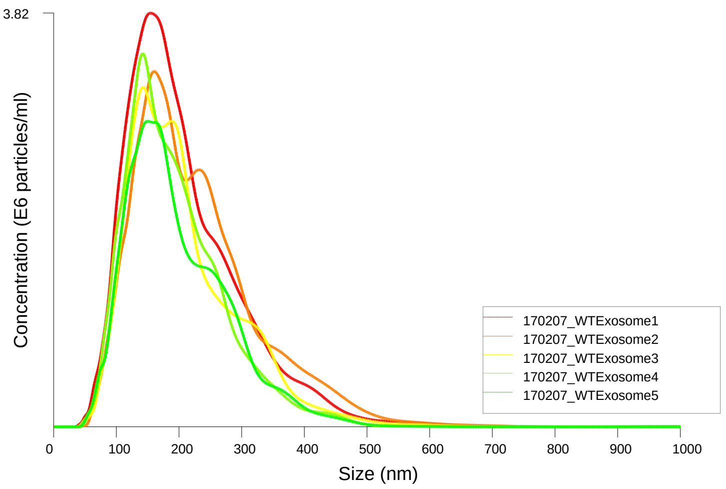


Size / Concentration

| | |
|---|---|
| Included files: | 170207_WTExosome1 |
| | 170207_WTExosome2 |
| | 170207_WTExosome3 |
| | 170207_WTExosome4 |
| | 170207_WTExosome5 |
| Date/Time of Report | 07/02/2017 16:11:41 |

**BATCH AVERAGE RESULTS:**

| | |
|---|---|
| Size Distribution: | Mean: 206 +/- 5.6 nm |
| | Mode: 149 +/- 3.5 nm |
| | SD: 87 +/- 3.7 nm |
| | D10: 112 +/- 2.1 nm |
| | D50: 187 +/- 5.1 nm |
| | D90: 325 +/- 11.9 nm |
| Total Concentration: | 46.57 +/- 2.87 particles/frame |
| | 5.51 +/- 0.35 E8 particles/ml |
| Total Completed Tracks: | 13203 |
| Average Drift Velocity: | 199 nm/s |

**CAPTURE SETTINGS**

| | |
|---|---|
| Camera Type: | sCMOS |
| Shutter length: | 11.2357 ms |
| Shutter setting: | 450 |
| Camera gain: | 250 |
| Frame rate: | Varied |

**ANALYSIS SETTINGS**

| | |
|---|---|
| Background Extract: | On |
| Detection Threshold: | 10 - Multi |
| Blur: | AUTO |
| Min track length: | AUTO |
| Min expected size: | AUTO |
| Temperature: | 22.0, 22.0, 22.0, 22.0, 22.0 ºC |
| Viscosity: | 0.95, 0.95, 0.95, 0.95, 0.95 cP |

**WARNINGS**

Vibration detected during analysis
