## Supplementary material for "Multi-omic profiling of tyrosine kinase inhibitor-resistant K562 cells suggests metabolic reprogramming to promote cell survival": Exosome supplemental file: 170127_blank1-002-BATCH-report.pdf

NANOSIGHT

### BATCH ANALYSIS REPORT

Nanoparticle Tracking Analysis (NTA) Version 2.3 Build 0034

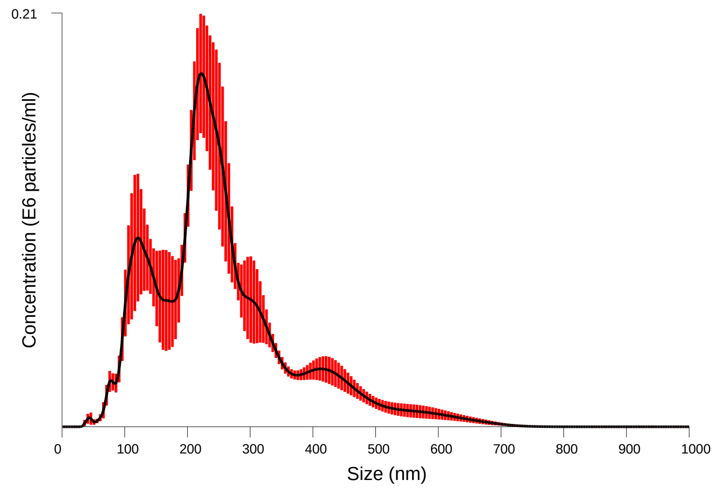


Averaged Size / Concentration
Red error bars indicate +/- 1 standard error of the mean

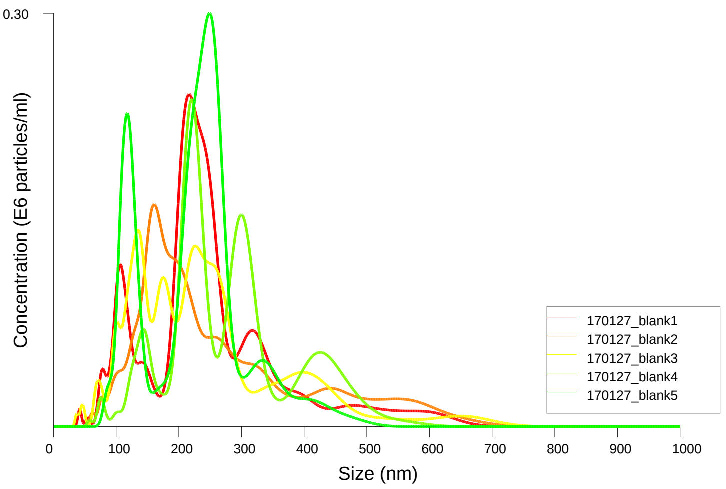


Size / Concentration

| | |
|---|---|
| Included files: | 170127_blank1 |
| | 170127_blank2 |
| | 170127_blank3 |
| | 170127_blank4 |
| | 170127_blank5 |
| Date/Time of Report | 27/01/2017 20:17:50 |

**BATCH AVERAGE RESULTS:**

| | |
|---|---|
| Size Distribution: | Mean: 259 +/- 9.6 nm |
| | Mode: 195 +/- 21.0 nm |
| | SD: 123 +/- 15.2 nm |
| | D10: 126 +/- 8.1 nm |
| | D50: 235 +/- 7.5 nm |
| | D90: 428 +/- 27.1 nm |
| Total Concentration: | 2.53 +/- 0.11 particles/frame |
| | 0.29 +/- 0.01 E8 particles/ml |
| Total Completed Tracks: | 451 |
| Average Drift Velocity: | 348 nm/s |

**CAPTURE SETTINGS**

| | |
|---|---|
| Camera Type: | sCMOS |
| Shutter length: | 14.9987 ms |
| Shutter setting: | 600 |
| Camera gain: | 250 |
| Frame rate: | Varied |

**ANALYSIS SETTINGS**

| | |
|---|---|
| Background Extract: | On |
| Detection Threshold: | 10 - Multi |
| Blur: | AUTO |
| Min track length: | AUTO |
| Min expected size: | AUTO |
| Temperature: | 22.0, 22.0, 22.0, 22.0, 22.0 ºC |
| Viscosity: | 0.95, 0.95, 0.95, 0.95, 0.95 cP |

**WARNINGS**

Vibration detected during analysis

Auto max jump failed in some analyses
