## Supplementary material for "Multi-omic profiling of tyrosine kinase inhibitor-resistant K562 cells suggests metabolic reprogramming to promote cell survival": Exosome supplemental file: 170208extraction buffer Control1-001-BATCH-report.pdf

### NANOSIGHT

### BATCH ANALYSIS REPORT

Nanoparticle Tracking Analysis (NTA) Version 2.3 Build 0034

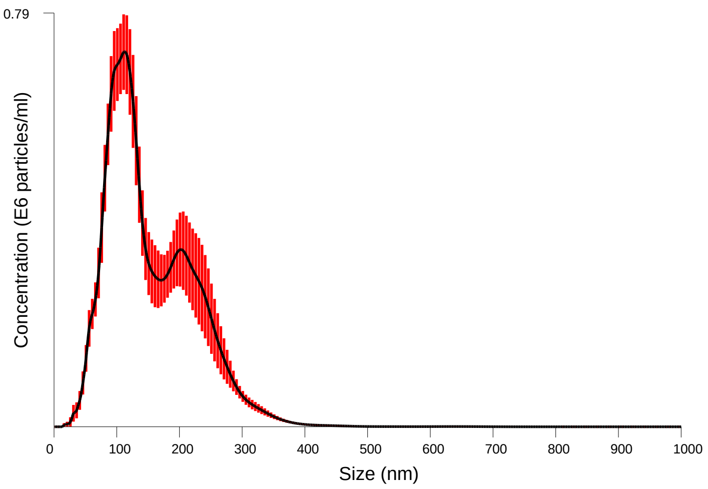


Averaged Size / Concentration
Red error bars indicate +/- 1 standard error of the mean

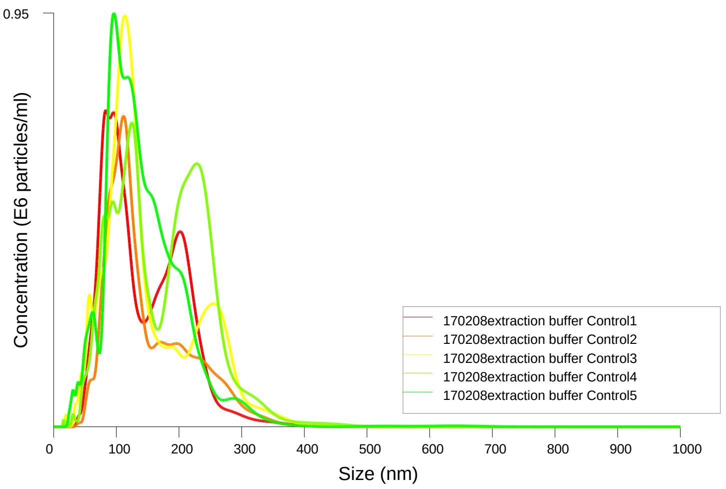


Size / Concentration

| | |
|---|---|
| Included files: | 170208extraction buffer Control1 |
| | 170208extraction buffer Control2 |
| | 170208extraction buffer Control3 |
| | 170208extraction buffer Control4 |
| | 170208extraction buffer Control5 |
| Date/Time of Report | 08/02/2017 14:48:03 |

**BATCH AVERAGE RESULTS:**

| | |
|---|---|
| Size Distribution: | Mean: 153 +/- 5.5 nm |
| | Mode: 105 +/- 7.4 nm |
| | SD: 68 +/- 2.3 nm |
| | D10: 79 +/- 1.6 nm |
| | D50: 137 +/- 9.3 nm |
| | D90: 244 +/- 9.3 nm |
| Total Concentration: | 7.57 +/- 0.62 particles/frame |
| | 0.88 +/- 0.07 E8 particles/ml |
| Total Completed Tracks: | 1783 |
| Average Drift Velocity: | 156 nm/s |

**CAPTURE SETTINGS**

| | |
|---|---|
| Camera Type: | sCMOS |
| Shutter length: | 30.024 ms |
| Shutter setting: | 1200 |
| Camera gain: | 500 |
| Frame rate: | Varied |

**ANALYSIS SETTINGS**

| | |
|---|---|
| Background Extract: | On |
| Detection Threshold: | 10 - Multi |
| Blur: | AUTO |
| Min track length: | AUTO |
| Min expected size: | AUTO |
| Temperature: | 22.0, 22.0, 22.0, 22.0, 22.0 ºC |
| Viscosity: | 0.95, 0.95, 0.95, 0.95, 0.95 cP |

**WARNINGS**

Vibration detected during analysis
