## Supplementary material for "Multi-omic profiling of tyrosine kinase inhibitor-resistant K562 cells suggests metabolic reprogramming to promote cell survival": Exosome supplemental file: 170208media extraction1-001-BATCH-report.pdf

### BATCH ANALYSIS REPORT

Nanoparticle Tracking Analysis (NTA) Version 2.3 Build 0034

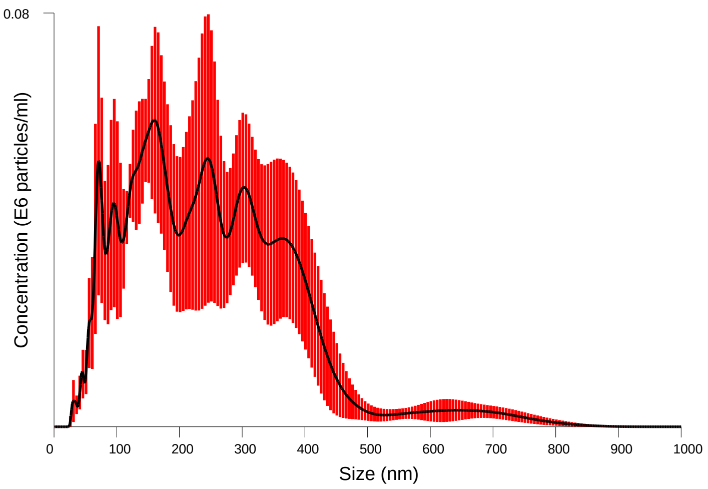


Averaged Size / Concentration
Red error bars indicate +/- 1 standard error of the mean

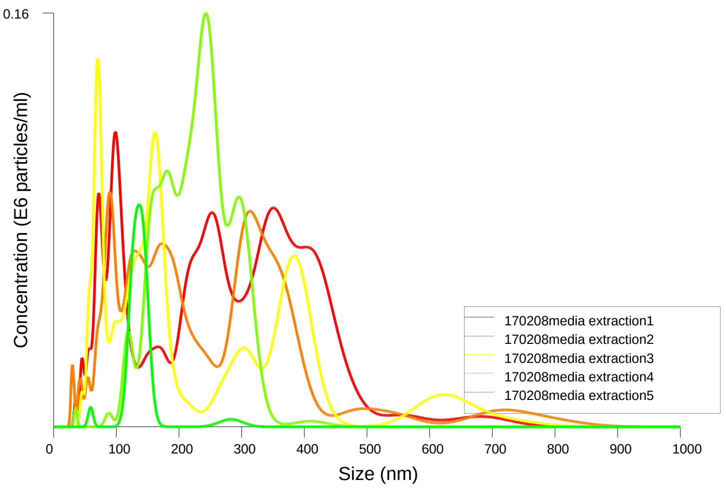


Size / Concentration

| | |
|---|---|
| Included files: | 170208media extraction1 |
| | 170208media extraction2 |
| | 170208media extraction3 |
| | 170208media extraction4 |
| | 170208media extraction5 |
| Date/Time of Report | 08/02/2017 16:00:28 |

**BATCH AVERAGE RESULTS:**

| | |
|---|---|
| Size Distribution: | Mean: 233 +/- 24.7 nm |
| | Mode: 127 +/- 30.8 nm |
| | SD: 110 +/- 27.4 nm |
| | D10: 104 +/- 14.3 nm |
| | D50: 214 +/- 25.7 nm |
| | D90: 347 +/- 54.2 nm |
| Total Concentration: | 1.51 +/- 0.33 particles/frame |
| | 0.18 +/- 0.04 E8 particles/ml |
| Total Completed Tracks: | 265 |
| Average Drift Velocity: | 480 nm/s |

**CAPTURE SETTINGS**

| | |
|---|---|
| Camera Type: | sCMOS |
| Shutter length: | Varied |
| Shutter setting: | Varied |
| Camera gain: | Varied |
| Frame rate: | Varied |

**ANALYSIS SETTINGS**

| | |
|---|---|
| Background Extract: | On |
| Detection Threshold: | 10 - Multi |
| Blur: | AUTO |
| Min track length: | AUTO |
| Min expected size: | AUTO |
| Temperature: | 22.0, 22.0, 22.0, 22.0, 22.0 °C |
| Viscosity: | 0.95, 0.95, 0.95, 0.95, 0.95 cP |

**WARNINGS**

Vibration detected during analysis
Auto max jump failed in some analyses
