## Supplementary material for "Multi-omic profiling of tyrosine kinase inhibitor-resistant K562 cells suggests metabolic reprogramming to promote cell survival": Metabolite profiling data and report: SupplementalReport_(E-160864)MINUV-HMT-002_MetabolomeAnalysisReport.pdf

Research Report

(F-SCOPE)

### Metabolome Profiles of K562 Cell Line by CE-TOFMS Analysis

Client：University of Minnesota

Report Number：MINUV-HMT-002

Report Date：December 18, 2016

Human Metabolome Technologies, Inc.

This report consists of 13 pages (Including title page).

#### 1. Purpose of Study

Analyzing the ionic metabolites including in the K562 cell line by CE-TOFMS.

#### 2. Summary

Metabolome analysis was performed in 6 samples of K562 cell line using CE-TOFMS[†1] in two modes for cationic and anionic metabolites. We detected 126 peaks of isotopic ions and their total amount for each compounds on the basis of HMT’s standard library.

[†1] Capillary Electrophoresis Time-of-Flight Mass Spectrometry

### 3. Material and Methods

#### 3-1 Materials

K562 cell line was sent from University of Minnesota (In this report, it was written as "University of Minnesota".) to HMT. Sample information was shown in Table 1.

##### Table 1 Sample Information

| Name | Amount ($\times 10^6$ cells) | Group |
|---|---|---|
| WT-1 | 7 | |
| WT-2 | 7 | Control |
| WT-3 | 7 | |
| IR-1 | 7 | |
| IR-2 | 7 | Drug |
| IR-3 | 7 | |

#### 3-2 Sample Preparation

The samples were treated in University of Minnesota according to the protocol provided by HMT (Document ID : [E-150782]) and sent to HMT. In HMT, the samples were resuspended in 50 μL of ultrapure water immediately before the measurement.

#### 4. Measurement

The compounds were measured in the Anion mode of CE-TOFMS based metabolome analysis in the following conditions [1-3].

Anionic Metabolites (Anion Mode)

Device

Agilent CE-TOFMS system (Agilent Technologies Inc.) Machine No. 2
Capillary: Fused silica capillary i.d. 50 μm × 80 cm

Analytical Condition

| | |
|---|---|
| Run buffer: | Anion Buffer Solution (p/n : H3302-1021) |
| Rinse buffer: | Anion Buffer Solution (p/n : H3302-1021) |
| Sample injection: | Pressure injection 50 mbar, 25 sec |
| CE voltage: | Positive, 30 kV |
| MS ionization: | ESI Negative |
| MS capillary voltage: | 3,500 V |
| MS scan range: | *m/z* 50-1,000 |
| Sheath liquid: | HMT Sheath Liquid (p/n : H3301-1020) |

### 5. Data Processing and Analysis

#### 5-1 Data Processing

Peaks detected in CE-TOFMS analysis were extracted using automatic integration software (MasterHands ver. 2.17.1.11 developed at Keio University) in order to obtain peak information including *m/z*, migration time (MT), and peak area. The peak area was then converted to relative peak area by the following equation[†2]. The peak detection limit was determined based on signal-noise ratio; S/N = 3.

$$^{\dagger 2}\ \text{Relative Peak Area} = \frac{\text{Metabolite Peak Area}}{\text{Internal Standard Peak Area} \times \text{Sample Amount}}$$

#### 5-2 Annotation of Peaks

Putative metabolites were then assigned from HMT's target library and their isotopic ions on the basis of *m/z* and MT. The tolerance was ±0.5 min in MT and ±30 ppm[†3] in *m/z*.

$$^{\dagger 3}\ \text{Mass error (ppm)} = \frac{\text{Measured Value} - \text{Theoretical Value}}{\text{Measured Value}} \times 10^6$$

#### 5-3 Quantitative Estimation of Target Metabolites

Absolute quantification was performed in total amount of each detected metabolite. All the metabolite concentrations were calculated by normalizing the peak area of each metabolite with respect to the area of the internal standard and by using standard curves, which were obtained by single-point (100 μM) calibrations.

### 6. Results

#### 6-1 Putative Metabolites

From CE-TOFMS measurement of 6 samples, 126 peaks of isotopic ions and their total amount for each compounds were detected and annotated on the basis of HMT's standard library (Table 2). The detail information is listed in the attached Excel file.

#### 6-2 Comparative Analysis between Study Groups

For detected 126 peaks, the results are summarized in Table 2 and the detail information was listed in the attached file.

#### 6-3 Quantitative Estimation of Target Metabolites

Among target metabolites, 27 metabolites were detected and quantified by using total amount of isotopic ions (Table 3).

***Information***

The results including individual peak information were listed in attached file.
(File name: *Project ID*_Metabolome Analysis Data)

***Table 2 "Putative Metabolites"***

- Peak ID consists of analysis mode and number. The alphabets shows measurement mode; Cation (C) and Anion (A) mode.
- The data with "Total" represents total detection values of isotopic ions for each comounds The data is calcurated result of actual peaks therefore have not assigned peak ID.
- Putative metabolites listed in "Compound name" were assigned on the basis of $m/z$ and MT. Those listed in "KEGG ID / HMDB ID / peptide" were assigned on the basis of $m/z$ only.
- "N.D."and "N.A" represent "Not Detected" and "Not Available", respectively. The target peaks which were N.D. for all samples are not listed in the table.
- The detection values of isotopic ions include those of naturally existed isotopes.
- "Ratio" was calculated between two indicated groups (*left*: numerator ,*right*: dominator). "*p*-value" was calculated on the basis of Welch's *t*-test.
- The information about each result was indicated under the table.

### Table 2 Putative Metabolites (1)

| ID | HMT DB † Compound name | Label | Relative Area Control Mean | Control S.D. | Drug Mean | Drug S.D. | Comparative Analysis Drug vs Control Ratio ¶ | *p*-value ‖ | |
|---|---|---|---|---|---|---|---|---|---|
| - | 2-Oxoglutaric acid | Total | 2.5E-02 | 1.6E-03 | 1.6E-02 | 8.6E-04 | 0.6 | 0.003 | ** |
| A_0002 | 2-Oxoglutaric acid | 0 | 1.7E-02 | 1.1E-03 | 1.0E-02 | 3.0E-04 | 0.6 | 0.006 | ** |
| A_0004 | 2-Oxoglutaric acid | 2 | 8.1E-03 | 6.5E-04 | 5.5E-03 | 5.9E-04 | 0.7 | 0.007 | ** |
| - | 2-Phosphoglyceric acid | Total | 5.6E-04 | N.A. | 1.2E-03 | 2.8E-04 | 2.1 | N.A. | |
| A_0008 | 2-Phosphoglyceric acid | 3 | 5.6E-04 | N.A. | 1.2E-03 | 2.8E-04 | 2.1 | N.A. | |
| - | 3-Hydroxybutyric acid | Total | 2.3E-03 | 1.3E-03 | 1.7E-03 | 2.0E-04 | 0.8 | 0.572 | |
| A_0010 | 3-Hydroxybutyric acid | 0 | 2.3E-03 | 1.3E-03 | 1.7E-03 | 2.0E-04 | 0.8 | 0.572 | |
| - | 3-Phosphoglyceric acid | Total | 6.2E-03 | 5.7E-04 | 7.5E-03 | 2.2E-04 | 1.2 | 0.049 | * |
| A_0014 | 3-Phosphoglyceric acid | 0 | 2.7E-03 | 2.2E-04 | 1.5E-03 | 1.5E-04 | 0.6 | 0.003 | ** |
| A_0017 | 3-Phosphoglyceric acid | 3 | 3.5E-03 | 3.6E-04 | 6.0E-03 | 2.8E-04 | 1.7 | 9.9E-04 | *** |
| - | 6-Phosphogluconic acid | Total | N.A. | N.A. | 6.7E-04 | 4.9E-05 | 1< | N.A. | |
| A_0022 | 6-Phosphogluconic acid | 6 | N.A. | N.A. | 6.7E-04 | 4.9E-05 | 1< | N.A. | |
| - | Acetyl CoA_divalent | Total | N.A. | N.A. | 6.4E-04 | N.A. | 1< | N.A. | |
| A_0029 | Acetyl CoA_divalent | 7 | N.A. | N.A. | 6.4E-04 | N.A. | 1< | N.A. | |
| - | ADP | Total | 2.4E-01 | 1.4E-02 | 2.2E-01 | 1.4E-02 | 0.9 | 0.136 | |
| A_0031 | ADP | 0 | 3.7E-02 | 1.7E-03 | 3.6E-02 | 2.0E-03 | 1.0 | 0.652 | |
| A_0032 | ADP | 1 | 6.2E-03 | 4.3E-04 | 5.4E-03 | 2.0E-04 | 0.9 | 0.063 | |
| A_0033 | ADP | 2 | 5.6E-03 | 2.5E-04 | 5.4E-03 | 4.5E-04 | 1.0 | 0.446 | |
| A_0034 | ADP | 3 | 5.1E-03 | 2.7E-04 | 4.9E-03 | 4.2E-04 | 1.0 | 0.554 | |
| A_0035 | ADP | 4 | 9.7E-03 | 6.4E-04 | 9.3E-03 | 3.7E-04 | 1.0 | 0.488 | |
| A_0036 | ADP | 5 | 1.0E-01 | 5.9E-03 | 1.2E-01 | 8.6E-03 | 1.1 | 0.074 | |
| A_0037 | ADP | 6 | 4.5E-02 | 3.6E-03 | 2.7E-02 | 1.1E-03 | 0.6 | 0.008 | ** |
| A_0038 | ADP | 7 | 1.9E-02 | 1.2E-03 | 8.8E-03 | 6.6E-04 | 0.5 | 0.001 | ** |
| A_0039 | ADP | 8 | 6.5E-03 | 6.6E-04 | 1.8E-03 | 2.3E-04 | 0.3 | 0.003 | ** |
| A_0040 | ADP | 9 | 1.2E-03 | 1.4E-04 | N.A. | N.A. | <1 | N.A. | |
| - | AMP | Total | 7.2E-02 | 3.0E-03 | 4.8E-02 | 4.7E-03 | 0.7 | 0.003 | ** |
| A_0042 | AMP | 0 | 1.2E-02 | 3.4E-04 | 9.3E-03 | 5.4E-04 | 0.8 | 0.005 | ** |
| A_0043 | AMP | 1 | 1.8E-03 | 1.5E-04 | 1.4E-03 | 1.5E-05 | 0.8 | 0.066 | |
| A_0044 | AMP | 2 | 1.7E-03 | 7.4E-05 | 1.2E-03 | 3.5E-04 | 0.7 | 0.115 | |
| A_0045 | AMP | 3 | 1.9E-03 | 1.2E-04 | 1.2E-03 | 9.9E-05 | 0.6 | 0.002 | ** |
| A_0046 | AMP | 4 | 3.4E-03 | 8.3E-05 | 2.4E-03 | 2.3E-04 | 0.7 | 0.010 | ** |
| A_0047 | AMP | 5 | 3.0E-02 | 1.8E-03 | 2.5E-02 | 2.3E-03 | 0.8 | 0.031 | * |
| A_0048 | AMP | 6 | 1.3E-02 | 7.5E-04 | 5.5E-03 | 8.2E-04 | 0.4 | 3.6E-04 | *** |
| A_0049 | AMP | 7 | 6.0E-03 | 5.3E-04 | 2.5E-03 | 2.3E-04 | 0.4 | 0.003 | ** |
| A_0050 | AMP | 8 | 1.8E-03 | 6.9E-05 | 5.5E-04 | N.A. | 0.3 | N.A. | |
| A_0051 | AMP | 9 | 4.9E-04 | 5.1E-05 | N.A. | N.A. | <1 | N.A. | |
| - | ATP | Total | 6.5E-01 | 2.0E-02 | 8.5E-01 | 6.4E-03 | 1.3 | 0.001 | ** |
| A_0053 | ATP | 0 | 1.2E-01 | 4.1E-03 | 1.5E-01 | 1.6E-03 | 1.2 | 0.003 | ** |
| A_0054 | ATP | 1 | 1.8E-02 | 6.6E-04 | 2.2E-02 | 7.5E-04 | 1.2 | 0.003 | ** |
| A_0055 | ATP | 2 | 1.7E-02 | 6.0E-04 | 2.1E-02 | 6.8E-04 | 1.2 | 0.002 | ** |
| A_0056 | ATP | 3 | 1.5E-02 | 3.8E-04 | 1.8E-02 | 7.8E-05 | 1.2 | 0.003 | ** |
| A_0057 | ATP | 4 | 2.6E-02 | 1.1E-03 | 3.5E-02 | 6.5E-04 | 1.3 | 6.2E-04 | *** |
| A_0058 | ATP | 5 | 2.8E-01 | 9.2E-03 | 4.7E-01 | 4.5E-03 | 1.7 | 8.7E-05 | *** |
| A_0059 | ATP | 6 | 1.1E-01 | 2.9E-03 | 9.2E-02 | 5.0E-04 | 0.9 | 0.011 | * |
| A_0060 | ATP | 7 | 4.8E-02 | 2.0E-03 | 3.3E-02 | 3.0E-04 | 0.7 | 0.005 | ** |
| A_0061 | ATP | 8 | 1.6E-02 | 9.9E-04 | 7.0E-03 | 1.6E-04 | 0.4 | 0.003 | ** |
| A_0062 | ATP | 9 | 3.0E-03 | 1.4E-05 | 1.1E-03 | 4.2E-05 | 0.4 | 3.9E-05 | *** |
| - | cis-Aconitic acid | Total | 9.6E-03 | 2.3E-03 | 7.3E-03 | 2.4E-03 | 0.8 | 0.301 | |
| A_0064 | cis-Aconitic acid | 0 | 2.2E-03 | 2.1E-04 | 1.9E-03 | 2.1E-04 | 0.9 | 0.291 | |
| A_0066 | cis-Aconitic acid | 2 | 6.6E-03 | 3.2E-04 | 5.2E-03 | 1.2E-04 | 0.8 | 0.010 | ** |
| A_0067 | cis-Aconitic acid | 4 | 2.4E-03 | 7.2E-05 | 2.8E-03 | N.A. | 1.1 | N.A. | |
| - | Citric acid | Total | 2.3E-01 | 4.7E-03 | 2.0E-01 | 8.5E-03 | 0.9 | 0.010 | * |
| A_0069 | Citric acid | 0 | 3.5E-02 | 3.3E-04 | 2.6E-02 | 1.3E-03 | 0.7 | 0.005 | ** |
| A_0070 | Citric acid | 1 | 9.4E-03 | 5.6E-04 | 7.4E-03 | 2.1E-04 | 0.8 | 0.016 | * |
| A_0071 | Citric acid | 2 | 9.1E-02 | 3.6E-03 | 7.6E-02 | 5.7E-03 | 0.8 | 0.023 | * |
| A_0072 | Citric acid | 3 | 2.1E-02 | 1.9E-04 | 1.9E-02 | 8.3E-04 | 0.9 | 0.022 | * |
| A_0073 | Citric acid | 4 | 3.6E-02 | 6.4E-04 | 3.4E-02 | 2.3E-04 | 0.9 | 0.015 | * |
| A_0074 | Citric acid | 5 | 2.6E-02 | 2.2E-04 | 2.5E-02 | 1.0E-03 | 1.0 | 0.184 | |
| A_0075 | Citric acid | 6 | 7.0E-03 | 4.2E-05 | 8.4E-03 | 3.1E-04 | 1.2 | 0.015 | * |
| - | CoA_divalent | Total | 7.5E-03 | 1.8E-04 | 8.0E-03 | 8.7E-04 | 1.1 | 0.384 | |

ID consists of analysis mode and number. 'A' showed anion mode.
N.D. (Not Detected): The target peak or metabolite was below detection limits.
N.A. (Not Available): The calculation was impossible because of insufficience of the data.
† Putative metabolites which were assigned on the basis of *m/z* and MT in HMT standard compound library.
¶ The ratio is of computed by using averaged detection values. The latter was used as denominator.
‖ The p-value is computed by Welch's t-test. (*<0.05, **<0.01, ***<0.001)
The data are sorted by Compound name in ascending order.

#### Table 2 Putative Metabolites (2)

| ID | HMT DB † Compound name | Label | Relative Area Control Mean | Control S.D. | Drug Mean | Drug S.D. | Comparative Analysis Drug vs Control Ratio ¶ | *p*-value ‖ | |
|---|---|---|---|---|---|---|---|---|---|
| A_0077 | CoA_divalent | 0 | 1.9E-03 | 3.0E-04 | 2.4E-03 | 2.3E-04 | 1.2 | 0.111 | |
| A_0082 | CoA_divalent | 5 | 3.1E-03 | 2.5E-04 | 4.0E-03 | 1.7E-04 | 1.3 | 0.009 | ** |
| A_0083 | CoA_divalent | 6 | 1.6E-03 | 1.2E-04 | 1.4E-03 | 2.4E-04 | 0.8 | 0.191 | |
| A_0084 | CoA_divalent | 7 | 8.1E-04 | 1.6E-04 | 8.2E-04 | N.A. | 1.0 | N.A. | |
| - | Dihydroxyacetone phosphate | Total | N.A. | N.A. | 1.0E-03 | 4.4E-04 | 1< | N.A. | |
| A_0088 | Dihydroxyacetone phosphate | 3 | N.A. | N.A. | 1.0E-03 | 4.4E-04 | 1< | N.A. | |
| - | Fructose 1,6-diphosphate | Total | 1.4E-02 | 5.4E-04 | 5.2E-02 | 1.2E-03 | 3.8 | 3.7E-05 | *** |
| A_0092 | Fructose 1,6-diphosphate | 0 | 5.8E-03 | 2.0E-04 | 8.7E-03 | 6.7E-04 | 1.5 | 0.012 | * |
| A_0094 | Fructose 1,6-diphosphate | 2 | N.A. | N.A. | 7.7E-04 | 4.9E-05 | 1< | N.A. | |
| A_0095 | Fructose 1,6-diphosphate | 3 | 1.1E-03 | 8.2E-05 | 5.5E-03 | 3.4E-04 | 5.1 | 0.001 | ** |
| A_0096 | Fructose 1,6-diphosphate | 4 | N.A. | N.A. | 1.1E-03 | 3.6E-05 | 1< | N.A. | |
| A_0097 | Fructose 1,6-diphosphate | 5 | 7.1E-04 | 1.3E-05 | 3.4E-03 | 4.2E-05 | 4.8 | 2.3E-05 | *** |
| A_0098 | Fructose 1,6-diphosphate | 6 | 6.4E-03 | 4.0E-04 | 3.3E-02 | 8.4E-04 | 5.2 | 2.6E-05 | *** |
| - | Fructose 6-phosphate | Total | 1.1E-03 | 1.1E-04 | 3.2E-03 | 3.6E-04 | 3.0 | 0.006 | ** |
| A_0103 | Fructose 6-phosphate | 6 | 1.1E-03 | 1.1E-04 | 3.2E-03 | 3.6E-04 | 3.0 | 0.006 | ** |
| - | Fumaric acid | Total | 4.2E-02 | 2.3E-03 | 3.7E-02 | 1.0E-03 | 0.9 | 0.061 | |
| A_0105 | Fumaric acid | 0 | 2.5E-02 | 1.4E-03 | 2.1E-02 | 5.5E-04 | 0.8 | 0.024 | * |
| A_0106 | Fumaric acid | 1 | 3.4E-03 | 1.9E-04 | 2.9E-03 | 6.3E-05 | 0.9 | 0.047 | * |
| A_0107 | Fumaric acid | 2 | 8.0E-03 | 4.6E-04 | 8.4E-03 | 6.0E-04 | 1.1 | 0.393 | |
| A_0108 | Fumaric acid | 3 | 5.9E-03 | 3.8E-04 | 5.3E-03 | 6.6E-04 | 0.9 | 0.275 | |
| - | Glucose 1-phosphate | Total | 2.4E-02 | 7.1E-04 | 3.1E-02 | 1.2E-03 | 1.3 | 0.002 | ** |
| A_0110 | Glucose 1-phosphate | 0 | 1.1E-02 | 9.7E-04 | 1.0E-02 | 1.0E-03 | 0.9 | 0.362 | |
| A_0111 | Glucose 1-phosphate | 1 | 9.0E-04 | 5.4E-05 | 8.2E-04 | 4.0E-05 | 0.9 | 0.110 | |
| A_0113 | Glucose 1-phosphate | 3 | 6.9E-04 | 6.0E-05 | 1.3E-03 | 2.1E-05 | 1.8 | 0.035 | * |
| A_0114 | Glucose 1-phosphate | 5 | 1.1E-03 | 2.1E-04 | 1.8E-03 | 9.5E-05 | 1.7 | 0.014 | * |
| A_0115 | Glucose 1-phosphate | 6 | 1.1E-02 | 6.7E-04 | 1.7E-02 | 7.7E-05 | 1.6 | 0.003 | ** |
| - | Glucose 6-phosphate | Total | 6.8E-03 | 7.2E-04 | 1.8E-02 | 2.7E-03 | 2.7 | 0.013 | * |
| A_0117 | Glucose 6-phosphate | 0 | 1.3E-03 | 2.9E-04 | 1.5E-03 | 2.3E-04 | 1.1 | 0.568 | |
| A_0119 | Glucose 6-phosphate | 2 | 8.1E-04 | 5.6E-05 | 1.3E-03 | 1.2E-04 | 1.7 | 0.008 | ** |
| A_0120 | Glucose 6-phosphate | 3 | 1.1E-03 | 2.1E-04 | 2.0E-03 | 1.5E-04 | 1.9 | 0.004 | ** |
| A_0121 | Glucose 6-phosphate | 4 | N.A. | N.A. | 9.6E-04 | 2.4E-05 | 1< | N.A. | |
| A_0122 | Glucose 6-phosphate | 5 | 5.2E-04 | 1.4E-04 | 1.5E-03 | 6.3E-05 | 2.8 | 0.041 | * |
| A_0123 | Glucose 6-phosphate | 6 | 3.3E-03 | 2.0E-04 | 1.1E-02 | 2.0E-03 | 3.5 | 0.018 | * |
| - | Isocitric acid | Total | 6.8E-03 | 1.9E-03 | 4.7E-03 | 9.0E-04 | 0.7 | 0.183 | |
| A_0128 | Isocitric acid | 0 | 2.2E-03 | 1.9E-04 | N.A. | N.A. | <1 | N.A. | |
| A_0130 | Isocitric acid | 2 | 4.9E-03 | 3.4E-04 | 4.2E-03 | 1.6E-04 | 0.9 | 0.063 | |
| A_0131 | Isocitric acid | 5 | 1.5E-03 | N.A. | 1.5E-03 | N.A. | 1.0 | N.A. | |
| - | Lactic acid | Total | 1.1E+00 | 7.1E-02 | 2.8E+00 | 1.0E-01 | 2.6 | 4.7E-05 | *** |
| A_0133 | Lactic acid | 0 | 1.8E-01 | 1.1E-02 | 2.6E-01 | 5.1E-03 | 1.5 | 0.002 | ** |
| A_0134 | Lactic acid | 1 | 1.3E-02 | 7.2E-04 | 2.4E-02 | 8.6E-04 | 1.8 | 8.6E-05 | *** |
| A_0135 | Lactic acid | 2 | 3.5E-02 | 2.4E-03 | 8.6E-02 | 3.3E-03 | 2.4 | 5.5E-05 | *** |
| A_0136 | Lactic acid | 3 | 8.6E-01 | 5.7E-02 | 2.4E+00 | 9.4E-02 | 2.8 | 7.9E-05 | *** |
| - | Malic acid | Total | 3.2E-01 | 9.9E-03 | 3.4E-01 | 1.8E-02 | 1.0 | 0.273 | |
| A_0138 | Malic acid | 0 | 1.7E-01 | 6.7E-03 | 1.6E-01 | 1.1E-02 | 1.0 | 0.496 | |
| A_0139 | Malic acid | 1 | 2.5E-02 | 7.6E-04 | 2.6E-02 | 9.5E-04 | 1.0 | 0.226 | |
| A_0140 | Malic acid | 2 | 6.6E-02 | 1.4E-03 | 7.6E-02 | 2.8E-03 | 1.1 | 0.014 | * |
| A_0141 | Malic acid | 3 | 4.5E-02 | 8.2E-04 | 5.1E-02 | 1.7E-03 | 1.1 | 0.015 | * |
| A_0142 | Malic acid | 4 | 1.4E-02 | 8.4E-04 | 1.9E-02 | 1.2E-03 | 1.4 | 0.004 | ** |
| - | Phosphoenolpyruvic acid | Total | 4.7E-03 | 6.1E-04 | 5.3E-03 | 8.9E-04 | 1.1 | 0.394 | |
| A_0144 | Phosphoenolpyruvic acid | 0 | 1.9E-03 | 2.5E-04 | 1.3E-03 | 3.4E-05 | 0.7 | 0.069 | |
| A_0147 | Phosphoenolpyruvic acid | 3 | 2.8E-03 | 3.6E-04 | 4.4E-03 | 1.3E-04 | 1.6 | 0.010 | * |
| - | PRPP | Total | 6.9E-03 | 4.2E-04 | 1.5E-02 | 1.1E-03 | 2.2 | 0.002 | ** |
| A_0152 | PRPP | 3 | 6.7E-04 | 2.4E-05 | 9.9E-04 | 1.4E-04 | 1.5 | 0.054 | |
| A_0153 | PRPP | 4 | 7.6E-04 | 9.4E-05 | 1.2E-03 | 2.6E-05 | 1.5 | 0.013 | * |
| A_0154 | PRPP | 5 | 5.5E-03 | 3.6E-04 | 1.3E-02 | 9.1E-04 | 2.4 | 0.002 | ** |
| - | Pyruvic acid | Total | 2.9E-03 | 2.1E-04 | 1.1E-02 | 3.5E-04 | 3.8 | 2.6E-05 | *** |
| A_0157 | Pyruvic acid | 3 | 2.9E-03 | 2.1E-04 | 1.1E-02 | 3.5E-04 | 3.8 | 2.6E-05 | *** |
| - | Ribulose 5-phosphate | Total | 3.9E-04 | N.A. | 9.8E-04 | 6.6E-05 | 2.5 | N.A. | |
| A_0166 | Ribulose 5-phosphate | 5 | 3.9E-04 | N.A. | 9.8E-04 | 6.6E-05 | 2.5 | N.A. | |
| - | Sedoheptulose 7-phosphate | Total | 2.1E-03 | 2.0E-04 | 4.9E-03 | 8.0E-04 | 2.3 | 0.022 | * |

ID consists of analysis mode and number. 'A' showed anion mode.
N.D. (Not Detected): The target peak or metabolite was below detection limits.
N.A. (Not Available): The calculation was impossible because of insufficience of the data.
† Putative metabolites which were assigned on the basis of *m/z* and MT in HMT standard compound library.
¶ The ratio is of computed by using averaged detection values. The latter was used as denominator.
‖ The p-value is computed by Welch's t-test. (*<0.05, **<0.01, ***<0.001)
The data are sorted by Compound name in ascending order.

### Table 2 Putative Metabolites (3)

| ID | HMT DB † Compound name | Label | Relative Area Control Mean | | Drug Mean | | Comparative Analysis Drug vs Control Ratio ¶ | p-value ‖ | |
|---|---|---|---|---|---|---|---|---|---|
| | | | Mean | S.D. | Mean | S.D. | Ratio ¶ | *p*-value ‖ | |
| A_0168 | Sedoheptulose 7-phosphate | 0 | 5.1E-04 | N.A. | 7.2E-04 | 6.3E-05 | 1.4 | N.A. | |
| A_0171 | Sedoheptulose 7-phosphate | 6 | N.A. | N.A. | 8.0E-04 | N.A. | 1< | N.A. | |
| A_0172 | Sedoheptulose 7-phosphate | 7 | 1.9E-03 | 1.7E-04 | 3.9E-03 | 3.6E-04 | 2.0 | 0.004 | ** |
| - | Succinic acid | Total | 2.8E-02 | 9.5E-04 | 3.6E-02 | 2.2E-03 | 1.3 | 0.018 | * |
| A_0174 | Succinic acid | 0 | 2.5E-02 | 8.6E-04 | 3.1E-02 | 2.1E-03 | 1.2 | 0.024 | * |
| A_0175 | Succinic acid | 1 | 3.8E-03 | 9.4E-05 | 5.0E-03 | 2.5E-04 | 1.3 | 0.007 | ** |

ID consists of analysis mode and number. 'A' showed anion mode.
N.D. (Not Detected): The target peak or metabolite was below detection limits.
N.A. (Not Available): The calculation was impossible because of insufficience of the data.
† Putative metabolites which were assigned on the basis of *m/z* and MT in HMT standard compound library.
¶ The ratio is of computed by using averaged detection values. The latter was used as denominator.
‖ The p-value is computed by Welch's t-test. (*<0.05, **<0.01, ***<0.001)
The data are sorted by Compound name in ascending order.

#### Table 3 Quantitative Estimation of Target Metabolites

| Metabolite | Concentration (pmol/$10^6$ cells) | | | | Comparative Analysis | | |
|---|---|---|---|---|---|---|---|
| | Control | | Drug | | Drug vs Control | | |
| | Mean | S.D. | Mean | S.D. | Ratio ¶ | *p*-value ‖ | |
| 2-Oxoglutaric acid | 375 | 25 | 238 | 13 | 0.6 | 0.003 | ** |
| 2-Phosphoglyceric acid | 12 | N.A. | 25 | 5.9 | 2.1 | N.A. | |
| 3-Hydroxybutyric acid | 60 | 35 | 46 | 5.3 | 0.8 | 0.572 | |
| 3-Phosphoglyceric acid | 143 | 13 | 172 | 5.2 | 1.2 | 0.049 | * |
| 6-Phosphogluconic acid | N.A. | N.A. | 13 | 0.9 | 1< | N.A. | |
| Acetyl CoA_divalent | N.A. | N.A. | 5.1 | N.A. | 1< | N.A. | |
| ADP | 2,966 | 176 | 2,701 | 172 | 0.9 | 0.136 | |
| AMP | 929 | 39 | 624 | 61 | 0.7 | 0.003 | ** |
| ATP | 6,920 | 208 | 9,020 | 68 | 1.3 | 0.001 | ** |
| cis-Aconitic acid | 69 | 17 | 52 | 17 | 0.8 | 0.301 | |
| Citric acid | 2,224 | 46 | 1,916 | 83 | 0.9 | 0.010 | * |
| CoA_divalent | 66 | 1.6 | 71 | 7.6 | 1.1 | 0.384 | |
| Dihydroxyacetone phosphate | N.A. | N.A. | 31 | 14 | 1< | N.A. | |
| Erythrose 4-phosphate | N.A. | N.A. | N.A. | N.A. | N.A. | N.A. | |
| Fructose 1,6-diphosphate | 160 | 6.2 | 601 | 14 | 3.8 | 3.7E-05 | *** |
| Fructose 6-phosphate | 21 | 2.2 | 63 | 7.1 | 3.0 | 0.006 | ** |
| Fumaric acid | 1,130 | 63 | 1,007 | 28 | 0.9 | 0.061 | |
| Glucose 1-phosphate | 432 | 13 | 555 | 21 | 1.3 | 0.002 | ** |
| Glucose 6-phosphate | 127 | 13 | 341 | 50 | 2.7 | 0.013 | * |
| Glyceraldehyde 3-phosphate | N.A. | N.A. | N.A. | N.A. | N.A. | N.A. | |
| Isocitric acid | 59 | 16 | 41 | 7.8 | 0.7 | 0.183 | |
| Lactic acid | 27,042 | 1,788 | 69,392 | 2,551 | 2.6 | 4.7E-05 | *** |
| Malic acid | 3,526 | 109 | 3,698 | 196 | 1.0 | 0.273 | |
| Phosphoenolpyruvic acid | 88 | 11 | 99 | 17 | 1.1 | 0.394 | |
| PRPP | 114 | 7.0 | 252 | 18 | 2.2 | 0.002 | ** |
| Pyruvic acid | 140 | 10 | 527 | 17 | 3.8 | 2.6E-05 | *** |
| Ribose 5-phosphate | N.A. | N.A. | N.A. | N.A. | N.A. | N.A. | |
| Ribulose 5-phosphate | 8.2 | N.A. | 20 | 1.4 | 2.5 | N.A. | |
| Sedoheptulose 7-phosphate | 37 | 3.6 | 86 | 14 | 2.3 | 0.022 | * |
| Succinic acid | 500 | 17 | 627 | 39 | 1.3 | 0.018 | * |

N.D. (Not Detected): The target peak or metabolite was below detection limits.
N.A. (Not Available): The calculation was impossible because of insufficience of the data.
¶ The ratio is of computed by using averaged detection values. The latter was used as denominator.
‖ The p-value is computed by Welch's t-test. (*<0.05, **<0.01, ***<0.001)
The data are sorted by Compound name in ascending order.

#### 7. Reference

1) T. Soga, D. N. Heiger: Amino acid analysis by capillary electrophoresis electrospray ionization mass spectrometry. *Anal.Chem.* **72**: 1236-1241, 2000.

2) T. Soga, Y. Ueno, H. Naraoka, Y. Ohashi, M. Tomita et al.: Simultaneous determination of anionic intermediates for *Bacillus subtilis* metabolic pathways by capillary electrophoresis electrospray ionization mass spectrometry. *Anal.Chem.* **74**: 2233-2239, 2002.

3) T. Soga, Y. Ohashi, Y. Ueno, H. Naraoka, M. Tomita et al.: Quantitative metabolome analysis using capillary electrophoresis mass spectrometry. *J. Proteome Res.* **2**: 488-494, 2003.

#### 8. Storage Place for Data, Documents and Samples

Data and Documents will be stored in the following place for 12 month after the submission of the final report. Samples will be stored for 6 months. If needed, the samples can be returned after the test. However, when the data re- analysis is not limited to this.

##### Storage Place for Documents

Document Depository in Human Metabolome Technologies, Inc.

Contents

・Contact Document
・Analysis Schedule
・Sample Preparation Procedure
・Analysis Condition
・Analysis Result
・Final Report

##### Storage Place for Electronic Files

Data Server in Human Metabolome Technologies, Inc.

Contents

・Analysis Data
・Analysis Result

##### Storage Place for Samples

Deep Freezer in Human Metabolome Technologies, Inc. (below -80℃)

---

(Contact Information)

Human Metabolome Technologies, Inc.
Metabolome Analysis Group
246-2 Mizukami, Kakuganji, Tsuruoka-shi, Yamagata 997-0052
Japan
