## Supplementary Information for "Multi-omic profiling of tyrosine kinase inhibitor-resistant K562 cells suggests metabolic reprogramming to promote cell survival"

Table of Contents

Supplemental Methods	2

Analysis of BcrAbl fusion transcripts for point mutations using IGV Browser	3

Inhibitor target profiles and FKPM plots for selected transcripts	4

Pathway analysis of gene and protein expression changes show altered signaling and response to oxidative stress/hypoxia	5

Evaluation of Axl upregulation and effects.	8

Exosome secretion analysis	11

Other observations:	12

### Supplemental Methods

**Data analysis**

*Gene set enrichment analysis (GSEA).* Gene set enrichment analysis (GSEA, <http://www.broadinstitute.org/gsea> ^(1-2)^ was performed on the normalized gene expression data (FPKMs). These analyses were performed as four separate comparisons: (1) K562 parental samples compared to all of the drug resistant samples, (2) K562 parental samples compared to all of the IR samples, (3) K562 parental samples compared to all of the nilotinib-resistant samples, and (4) K562 parental samples compared to all of the dasatinib-resistant samples. GSEA was used to compare these expression data to previously published lists of leukemia and CML-specific genes of interest and to the Molecular Signatures Database (MSigDB, v5.1) ^(2)^ curated gene sets collections H: hallmark gene sets, C2: curated gene sets, C4: computational gene sets, and C6: oncogenic gene sets.

*SWATH data processing and quantitation.* SWATH data files (.wiff and .wiff.scan) were converted to mzML in centroid mode using the AB SCIEX MS Data Converter tool (available from http://sciex.com/software-downloads-x2110), followed by conversion to mzXML using msconvert. ^(3)^ These files were then subjected to signal extraction with the DIA-Umpire SE module (version 2.0) ^(4)^ to generate .mgf files for database searching and peptide/protein identification. All three DIA-Umpire SE .mgf files (Q1, Q2 and Q3) from each SWATH data file were used in a search with Protein Pilot 5.0 (SCIEX) against the reviewed human SwissProt/UniProtKB database FASTA file (downloaded 06/07/2015) to identify peptides and proteins directly from the SWATH data for use as the spectral library. The spectral library was generated from the PP5 .group file using the PeakView 2.2 SWATHMicroApp (SCIEX), performing manual retention time alignment to endogenously detected peptides common to all samples. The spectral library and .wiff/.wiff.scan files were uploaded to the BaseSpace/OneOmics cloud environment (Illumina/SCIEX) using the CloudConnect Uploader (SCIEX), and the data processed through the OneOmics Protein Expression Workflow (SCIEX), which is based on quantitation and statistical normalization processes described previously. ^(5)^ In order to address keratin contamination (which is extremely common in proteomics datasets), keratin or epidermal-related proteins for which transcripts were not observed by RNAseq were excluded from interpretation by removing them from the spectral library file. The curated library file was then uploaded to the Basespace platform (Illumina) using the CloudConnect Uploader (SCIEX) for use in the above-mentioned OneOmics analyses.

### Analysis of BcrAbl fusion transcripts for point mutations using IGV Browser

To evaluate whether point mutations were likely to be contributing to TKI resistance in this model, we generated a custom chromosome containing the BcrAbl fusion sequence (mapped to the 210 breakpoint), and mapped the RNAseq reads from the Tophat BAM file to that sequence using IGV Browser. Mutations are shown as colored lines within the reads. As seen in the area of the read maps demarcated by two parallel vertical lines for each sample’s reads in Figure S1 below, no RNAseq reads were observed to contain mutations to the key C nucleotide that gives rise to the most common T315I gatekeeper mutation. Only two individual reads with point mutations to other nucleotides in the codon for T315 were observed in the NR cells, and none in the IR or DR cells. This strongly suggests that point mutations were not a factor in the resistance of these cell lines to their respective TKIs.

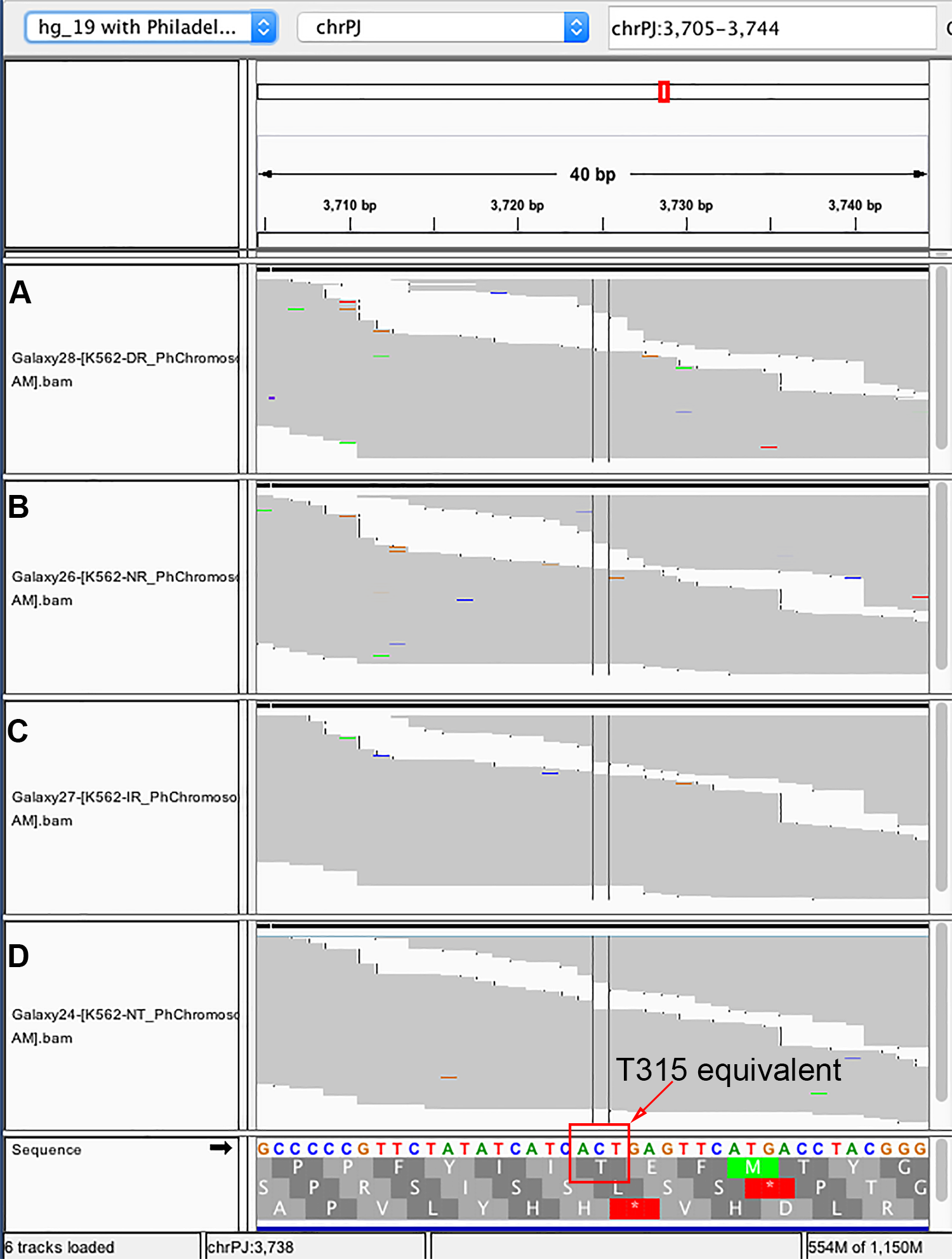

**Figure S1.** RNAseq reads mapped to Ph chromosome show no significant mutations to BcrAbl gatekeeper residue in any of the cell lines. (A) K562-DR, (B) K562-NR, (C) K562-IR, (D) K562-untreated.

### Inhibitor target profiles and FKPM plots for selected transcripts

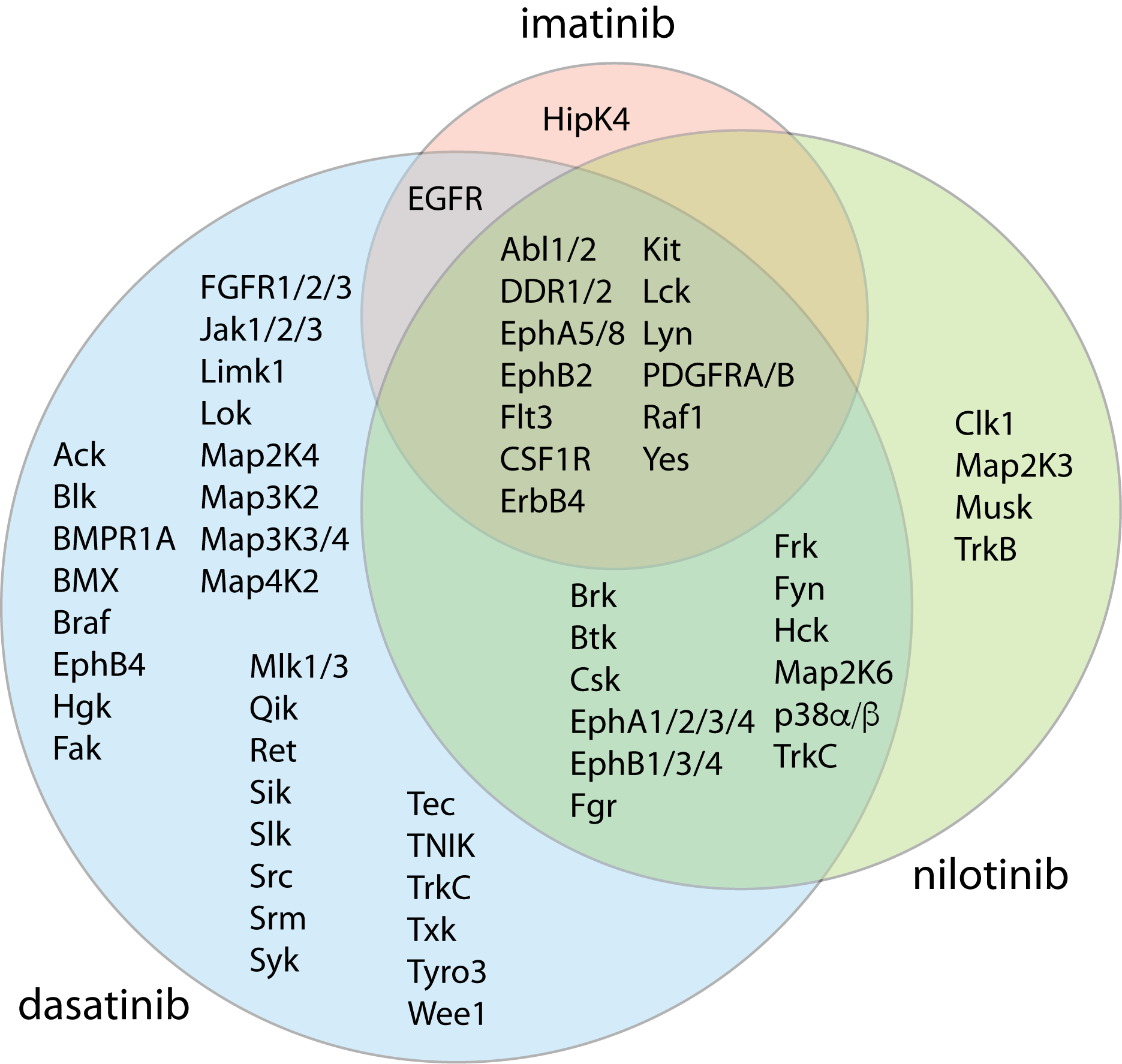

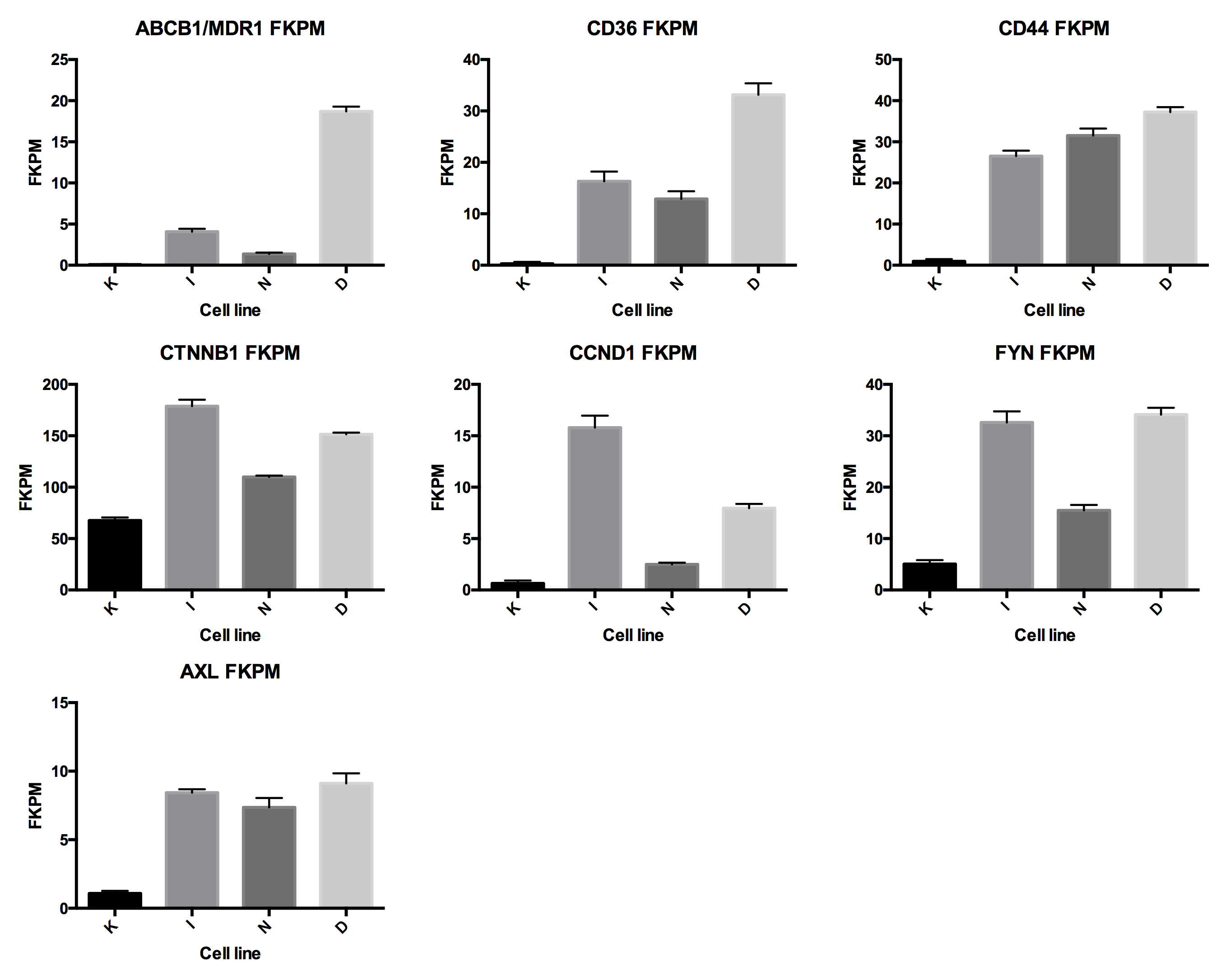

**Figure S2. Top:** Illustration of overlap in known target profiles for imatinib, nilotinib, and dasatinib. A Venn diagram was constructed to visualize data from previously reported *in vitro* kinase assay data published by Kitagawa et al. ^(6)^ that demonstrated targets for imatinib, nilotinib, and dasatinib. These three TKIs inhibited a set of common tyrosine kinases as well as some that are unique to each inhibitor. Kinase data from Kitagawa et al. were included in this diagram if measured IC_50_ was <10,000. Potency against Abl/BcrAbl is ranked dasatinib > nilotinib > imatinib. **Bottom:** RNAseq reads for the untreated (K562), IR (K562-IR), NR (K562-NR) and DR (K562-DR) cells were mapped to the custom genome that included the Ph chromosome-containing BcrAbl fusion gene. FKPM values were calculated (Supplementary Table S2) and plotted for selected transcripts as a visual comparison of expression levels for key genes using GraphPad Prism. Values shown represent the mean and SEM from triplicate RNAseq analyses.

### Gene Set Enrichment Analysis

We also used Gene Set Enrichment Analysis (GSEA) to compare our TKI resistance datasets to the signatures in the Molecular Signatures Database v5.1 (MSigDB, hallmark gene sets, H, curated genes sets, C2, computational gene sets, C4 and oncogenic signatures, C6, http://software.broadinstitute.org/gsea/msigdb/) ^(2)^ (Fig. S3, Supplementary Table S6). This analysis allowed us to screen for similarities between our datasets and the functionally annotated gene expression signatures in the database. In comparing the parental K562s to all TKI resistant samples (as a group and individually), we found that TKI resistant samples were enriched in signatures of hypoxia response (Fig. S3A). Conversely, we found that the parental cells are enriched in signatures of Myc-targets without significant changes in *MYC* transcript levels, which would suggest that TKI-resistant cells may downregulate or bypass Myc-directed gene expression (Fig. S3B). Intriguingly, recent work has implicated the Myc pathway as critical for CML leukemia stem cell survival. ^(7)^ Our gene expression data may imply that some TKI-resistant CML cells could bypass their dependence on Myc. These observations are also consistent with previously described opposing effects of HIF1 vs. Myc on cancer cell metabolism. ^(8)^ This is consistent with the aforementioned reports of metabolic reprogramming as a mechanism for avoiding apoptosis and inducing general drug resistance. Generally, however, hypoxia-related metabolic reprogramming in leukemia is thought to depend on the physiologically hypoxic and stromal environment of bone marrow, ^(9-11)^ or as mentioned above, the adipose niche. ^(12)^ Given that the drug-resistant cell lines described in this work were established and maintained under normoxic conditions in suspension culture, the data shown in this manuscript suggest that these changes may not necessarily be hypoxia-dependent and might not require the bone marrow stromal environment, similar to other previous reports. ^(13-14)^

Initially we considered that Myc protein levels may be decreased in the TKI resistant cells through post-transcriptional or protein degradation mechanisms. However—pseudo-Western blot immunoanalysis using the ProteinSimple Wes “SimpleWestern” system showed that Myc protein levels were essentially the same in all the TKI resistant cell lines as in the parental cells despite some differences at the transcript level (Fig. S3B), suggesting a post-translational mechanism for downregulating Myc targets despite the presence of Myc protein. Since Myc is known to regulate transcriptional programs differently depending on its heterodimerization with other transcription factors, it is also possible that it participates in some alternative heterodimer in the TKI resistant cells that suppresses canonical MYC target activation (e.g. Mad or Mxi). ^(15)^

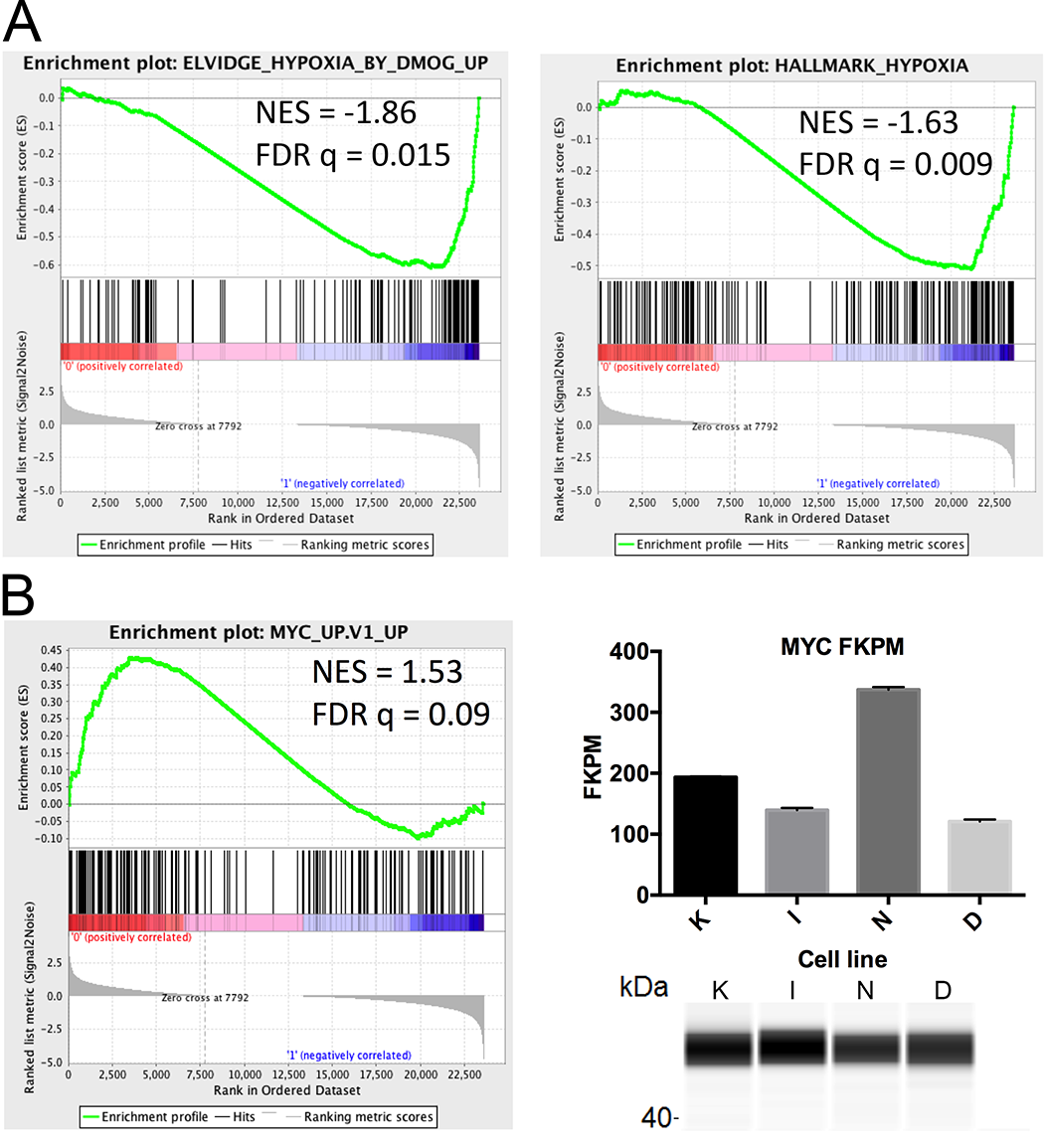

**Figure S3. Gene set enrichment analysis reveals pathways altered with the emergence of treatment resistance.** The gene expression profiles of K562 parental cells and drug resistant cells were compared using gene set enrichment analysis, selected sets illustrated here. (A) Hypoxia gene sets enriched in TKI-resistant cells. (B) Myc gene set enriched in K562 parental cells, but not TKI resistant cells, despite the presence of Myc transcript (top right) and Myc protein (lower right) in the TKI resistant samples.

### Evaluation of Axl upregulation and effects.

We found transcription of the receptor tyrosine kinase Axl to be upregulated by approximately 2-fold in all three cell lines when compared to the parental K562 cell line, validated at the protein level by Western blot (Fig. S4). Axl has been shown to contribute to imatinib and nilotinib resistance in CML, however its role seems to be independent of its kinase activity and it instead operates through a LYN-dependent scaffolding function ^(16-17)^. Therefore, we first chose Axl for further follow up to test the relevance of our models to other previously characterized systems.

***Experimental procedures***

**Growth Inhibition Assay**. *Inhibitor co-treatment experiment.* K562, K562-IR, K562-NR, and K562-DR cells were seeded at 15,000 cells per well in 96-well plates and dosed with the indicated TKI concentrations or vehicle control (0.1% DMSO). For experiments on imatinib, nilotinib and dasatinib, cells were treated with a combination of the particular TKI at the indicated concentrations in the presence of Axl inhibitor R428 (100 nM in DMSO, Med Chem Express #HY-15150) or additional DMSO vehicle. Plates were incubated for three days at 37°C in a 5% CO2 humidified atmosphere. Following incubation, XTT reagent (ATCC) was added according to manufacturer’s protocol and plates were incubated at 37°C in a 5% CO2 humidified atmosphere for 3 hours. Specific absorbance was read on a Biotek Synergy4 plate reader at 475 nm and non-specific absorbance read at 660 nm. Absorbance values at 660 nm were subtracted from 475 nm absorbance values before further data processing. To generate IC50 values, data were plotted in GraphPad Prism and curves were fit using non-linear three-parameter slope, with comparison of fits and IC50 values using the Extra sum of squares F test.

*siRNA knockdown experiment.* K562, K562-IR, K562-NR, and K562-DR cells were transfected with 100nM of SignalSilence^®^ AXL siRNA (Cell Signaling #6263), or SignalSilence^®^ Control siRNA (Cell Signaling #6568) using a Neon Transfection system (Thermo Fisher Scientific) and incubated at 37°C in a 5% CO2 humidified atmosphere. The cells were seeded at 10,000 cells per well in 96-well plates and dosed with the indicated TKI concentrations or vehicle control (0.1% DMSO) 24 hours after transfection. The remaining cells were reserved for Western blot analysis. Plates were incubated for two days at 37°C in a 5% CO2 humidified atmosphere. Following incubation, XTT reagent (ATCC) was added according to manufacturer’s protocol and plates were incubated at 37°C in a 5% CO2 humidified atmosphere for 3 hours. Specific absorbance was read on a Biotek Synergy4 plate reader at 475 nm and non-specific absorbance read at 660 nm. Absorbance values at 660 nm were subtracted from 475 nm absorbance values before further data processing. To generate IC50 values, data were plotted in GraphPad Prism and curves were fit using non-linear three-parameter slope, with comparison of fits and IC50 values using the Extra sum of squares F test.

**Western Blot Analysis.** Cells were treated as described above and pelleted by centrifugation. Media was removed, and cells were washed in ice cold PBS. After washing, lysis buffer (PhosphoSafe Extraction Buffer supplemented with 4 mM EDTA and Complete protease inhibitor (Roche)) was added to the cell pellet and cells were incubated on ice for 30 minutes. Cell pellets were vortexed, then centrifuged at 4° C for 20 minutes to remove insoluble cellular debris and lysate was assayed for protein content. 100 μg of protein was mixed with Laemmli protein gel loading buffer (BioRad) and incubated at 95° C for 5 minutes. Lysate were loaded onto a 10% Mini-Protean TGS gel 59 59 (BioRad) and separated at constant 150 V for 1 hour. Proteins were transferred to a PVDF membrane (Millipore) and blocked with 5% BSA in TBST (tris-buffered saline supplemented supplemented with 0.05% Tween-20). Primary antibodies (AXL (Cell Signaling, #8661) or GAPDH (Sigma Aldrich G8795) diluted 1:1000 in 5% BSA + TBST) were incubated with membrane overnight at 4° C. Following 3 x 5 minute washes with TBST, the membrane was incubated with donkey anti-rabbit tagged with IRDye 800 or donkey anti-mouse tagged with IRDye 680(1:5000 in TBST) for 1 hours at room temperature. After 3 x 5 minute washes with TBST, the membrane was imaged on a LICOR Odyssey infrared scanner.

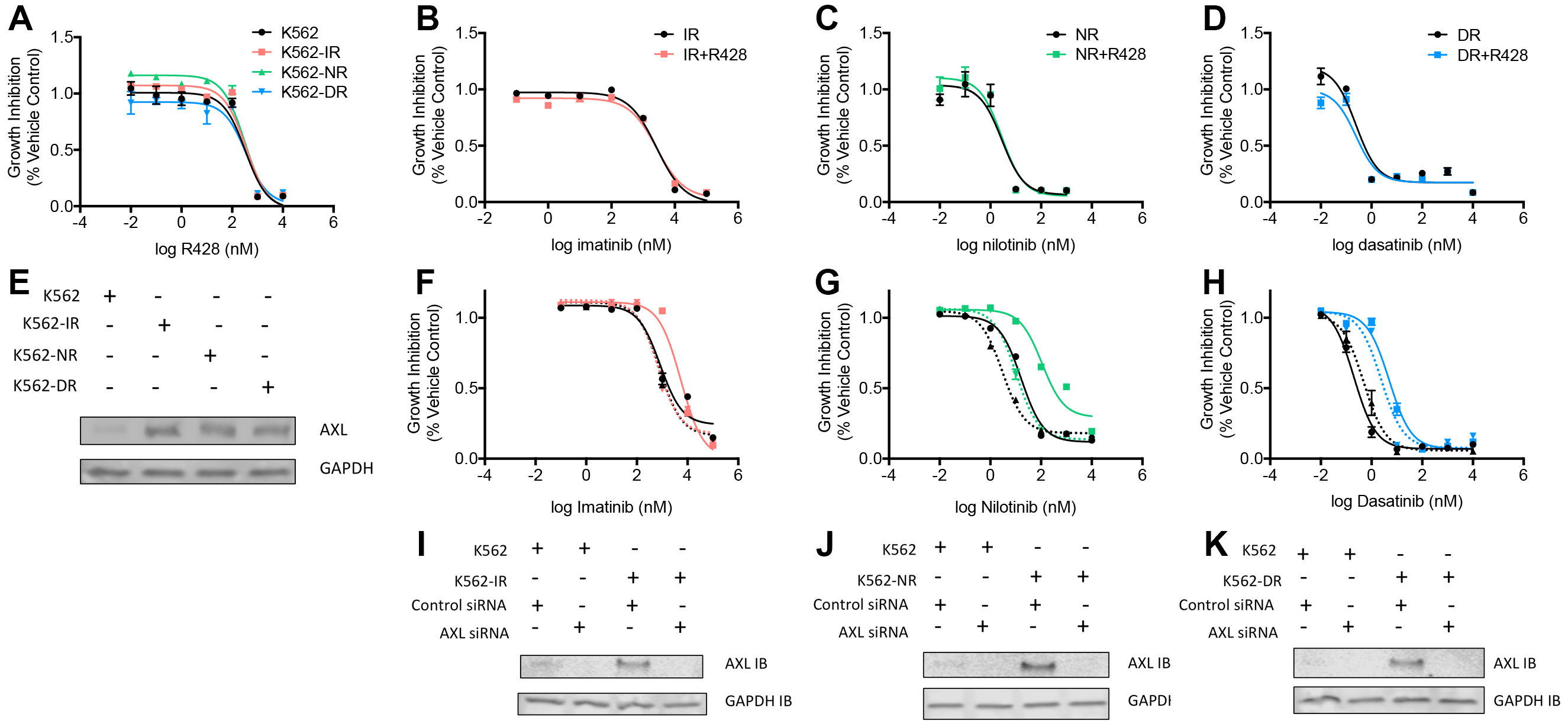

**Figure S4. Testing effect of Axl inhibition and knockdown on TKI resistant cell lines and sensitivity to corresponding TKIs. (**A) R428 (Axl inhibitor) dose-response curves for untreated and IR (orange), NR (green) and DR (blue) cell lines. No significant different was seen in the IC_50_ values for the four cell lines. (B-D) Dose-response curves from co-treatment of IR (B), NR (C) and DR (D) cells with R428 (100 nM) and the TKI to which each had developed resistance. No significant difference was seen in the IC_50_ values for the treatments with and without R428. (E) Western blot showing Axl protein expression levels for the four cell lines. (F-H) Dose-response curves for IR (F, orange), NR (G, green), or DR (H, blue) or untreated control (F, G, H, black) cells treated with the TKI to which each had developed resistance, pre-incubated with control siRNA (solid lines) or AXL knockdown siRNA (dotted lines). (I-K) Western blot controls showing Axl knockdown for each dose-response experiment: IR, NR, and DR, respectively.

**AXL expression is required for nilotinib and imatinib, but not dasatinib resistance**

We also performed some further follow-up on AXL, given the potential for it to play a role in the reprogramming of signaling in these cells and being an alternative target to counteract resistance. ^(18)^ While it was not detected by proteomics (which is common for membrane-bound receptor tyrosine kinases), it was detected as upregulated by immunoblotting (Fig. S4E), thus was increased at both the protein and mRNA level. We examined its possible role in resistance for all three cell lines by inhibiting its kinase activity with axitinib or knocking AXL expression with siRNA. These studies demonstrated that its role was not the same in each cell line (Fig. S4B,C,D,F,G,H). While all the cells were sensitive to axitinib, it did not resensitize to imatinib, nilotinib, or dasatinib. Those data highlight the potential challenges of interpreting the significance of differential gene expression arising from different treatments, even for seemingly identical markers, and supported the relevance of more comprehensive pathway-level evaluation of changes that we described above.

Despite having been implicated in imatinib and nilotinib resistance, Axl has never been implicated in dasatinib resistance. Axl mediates nilotinib resistance through Lyn, ^(16)^ which is a target of dasatinib—therefore we sought to determine if inhibition of Axl kinase activity and/or knockdown of Axl using siRNA in K562-DR cells also resensitized them to dasatinib. We incubated the drug resistant cells with increasing concentrations of imatinib, nilotinib or dasatinib in the presence or absence of the Axl inhibitor axitinib (R428), and also after siRNA knockdown of Axl protein. Inhibitor combination experiments showed no difference in IC_50_ for the inhibitor to which each cell line had become resistant in the presence or absence of axitinib; however, given that both untreated cells (which express lower levels of Axl) and TKI-resistant cells were sensitive to axitinib, we couldn’t rule out the possibility that axitinib’s major effects on these cells are coming from its activity on a different target. The results of the knockdown experiment were more specific for Axl, and confirmed previous reports ^(17)^ that reducing Axl levels in imatinib and nilotinib resistant cells significantly resensitized the cells to those two inhibitors (Figure S4F and G). This indicates that AXL scaffolding is likely an important mediator of acquired imatinib and nilotinib resistance in our model. Interestingly AXL knockdown seemed to further sensitize the WT cells to nilotinib (Figure S4G). On the other hand, resensitization of the K562-DR cell line by Axl knockdown was modest, and sensitivity of K562 and K562-DR cells to dasatinib was largely unaffected by Axl knockdown (Figure S4H), which suggests that AXL does not play a key role in dasatinib resistance of K562-DR cells.

### Exosome secretion analysis

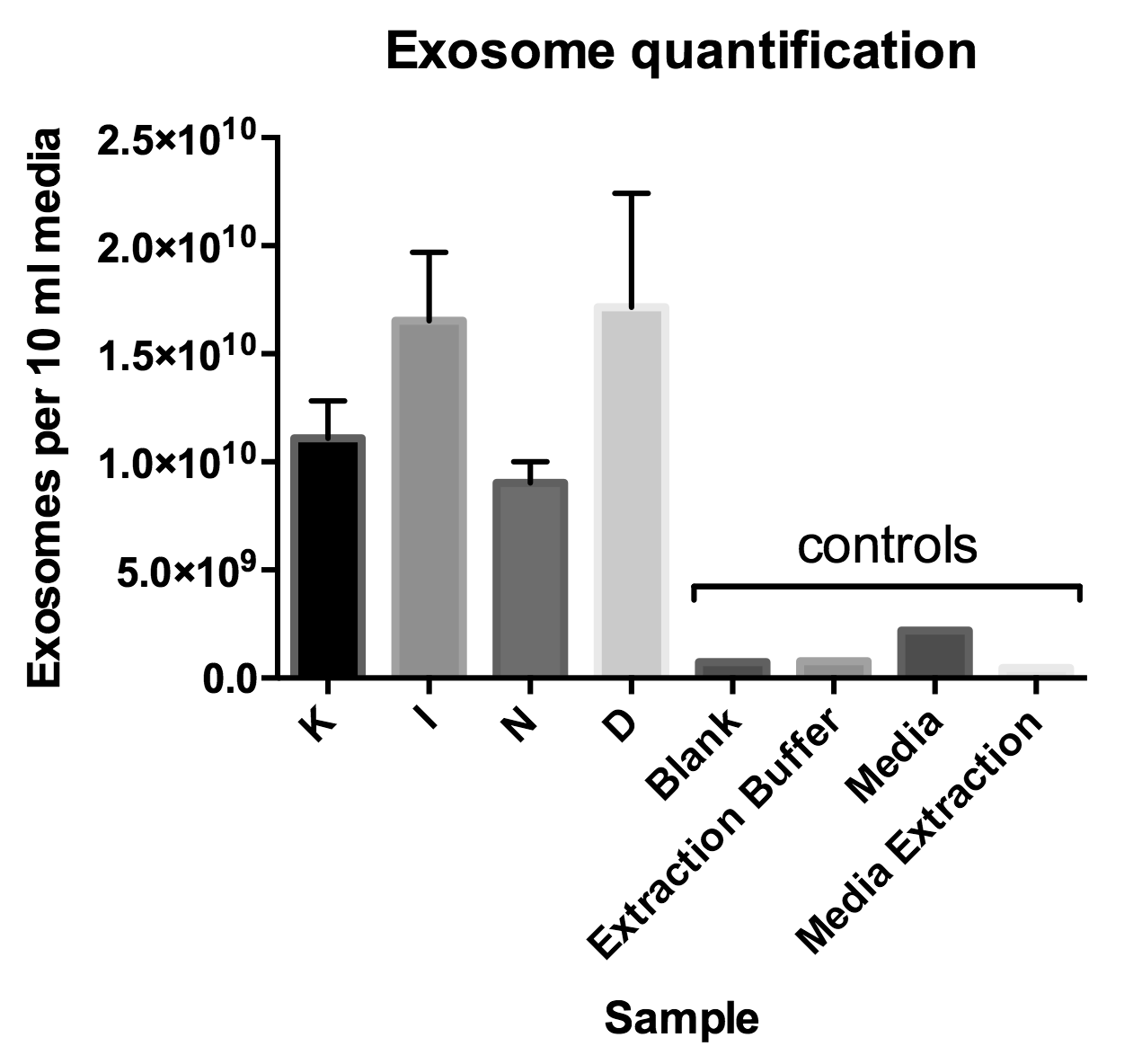

**Figure S5.** Cell lines were cultured to log phase growth (0.8 x 10^6^ cells/ml) in IMDM media prepared with exosome-depleted fetal bovine serum (Gibco/ThermoFisher Scientific #A2720801). Aliquots of cultured media from each cell line were isolated by centrifuging the culture and collecting the supernatant. Culture media (10 ml each) was treated with Total Exosome Isolation Reagent (ThermoFisher Scientific #4478359) according to the manufacturer’s instructions: samples were incubated at 4 ^o^C overnight, collected by centrifugation at 10,000 x g for 60 min, the supernatant discarded and the pellet resuspended in PBS (10 ml) and analyzed on the NanoSight (Malvern) to count nanoparticles by size and determine concentration. Analyses were performed in biological quadruplicate for each cell line (K, I, N and D). Negative controls (PBS buffer blank, mock processing with extraction buffer alone, prepared media diluted into PBS alone, and mock extraction of prepared media without cells were performed (single measurement each) to validate that particle detection was resulting from cell culture.

### Other observations:

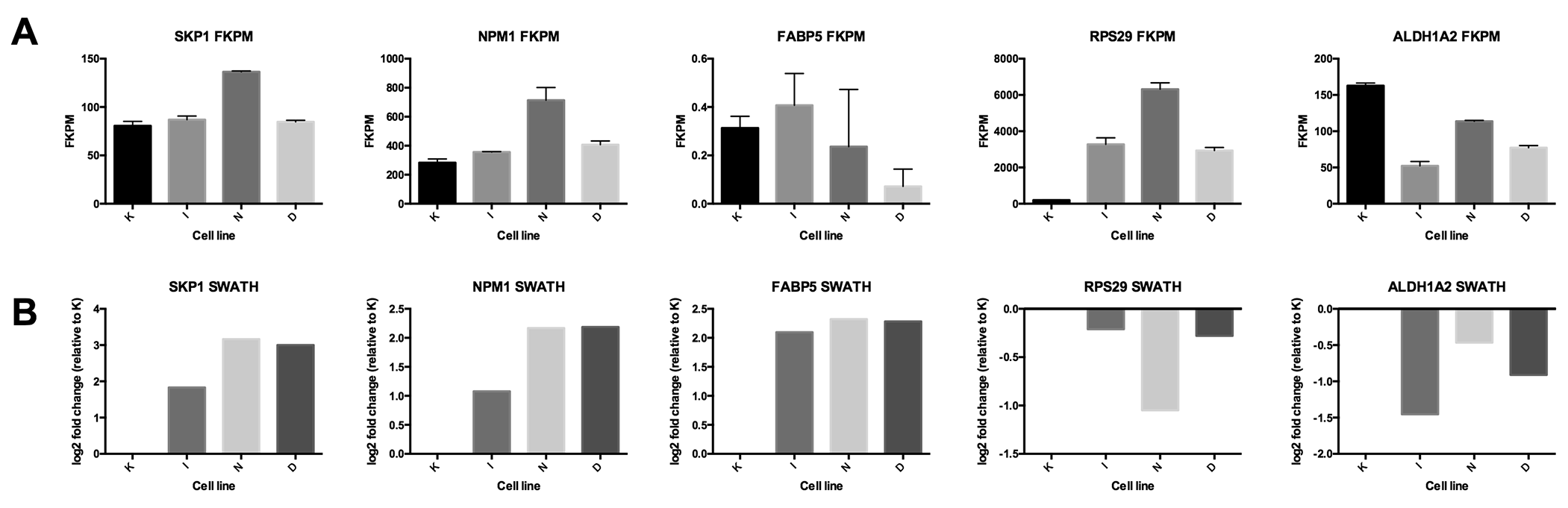

**Figure S6. Additional comparisons between mRNA and protein.** A) FKPM values plotted for selected gene transcripts. Error bars represent SEM for three replicate sequencing runs. K = control K562 cells, I = imatinib resistant, N = nilotinib resistant, D = dasatinib resistant. C) SWATH-MS quantitation values from the OneOmics workflow plotted as log2 fold change for resistant lines relative to K, with unchanged K column shown for reference.

A number of genes seem to be upregulated at the protein level but not the mRNA level or vice versa (Fig. S6), suggesting potential post-transcriptional/translational mechanisms of drug resistance. Notable among these was the E3 ubiquitin ligase Skp1, which, if increased, could go on to affect the ubiquitylation and post-translational degradation of other proteins. Others were NPM1, a protein that is mutated/overexpressed and implicated in other hematological malignancies, and FABP5, a fatty acid transporter protein associated with cell growth and metastatic potential in other cancers ^(19-21)^ and whose family member FABP4 is implicated in CD36-related TKI resistance in a recent study from the Jordan group, ^(12)^ both increased at the protein level but not at the mRNA level. The ribosomal subunit RPS29 was increased at the mRNA level, but not the protein level, suggesting potential translational or post-translational regulation for that transcript/protein.

**Oncoprotein and tumor suppressor levels may be regulated post-transcriptionally or post-translationally in TKI-resistant cells**

One of the key observations of the combined mRNA and protein analysis (Fig. 4A) was that post-transcriptional regulation of protein level seems to be a substantial source of difference between the drug sensitive and resistant cells. Several speculations can be made based on the particular differences detected. For example, we observed seemingly paradoxical behaviors related to MYC signaling. Despite low to modest fold change in MYC transcript and protein levels, Myc-mediated gene expression appears to be significantly downregulated in resistant cells based on GSEA. This apparent downregulation of Myc targets suggests that survival of the resistant cells is either Myc-independent, or represents some alternative regulation of Myc through different binding partners or localization (resulting in e.g. changes in which transcriptional programs Myc participates), which would argue for more detailed examination of Myc’s role in this type of resistance.

The protein nucleophosmin (NPM1) was observed at increased protein levels despite little to no change in mRNA expression. NPM1 plays a role in nuclear/cytoplasmic shuttling and protein stability of the tumor suppressor PTEN. PTEN exerts most of its tumor suppressor function in the nucleus, and is normally regulated by mono-ubiquitylation-dependent shuttling away from nuclear targets to the cytoplasm, where it is deubiquitylated by a protein called HAUSP/USP7 in order to avoid further polyubiquitylation and proteasomal degradation, to maintain PTEN levels for shuttling back upon demand. According to literature models, normally nuclear NPM1 interacts with and inhibits HAUSP/USP7 to prevent nuclear deubiquitination of mono-ubiquitinated PTEN (which would suppress shuttling to the cytoplasm). Thus when NPM1 is overexpressed and aberrantly localized to the cytoplasm, mono-ubiquitinated PTEN is shuttled to the cytoplasm and binds to NPM1 which prevents the protective deubiquitylation, ultimately leading to PTEN polyubiquitylation and degradation. ^(22)^ Interestingly, HAUSP/USP7 is activated by phosphorylation by Bcr-Abl in CML in the cytoplasm, however HAUSP/USP7 inhibitor proteins (such as NPM1) can oppose this activation, and leave PTEN vulnerable to degradation in the cytoplasm. ^(23)^ We observed downregulation of PTEN signaling by Ingenuity Pathway Analysis of the RNAseq data (Fig. 2C), despite a lack of PTEN deregulation at the mRNA level. It is plausible that increased NPM1 protein in TKI resistant cells may disrupt PTEN activity through these types of ubiquitin-mediated mechanisms, however more in-depth characterization would be necessary to confirm this suggestion.

ALDH1A2, a retinal dehydrogenase, was slightly decreased at the mRNA level and at the protein level (Fig. S6)—ALDH1A2 has been proposed as a tumor suppressor in prostate cancer, ^(24)^ so its downregulation being linked to TKI resistance could signify that it has tumor suppressor-like activity in these CML cells as well.

While some of the genes/proteins exhibiting apparent post-transcriptional regulation in common share related functions or have links to known cancer-related proteins, such as Myc and NPM1, there are several seemingly disparate mechanisms represented overall by those features. It is not possible at this point to definitively determine whether these comprise networks of proteins regulated by related upstream mechanisms, however it is reasonable to speculate that these pathways could reveal a role for post-transcriptional and/or post-translational regulation of metabolism in drug resistance that warrants future investigation. One possibility is that the post-transcriptional (e.g. splicing-related) or post-translational regulation of this network of proteins may enable these abnormal hematopoietic progenitor cells to adapt to stress more rapidly than may be required for evolution of targeted point mutation-based resistance. Such adaptation could potentially produce conditions favorable to the evolution of drug-resistant point mutations over time by making populations of cells harboring mutations better able to survive.

**References**

1.	Mootha, V. K.; Lindgren, C. M.; Eriksson, K. F.; Subramanian, A.; Sihag, S.; Lehar, J.; Puigserver, P.; Carlsson, E.; Ridderstrale, M.; Laurila, E.; Houstis, N.; Daly, M. J.; Patterson, N.; Mesirov, J. P.; Golub, T. R.; Tamayo, P.; Spiegelman, B.; Lander, E. S.; Hirschhorn, J. N.; Altshuler, D.; Groop, L. C., PGC-1alpha-responsive genes involved in oxidative phosphorylation are coordinately downregulated in human diabetes. *Nat Genet* **2003,** *34* (3), 267-73.

2.	Subramanian, A.; Tamayo, P.; Mootha, V. K.; Mukherjee, S.; Ebert, B. L.; Gillette, M. A.; Paulovich, A.; Pomeroy, S. L.; Golub, T. R.; Lander, E. S.; Mesirov, J. P., Gene set enrichment analysis: a knowledge-based approach for interpreting genome-wide expression profiles. *Proc Natl Acad Sci U S A* **2005,** *102* (43), 15545-50.

3.	Chambers, M. C.; Maclean, B.; Burke, R.; Amodei, D.; Ruderman, D. L.; Neumann, S.; Gatto, L.; Fischer, B.; Pratt, B.; Egertson, J.; Hoff, K.; Kessner, D.; Tasman, N.; Shulman, N.; Frewen, B.; Baker, T. A.; Brusniak, M. Y.; Paulse, C.; Creasy, D.; Flashner, L.; Kani, K.; Moulding, C.; Seymour, S. L.; Nuwaysir, L. M.; Lefebvre, B.; Kuhlmann, F.; Roark, J.; Rainer, P.; Detlev, S.; Hemenway, T.; Huhmer, A.; Langridge, J.; Connolly, B.; Chadick, T.; Holly, K.; Eckels, J.; Deutsch, E. W.; Moritz, R. L.; Katz, J. E.; Agus, D. B.; MacCoss, M.; Tabb, D. L.; Mallick, P., A cross-platform toolkit for mass spectrometry and proteomics. *Nat Biotechnol* **2012,** *30* (10), 918-20.

4.	Tsou, C. C.; Avtonomov, D.; Larsen, B.; Tucholska, M.; Choi, H.; Gingras, A. C.; Nesvizhskii, A. I., DIA-Umpire: comprehensive computational framework for data-independent acquisition proteomics. *Nat Methods* **2015,** *12* (3), 258-64, 7 p following 264.

5.	Lambert, J. P.; Ivosev, G.; Couzens, A. L.; Larsen, B.; Taipale, M.; Lin, Z. Y.; Zhong, Q.; Lindquist, S.; Vidal, M.; Aebersold, R.; Pawson, T.; Bonner, R.; Tate, S.; Gingras, A. C., Mapping differential interactomes by affinity purification coupled with data-independent mass spectrometry acquisition. *Nat Methods* **2013,** *10* (12), 1239-45.

6.	Kitagawa, D.; Yokota, K.; Gouda, M.; Narumi, Y.; Ohmoto, H.; Nishiwaki, E.; Akita, K.; Kirii, Y., Activity-based kinase profiling of approved tyrosine kinase inhibitors. *Genes Cells* **2013,** *18* (2), 110-22.

7.	Abraham, S. A.; Hopcroft, L. E.; Carrick, E.; Drotar, M. E.; Dunn, K.; Williamson, A. J.; Korfi, K.; Baquero, P.; Park, L. E.; Scott, M. T.; Pellicano, F.; Pierce, A.; Copland, M.; Nourse, C.; Grimmond, S. M.; Vetrie, D.; Whetton, A. D.; Holyoake, T. L., Dual targeting of p53 and c-MYC selectively eliminates leukaemic stem cells. *Nature* **2016,** *534* (7607), 341-6.

8.	Martinez-Outschoorn, U. E.; Peiris-Pages, M.; Pestell, R. G.; Sotgia, F.; Lisanti, M. P., Cancer metabolism: a therapeutic perspective. *Nat Rev Clin Oncol* **2017,** *14* (1), 11-31.

9.	Ng, K. P.; Manjeri, A.; Lee, K. L.; Huang, W.; Tan, S. Y.; Chuah, C. T.; Poellinger, L.; Ong, S. T., Physiologic hypoxia promotes maintenance of CML stem cells despite effective BCR-ABL1 inhibition. *Blood* **2014,** *123* (21), 3316-26.

10.	Kumar, A.; Bhattacharyya, J.; Jaganathan, B. G., Adhesion to stromal cells mediates imatinib resistance in chronic myeloid leukemia through ERK and BMP signaling pathways. *Sci Rep* **2017,** *7* (1), 9535.

11.	Irigoyen, M.; Garcia-Ruiz, J. C.; Berra, E., The hypoxia signalling pathway in haematological malignancies. *Oncotarget* **2017,** *8* (22), 36832-36844.

12.	Ye, H.; Adane, B.; Khan, N.; Sullivan, T.; Minhajuddin, M.; Gasparetto, M.; Stevens, B.; Pei, S.; Balys, M.; Ashton, J. M.; Klemm, D. J.; Woolthuis, C. M.; Stranahan, A. W.; Park, C. Y.; Jordan, C. T., Leukemic Stem Cells Evade Chemotherapy by Metabolic Adaptation to an Adipose Tissue Niche. *Cell Stem Cell* **2016,** *19* (1), 23-37.

13.	Boulahbel, H.; Duran, R. V.; Gottlieb, E., Prolyl hydroxylases as regulators of cell metabolism. *Biochemical Society transactions* **2009,** *37* (Pt 1), 291-4.

14.	Zhao, F.; Mancuso, A.; Bui, T. V.; Tong, X.; Gruber, J. J.; Swider, C. R.; Sanchez, P. V.; Lum, J. J.; Sayed, N.; Melo, J. V.; Perl, A. E.; Carroll, M.; Tuttle, S. W.; Thompson, C. B., Imatinib resistance associated with BCR-ABL upregulation is dependent on HIF-1alpha-induced metabolic reprograming. *Oncogene* **2010,** *29* (20), 2962-72.

15.	Bernards, R., Transcriptional regulation. Flipping the Myc switch. *Current biology : CB* **1995,** *5* (8), 859-61.

16.	Gioia, R.; Leroy, C.; Drullion, C.; Lagarde, V.; Etienne, G.; Dulucq, S.; Lippert, E.; Roche, S.; Mahon, F. X.; Pasquet, J. M., Quantitative phosphoproteomics revealed interplay between Syk and Lyn in the resistance to nilotinib in chronic myeloid leukemia cells. *Blood* **2011,** *118* (8), 2211-21.

17.	Dufies, M.; Jacquel, A.; Belhacene, N.; Robert, G.; Cluzeau, T.; Luciano, F.; Cassuto, J. P.; Raynaud, S.; Auberger, P., Mechanisms of AXL overexpression and function in Imatinib-resistant chronic myeloid leukemia cells. *Oncotarget* **2011,** *2* (11), 874-85.

18.	Ben-Batalla, I.; Erdmann, R.; Jorgensen, H.; Mitchell, R.; Ernst, T.; von Amsberg, G.; Schafhausen, P.; Velthaus, J. L.; Rankin, S.; Clark, R. E.; Koschmieder, S.; Schultze, A.; Mitra, S.; Vandenberghe, P.; Brummendorf, T. H.; Carmeliet, P.; Hochhaus, A.; Pantel, K.; Bokemeyer, C.; Helgason, G. V.; Holyoake, T. L.; Loges, S., Axl Blockade by BGB324 Inhibits BCR-ABL Tyrosine Kinase Inhibitor-Sensitive and -Resistant Chronic Myeloid Leukemia. *Clin Cancer Res* **2017,** *23* (9), 2289-2300.

19.	Powell, C. A.; Nasser, M. W.; Zhao, H.; Wochna, J. C.; Zhang, X.; Shapiro, C.; Shilo, K.; Ganju, R. K., Fatty acid binding protein 5 promotes metastatic potential of triple negative breast cancer cells through enhancing epidermal growth factor receptor stability. *Oncotarget* **2015,** *6* (8), 6373-85.

20.	Forootan, F. S.; Forootan, S. S.; Gou, X.; Yang, J.; Liu, B.; Chen, D.; Al Fayi, M. S.; Al-Jameel, W.; Rudland, P. S.; Hussain, S. A.; Ke, Y., Fatty acid activated PPARgamma promotes tumorigenicity of prostate cancer cells by up regulating VEGF via PPAR responsive elements of the promoter. *Oncotarget* **2016,** *7* (8), 9322-39.

21.	Wang, W.; Chu, H. J.; Liang, Y. C.; Huang, J. M.; Shang, C. L.; Tan, H.; Liu, D.; Zhao, Y. H.; Liu, T. Y.; Yao, S. Z., FABP5 correlates with poor prognosis and promotes tumor cell growth and metastasis in cervical cancer. *Tumour Biol* **2016,** *37* (11), 14873-14883.

22.	Noguera, N. I.; Song, M. S.; Divona, M.; Catalano, G.; Calvo, K. L.; Garcia, F.; Ottone, T.; Florenzano, F.; Faraoni, I.; Battistini, L.; Colombo, E.; Amadori, S.; Pandolfi, P. P.; Lo-Coco, F., Nucleophosmin/B26 regulates PTEN through interaction with HAUSP in acute myeloid leukemia. *Leukemia* **2013,** *27* (5), 1037-43.

23.	Morotti, A.; Panuzzo, C.; Crivellaro, S.; Pergolizzi, B.; Familiari, U.; Berger, A. H.; Saglio, G.; Pandolfi, P. P., BCR-ABL disrupts PTEN nuclear-cytoplasmic shuttling through phosphorylation-dependent activation of HAUSP. *Leukemia* **2014,** *28* (6), 1326-33.

24.	Kim, H.; Lapointe, J.; Kaygusuz, G.; Ong, D. E.; Li, C.; van de Rijn, M.; Brooks, J. D.; Pollack, J. R., The retinoic acid synthesis gene ALDH1a2 is a candidate tumor suppressor in prostate cancer. *Cancer Res* **2005,** *65* (18), 8118-24.
